# Membrane transporter Progressive Ankylosis Protein Homolog (*ANKH*/*Ank*) partially mediates senescence-derived extracellular citrate and is regulated by DNA damage, inflammation and ageing

**DOI:** 10.1101/2024.08.30.609895

**Authors:** Emma Naomi James, Muy-Teck Teh, Yufeng Li, Christine Wagner-Bock, Zahra Falah Al-Khateeb, Lee Peng Karen-Ng, Terry Roberts, Linnea Synchyshyn, Amy Lewis, Ana O’Loghlen, Andrew Silver, Adina Teodora Michael-Titus, Mark Bennett, Jacob Guy Bundy, Maria Elzbieta Mycielska, Eric Kenneth Parkinson

## Abstract

A considerable body of recent evidence supports citrate transport as a major regulator of organismal lifespan and healthspan. Citrate accumulates outside senescent cells *in vitro* and *in vivo*. However, the detailed mechanism of senescent cell extracellular citrate (EC) accumulation is not clear. We show here that EC is partially mediated by a newly described plasma membrane citrate transporter *ANKH/SLC62A1* (progressive human ankylosis - *ANKH*) in senescent fibroblasts. Analogous to interleukin 6 (IL-6), EC and/or *ANKH* are regulated by telomere dysfunction, the p38 mitogen-activated kinase axis, transforming growth factor beta and p53, but in contrast not by steroids, sodium butyrate, or Ataxia Telangiectasia Mutated (ATM). *ANKH* was upregulated in other senescent cell types relevant to ageing but not keratinocytes. In contrast, EC and *ANKH* were inhibited in dividing and senescent fibroblasts by interleukin 1α (IL-1α) in parallel with increased IL-6 secretion. Loss- and gain of function mutations of *ANKH/Ank* are associated with disease and interestingly, *Ank* is also downregulated in both aged mouse liver and brain tissues in parallel with increased senescence markers and several cytokines, suggesting that inflammatory cytokines could inhibit EC production *in vivo*. These data identify *ANKH*/*Ank* as a novel regulator of senescence-derived EC in both humans and mice.

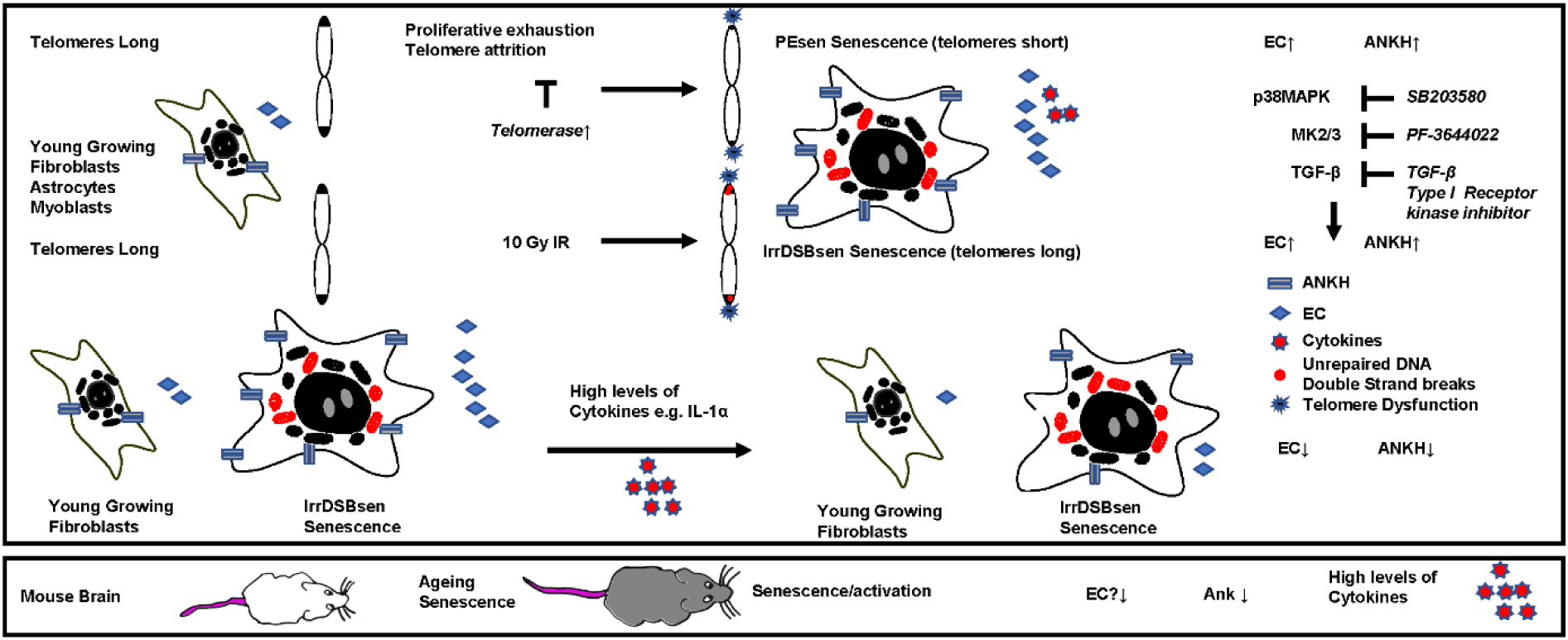

## Introduction

Senescent cells are important to the progression of a wide variety of age-related conditions including Alzheimer’s disease, Parkinson’s disease, atherosclerosis, osteoporosis, osteoarthritis, frailty, type 2 diabetes, memory loss, hepatic steatosis and cancer ^1^. The deletion of senescent cells in mice slows ageing and ameliorates age-related conditions. However, the understanding of the mechanisms by which senescent cells contribute to these diseases, especially in human subjects, remains incomplete.

The tricarboxylic acid cycle metabolite citrate, can transfer between the cytoplasm and the extracellular space in both directions ^2,3^ and accumulates in the body fluids of both ageing humans ^4^ (reviewed in ^5^) and mice ^6,7^, including cerebrospinal fluid ^4^ and urine ^6^. A considerable body of recent evidence now supports citrate uptake and export as a major regulators of organismal lifespan and health span ^8–14^ including development of cancer ^15,16^. However, whether citrate has a detrimental or protective effects depends on factors such as diet, tissue and species ^2,8–12,14^ and is normally tightly regulated ^17^. Therefore, it is currently unclear whether citrate is beneficial or detrimental to health and in which context.

Our previous work has shown that extracellular citrate (EC) is upregulated following the proliferative exhaustion (PEsen) or irreparable DNA double strand break (IrrDSBsen)- induced fibroblast senescence ^18^ and is regulated by telomerase *in vitro* and in the human disease Dyskeratosis Congenita *in vivo* ^19^. A possible explanation for this increase in EC is that senescent fibroblasts ^18^ and melanocytes ^20^, but not breast epithelial cells ^21^, shift their energy metabolism towards glycolysis, as intracellular citrate inhibits phosphofructokinases 1 and 2 and hence glycolysis ^22^. Interestingly, inhibiting glycolysis may slow senescence *in vitro*^20,23^ and improve diabetic wound repair in mice ^23^. Consequently, citrate export and glycolysis may represent ways of regulating senescent cell production and ageing, but the mechanism of EC production by senescent cells and its importance for healthy aging remain unclear.

Recently, a plasma membrane transporter Progressive Ankylosis Protein Homolog *ANKH/SLC62A1* (*ANKH*) previously known to transport pyrophosphate was shown to export citrate and to a lesser extent malate and succinate ^3^. However, our previous work found that senescent fibroblasts do not show elevated levels of inorganic phosphate or pyrophosphate^18^. *ANKH*/*Ank* loss-of-function mutations lead to ankylosis due to the retention of pyrophosphate in the bone ^24^ and citrate accumulation in vascular smooth muscle cells leads to aortic aneurism ^13^. Furthermore, *ANKH* mutations have also been linked to chondrocalcinosis and calcification of the arteries and arthritis ^25^. *Ank* knockout mice ^3^ and humans with loss of *ANKH* function ^26^ have lower levels of plasma and urine citrate and forced expression of *ANKH in vitro* increases extracellular citrate ^3^. In addition, recent extensive genome-wide association screens revealed an association of the *ANKH* gene with a risk of developing Alzheimer’s disease (AD) ^27,28^, other forms of dementia ^29^ and type II diabetes ^30^. Overall, ANKH appeared to be a good candidate for a role in citrate export in senescent cells, but little was known of its connection with senescence or its regulation by known senescence regulatory pathways.

We report that *ANKH* and EC are upregulated concomitantly in a variety of senescent cells and that both were regulated in by some of the molecular pathways that regulate the senescence associated secretory phenotype (SASP) proteins, but not all. Our results identify a potentially important role for *ANKH*/*Ank* in the regulation of EC in many types of senescent cell and by inference, age-related diseases.

## Results

### Identification of ANKH as a candidate citrate exporter in human cellular senescence

Recently, ANKH was identified as a plasma membrane exporter of citrate and to a lesser extent malate ^3^, two metabolites we found accumulated in the conditioned medium of both PEsen and IrrDSBsen senescent fibroblasts ^18^. Hence, we screened 3 fibroblast lines (NHOF-1, BJ and IMR90) for the expression of several plasma membrane transporters including ANKH in both confluent young (cell cycle arrested) and IrrDSBsen fibroblasts. The mitochondria/cytoplasm citrate exporter SLC25A1 (CiC), but not its plasma membrane form (pmCiC), was consistently upregulated in IrrDSBsen cells (Supplementary Figure 1). The same cells also showed an increase of *ANKH* expression in all three IrrDSBsen cultures (Figure 1A); an increase of between 8- and 9.6-fold relative to the confluent young quiescent controls (Figures 1B and 1C). Other plasma membrane transporters, the monocarboxylate transporters MCT1 (Supplementary Figure 1) and MCT4 (Supplementary Figure 1) showed either no upregulation (MCT1) or undetectable expression (MCT4); the neutral amino acid transporter ASCT2 was downregulated in all three IrrDSBsen lines (Supplementary Figure 1). All the full-length blots are shown in Supplementary Figure 2). We next examined whether *ANKH* expression was upregulated in senescent fibroblasts at the mRNA level. In IrrDSBsen NHOF-1 cells this was the case, as assessed by two *ANKH* primer sets (Figure 1D and 1E). P16^INK4A^ was not generally elevated in NHOF-1 cells upon senescence (Figure 1F), but another senescence effector p21^WAF^showed a strong trend for elevation in parallel to *ANKH* (Figure 1G).

**Figure 1.**
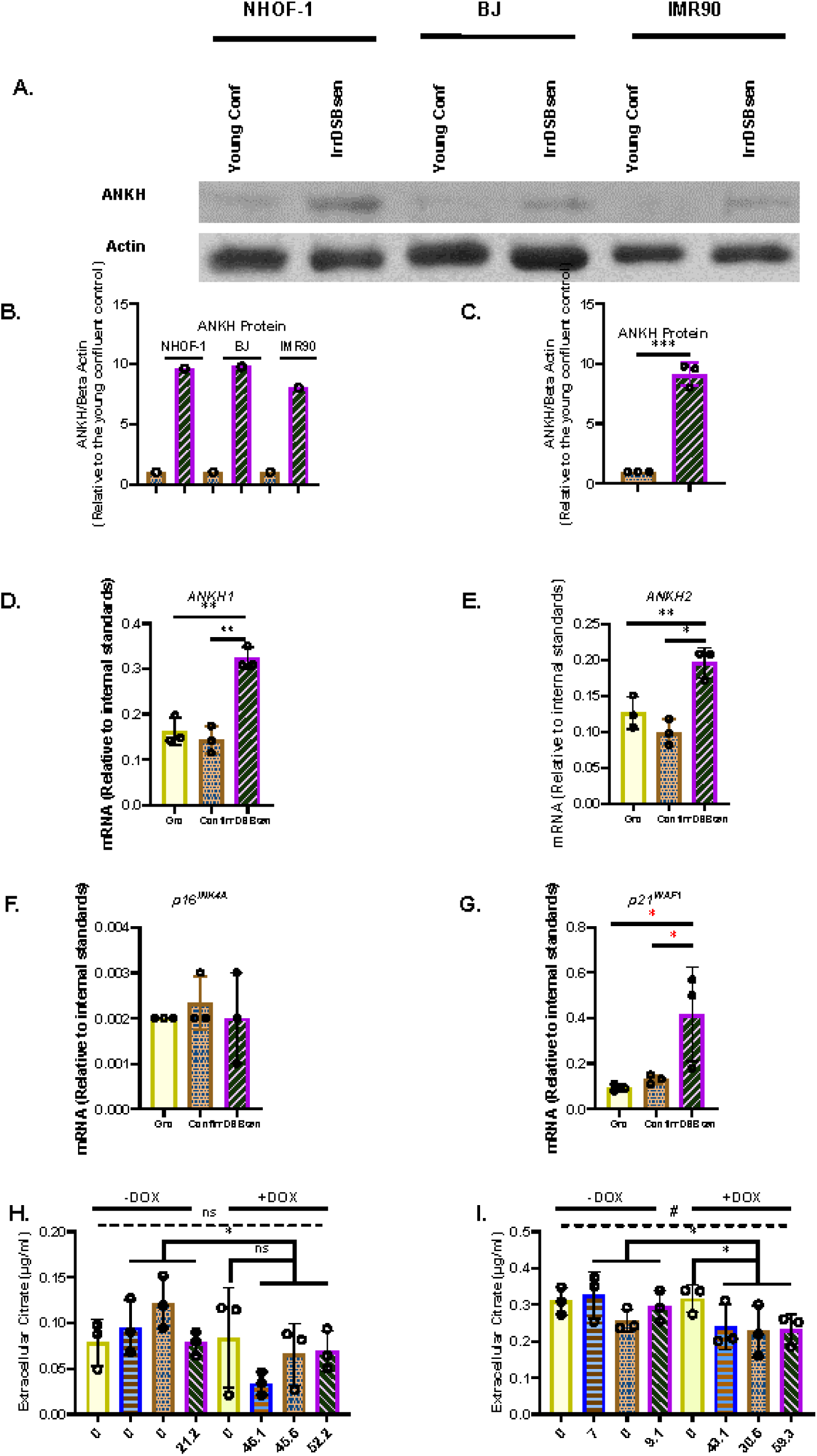
*ANKH* protein and transcript are upregulated in IrrDSBsen human fibroblasts. A. Shows a western blot of *ANKH* protein in 3 lines of quiescent confluent (Conf) young human fibroblasts and the same lines induced to senesce by ionising radiation (IrrDSBsen). Beta action was used as a loading control. B. Image J quantitation of the blot in A. showing the levels of *ANKH* signal versus beta actin in all conditions after background subtraction. The full-length blots are shown in supplementary figure 2. C. The average of the values in B. D. The expression of *ANKH* mRNA in growing confluent and IrrDSBsen NHOF-1 cells using primer set 1. E. As in D. but the results of primer set 2 in the same experiments. F. As in D. but the expression of the senescence marker p16^INK4A^in the same experiments. G. As in D. but the expression of the senescence marker p21^WAF1^ in the same experiments. Yellow plain bars = growing cells; orange stippled bars = confluent cells; purple hatched bars = IrrDSBsen cells. H. The effect of inducible *ANKH* shRNA vectors and their non-targeting (NT) controls with and without the doxycyclin inducer (DOX) on growing NHOF-1 cells. The numbers below each bar represent the level of *ANKH* mRNA knockdown. Yellow plain bars = NT control; blue horizontal striped bars = shRNA #1002; orange stippled bars = shRNA #2500; purple left-right diagonally striped bars = shRNA #5474. I. The effect of inducible *ANKH* shRNA vectors and their non-targeting (NT) controls with and without the doxycyclin inducer (DOX) on IrrDSBsen NHOF-1 cells. The numbers below each bar represent the level of *ANKH* mRNA knockdown. The symbols are the same as for H. (see also supplementary Figure 3). In all experiments * = P< 0.05; ** = P < 0.01; *** = P < 0.001; * P > 0.05; < 0.1; ns = not significant. Data in H. and I. were also analysed by one-way ANOVA (dashed line) # = P< 0.05; ns = not significant. The results are averages +/- standard deviation. N = 3. The experiments in A.- C, D.- G and H-I. were performed at different times but using identical cell culture protocols and reagents.

### The conditional knockdown of *ANKH* mRNA reduces EC in IrrDSBsen human fibroblasts

To evaluate whether *ANKH* was a mediator of EC in senescent fibroblasts we transduced NHOF-1 fibroblasts with 3 commercially validated inducible shRNA vectors targeted against *ANKH* and a non-targeting (NT) control. We then measured the extent of *ANKH* mRNA knockdown and the effect on EC. The results confirmed our previously published data that EC was 4-fold higher in the IrrDSBsen fibroblast conditioned medium than in the growing controls. Inducible shRNA knockdown of *ANKH* inhibited *ANKH* mRNA expression from between 46% and 52% in growing NHOF-1 cells following induction by doxycyclin (DOX; P = 0.001 to P = 0.02 – Supplementary Figure 3) but did not reduce EC levels significantly compared to the NT vector following induction by DOX (P =0.47 by the Welch’s T test; Figure 1H), due to the very low levels of EC in the growing control group and a very low outlier in the NT group. However, the difference between the DOX-induced and control shRNA groups was highly significant (P = 0.005). The same vectors reduced *ANKH* mRNA expression from between 31% and 59% in IrrDSBsen in NHOF-1 cells (Supplementary Figure 3A.B.E.F.) and EC was reduced by between 24 and 28% by all three shRNA vectors relative to the NT vector following induction by DOX (P = 0.03) and was reduced by 12 % (vector #2500) and 27% relative to the non-induced controls (P = 0.025 – Figure 1I). There was no significant effect of DOX on either *ANKH* mRNA or EC in the NT controls, EC in the medium blanks or the cell counts in any of the experimental groups. There was also no significant reduction of p16^INK4A^ (Supplementary Figure 3C and 3G) and p21^WAF^ (Supplementary Figure 3D and 3H) mRNA levels or SA-βGal relative to the NT controls by of the shRNA vectors. However, vectors #1002 and #5474 did show a very small significant induction of p16^INK4A^ in growing cells only. Although the effect of shRNA knockdown on EC reduction was small it was significant in IrrDSBsen cells and given the modest level of *ANKH* mRNA knockdown the data indicate that *ANKH* at least partially mediates EC accumulation following fibroblast senescence.

### *ANKH* mRNA is upregulated in parallel with senescence markers in PEsen human fibroblasts and is downregulated by the canonical function of telomerase

Next, we tested whether *ANKH* was upregulated in PEsen human fibroblasts in parallel with senescence markers and showed that this is indeed the case in NHOF-1 cells (Figure 2A and 2B); p16^INK4A^ (Figure 2C), p21^WAF1^ (Figure 2D) and SA-βGal (Figure 2E) are strongly upregulated in the same cells. Comparable results were obtained with PEsen BJ cells (Supplementary Figure 4A-E), except that p16^INK4A^ was not upregulated ((Supplementary Figure 4C).

**Figure 2.**
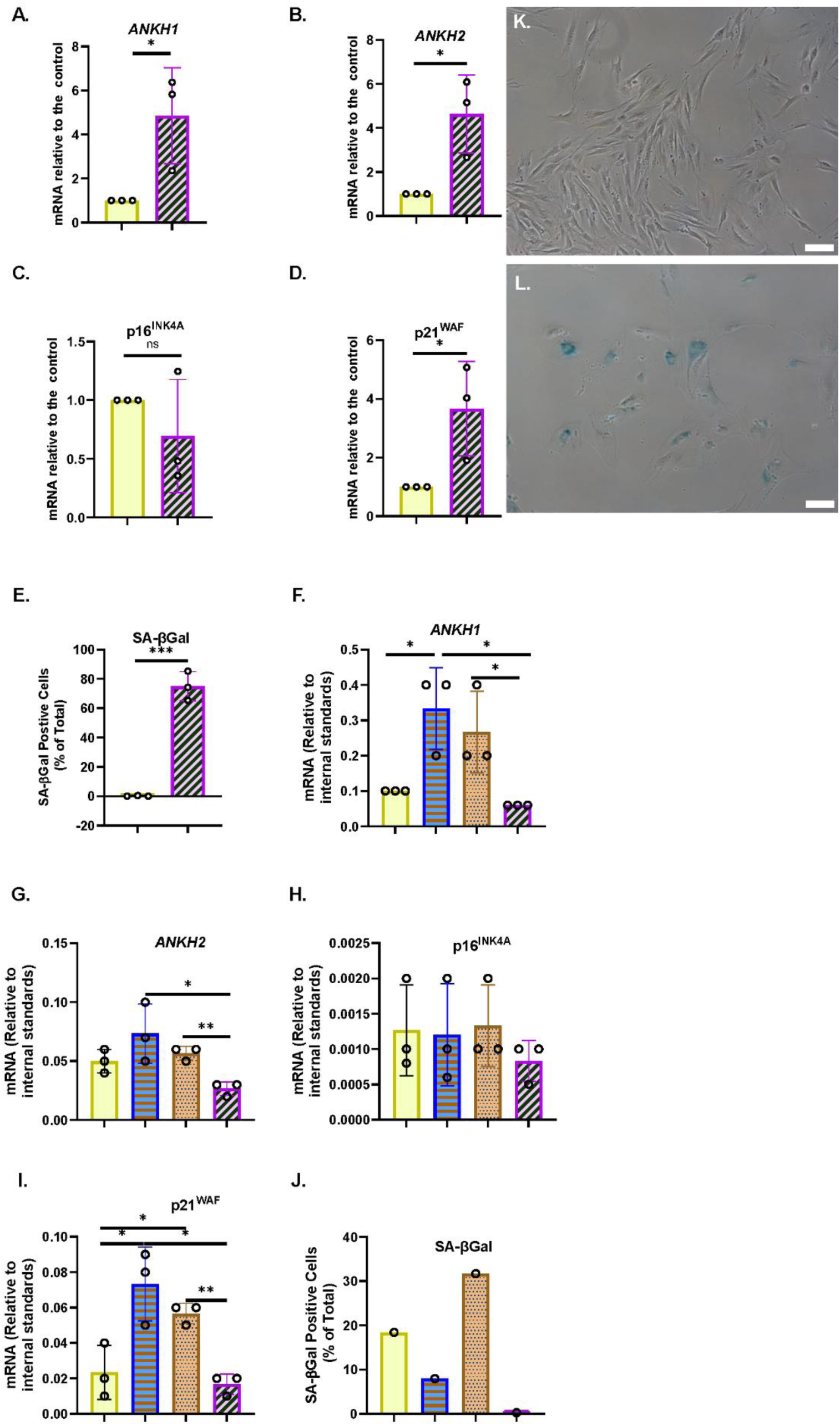
*ANKH* transcript is upregulated in PEsen human fibroblasts and downregulated by telomerase. A. The expression of *ANKH* mRNA in growing and PEsen NHOF-1 cells using primer set 1. B. As in A. but the results of primer set 2 in the same experiments. C. As in A. but the expression of the senescence marker p16^INK4A^in the same experiments. D. As in A. but the expression of the senescence marker p21^WAF1^ in the same experiments. E. As in A. but showing SA-β Gal expression. A-E. yellow plain bars = growing young cells 20.2-21 MPDs; blue horizontal striped bars = PEsen cells 68.2 MPDs. * = P< 0.05; ** = P < 0.01; *** = P < 0.001; * P > 0.05 <0.1; ns = not significant. The results are averages +/- standard deviation. N = 3. F. The expression of *ANKH* mRNA in NHOF-1 PURO, TERT-HA, DN TERT and TERT cell using primer set 1. N =3. G. As in F. but the results of primer set 2 in the same experiments. H. As in F. but the expression of the senescence marker p16^INK4A^in the same experiments. I. As in F. but the expression of the senescence marker p21^WAF1^ in the same experiments. J. As in F. but showing SA-β Gal expression. K. and L. Representative images of SA-βGal staining in E; K. Young growing NHOF-1 fibroblasts (20.2 MPDs), L. NHOF-1 cells (68.2 MPDs). Bar = 100µm F-J. A-E. Plain yellow bars = LATE PURO; blue horizontal striped bars TERT-HA, brown stippled bars DN TERT and purple left-right diagonally striped bars TERT all after completing 60 MPDs. * = P< 0.05; ** = P < 0.01; *** = P < 0.001; * > 0.05 < 0.1; ns = not significant. The results are averages +/- standard deviation. N = 4, except J.

We have already reported that the canonical function of telomerase reduces the accumulation of EC in parallel with bypassing senescence, so we next considered whether the transduction of NHOF-1 and BJ cells with *TERT,* but not the non-canonical variant *TERT-HA* ^31^ or the dominant-negative (catalytically dead in fibroblasts) mutant *DNTERT* ^32^, regulated the expression of *ANKH*. The results showed that in three independent NHOF-1 cultures and one BJ culture (Figure 2E, F; Supplementary Figure 4F, 4G) *TERT* expression reduced *ANKH* expression relative to the PEsen PURO controls. This occurred in parallel with p16^INK4A^ (Figure 2H), p21^WAF1^ (Figure 2I) and SA-βGal (Figure 2J-L; Supplementary Figure 4H). The *TERT-HA* and *DNTERT* controls, which fail to elongate telomeres ^32,33^, did not reduce *ANKH* transcript levels or any senescence marker in either NHOF-1 (Figure 2) or BJ cells (Supplementary Figure 4). All *TERT* constructs were expressed in late passage NHOF-1 (Supplementary Figure 5A) and BJ cells (Supplementary Figure 5B). Only *TERT* increased telomerase activity in NHOF-1 cells (Supplementary Figure 5C), and only *TERT* and *TERT-HA* increased telomerase activity in BJ cells (Supplementary Figure 5D), as previously reported ^31^.

### *ANKH* mRNA and EC are regulated by the p38 mitogen-activated kinase, Mitogen-activated protein kinase-activated protein kinase2/3 and the associated axis, but not by the Ataxia Telangiectasia Mutated kinase

We have previously reported that the kinetics of EC upregulation following the induction of IrrDSBsen are similar to that reported for the SASP and in particular, interleukin-6 (IL-6) ^34^. As p38 mitogen-activated kinase (p38MAPK) ^35^, Mitogen-activated protein kinase-activated protein kinase (MK2/3) ^36^ and Ataxia Telangiectasia Mutated (ATM ^37^) regulate the interleukins of the SASP, we tested whether pharmacological inhibition of these kinases would regulate EC and *ANKH*. Figure 3 shows the effect of the p38MAPK and MK2/3 inhibitors, SB203580 and PF-3633022 respectively, on young growing NHOF-1 and the same cells induced to senesce after the induction of IrrDSBsen. As reported previously, EC increased about 3-fold following IrrDSBsen and showed a trend for reduction with both drugs in both growing and IrrDSBsen groups, but the data were variable (Figure 3A). When the data was normalised to the controls for each experiment (Figure 3B) SB203580 showed a highly significant reduction of EC in IrrDSBsen cells, but not growing cells, and PF-3633022 inhibited EC significantly in both growing and IrrDSBsen groups. Taken together the data shows that EC is regulated by both p38MAPK and MK2/3.

**Figure 3.**
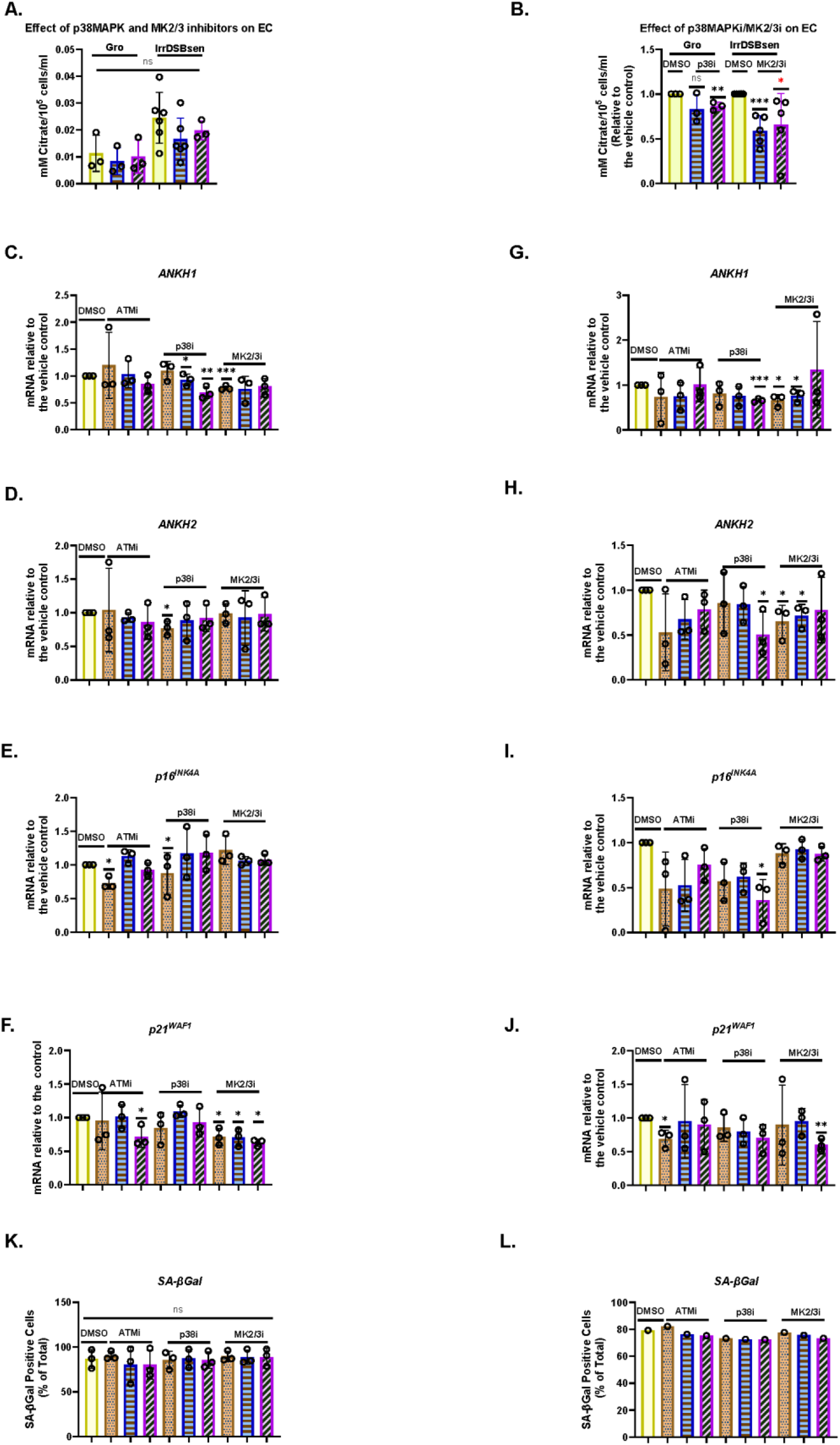
The p38 mitogen-activated kinase, the (*p38MAPK*)/MAPK-activated kinase2/3 (*MAPKAPK 2/3*) and the Ataxia Telangiectasia Mutated (*ATM*) kinase regulate EC, secreted IL-6 and *ANKH* mRNA expression differently in human fibroblasts. A, The effect of SB203580 (p38 inhibitor) and PF-3644022 (MK2/3 inhibitor) on EC in young growing and IrrDSBsen NHOF-1 cells (N =3). Plain yellow bars DMSO (vehicle) control; blue horizontal striped bars = SB203580 10 µM; purple left-right diagonally striped bars = PF- 3644022 2.5 µM B. The same data as in A and a separate series of experiments but normalised to DMSO vehicle control (N = 6). C-K. The effect of the kinase inhibitors SB203580 (p38MAPK) the PF-3644022 (MK2/3) and KU55933 (ATM) on *ANKH* transcript and senescence markers in IrrDSBsen cells. C-F BJ cells; G-J NHOF-1 cells. C. and G. The expression of *ANKH* mRNA using primer set 1. D. and H. The expression of *ANKH* mRNA using primer set 2. E. and me. The expression of p16*^INK4A^* mRNA. F. and J. The expression of p21^WAF1^ mRNA. K. and L. SA-β Gal expression (%). Plain yellow bars DMSO (vehicle) control; brown stippled bars 1 µM; blue horizontal striped bars 3 µM; purple left-right diagonally striped bars 10 µm * = P< 0.05; ** = P < 0.01; *** = P < 0.001; * P > 0.05 < 0.1; ns = not significant. The results are averages +/- standard deviation (N =3 except in B. where N =6). The experiments in A.- B. and C.- L. were performed at different times but using identical cell culture protocols and reagents.

Next, we evaluated the effect of SB203580 and PF-3633022, as well as the *ATM* kinase inhibitor KU55933, on *ANKH* mRNA expression and senescence markers p16^INK4A^ and p21^WAF1^ over a larger dose range in both IrrDSBsen BJ (Figure 3C-G) and NHOF-1 cells (Figure 3H-L). The data showed that in both BJ and NHOF-1, SB203580 (between 1 and 10 µM) inhibited *ANKH* expression when assayed by both *ANKH* primer sets (*ANKH1* and *ANKH2*); PF-3633022 inhibited *ANKH* expression between 1 and 2.5 µM, although this was clearer with primer set *ANKH*1. The biphasic response of human fibroblasts to PF-3633022 regarding IL-6 secretion has been documented previously ^38^ and the effect on *ANKH* mRNA expression was similar. KU55933 did not affect *ANKH* expression over this period. The above doses of the inhibitors did not consistently affect the expression of p16^INK4A^ or p21^WAF1^, but 10 µM SB-203580 did reduce p16^INK4A^ expression in NHOF-1 cells (Figure 3I) and PF-3633022 reduced p21^WAF1^ expression in BJ cells at 1-10 µM (Figure 3F) and at 10 µM in NHOF-1 (Figure 3J). None of the inhibitors affected the level of senescence as assessed by the SA-βGal assay over the 3-day period of incubation in either BJ or NHOF-1 cells (Figure 3G and 3L).

Taken together our data shows that EC in IrrDSBsen cells is regulated in parallel with IL-6 and *ANKH* by p38MAPK and MK2/3 independently of cell cycle arrest and the senescence phenotype, but not by the *ATM* kinase.

### EC is regulated by the p38 mitogen-activated kinase, Mitogen-activated protein kinase-activated protein kinase2/3 and the associated axis, independently of cell volume, the Ataxia Telangiectasia Mutated kinase and IL-6

To evaluate the effect of SB203580 and KU55933 on EC and IL-6 more thoroughly we exposed both growing and IrrDSBsen BJ cells to the drugs for 8 days individually and in combination. KU55933 alone had no effect on growing BJ cells but SB203580, either alone or in combination with KU55933 dramatically increased proliferation rate (Supplementary Figure 6A), reduced cell volume (Supplementary Figure 6B) and decreased SA-βGal (Supplementary Figure 6C), in line with previous reports that inhibiting p38MAPK antagonises cellular senescence ^39^. SB203580 and KU55933, but not KU55933 alone, also dramatically decreased EC (Supplementary Figure 6D) and the same combinations had similar effects in IrrDSBsen cells (Supplementary Figure 6D), which are likely due to SB203580 alone (see above). The same combinations did not affect IL-6 levels in growing BJ cells. However, KU55933 alone had a dramatic effect on IrrDSBsen cells, causing a many-fold induction in secreted IL-6 and this effect was almost ablated by SB203580 suggesting that the inhibition of ATM in senescent cells exacerbates signaling through p38MAPK to increase IL-6 under the conditions described here (Supplementary Figure 6E). Taken together these data sets also highlight the fact that EC and IL-6 are independently regulated in senescent cells, which is further supported by data below.

### EC and *ANKH* are not suppressed by physiological levels of steroids and are regulated independently from interleukin-6 in IrrDSBsen human fibroblasts

It has been reported that both cortisol (hydrocortisone – HC) and corticosterone (CST) within the physiological range suppress many components of the SASP, including IL-6 ^40^. This may explain why SASP factors are not that high in human plasma in clinical trials of senolytic drugs ^41^, during human chronological ageing ^42^ or in the telomeropathy DC ^19^. We therefore tested the effect of HC and CST on EC, secreted IL-6 ^40^ and *ANKH* expression over an 8-day period. In IrrDSBsen BJ cells IL-6 is significantly reduced by HC and CST, as previously reported by others in a different setting ^40^, but EC is not (Figures 4A-4C). This supports the argument that EC and IL-6 are regulated independently. Interestingly, *ANKH* expression tended to be increased, especially with high doses of HC, rather than decreased (Figure 4D and 4E); there was no significant effect on p16^INK4A^ (Figure 4F), p21^WAF^ (Figure 4G) or SA- βGal (Figure 4H) and so the steroid effects on IL-6 were independent of senescence. Comparable results were obtained with NHOF-1 cells (Supplementary Figure 7).

**Figure 4.**
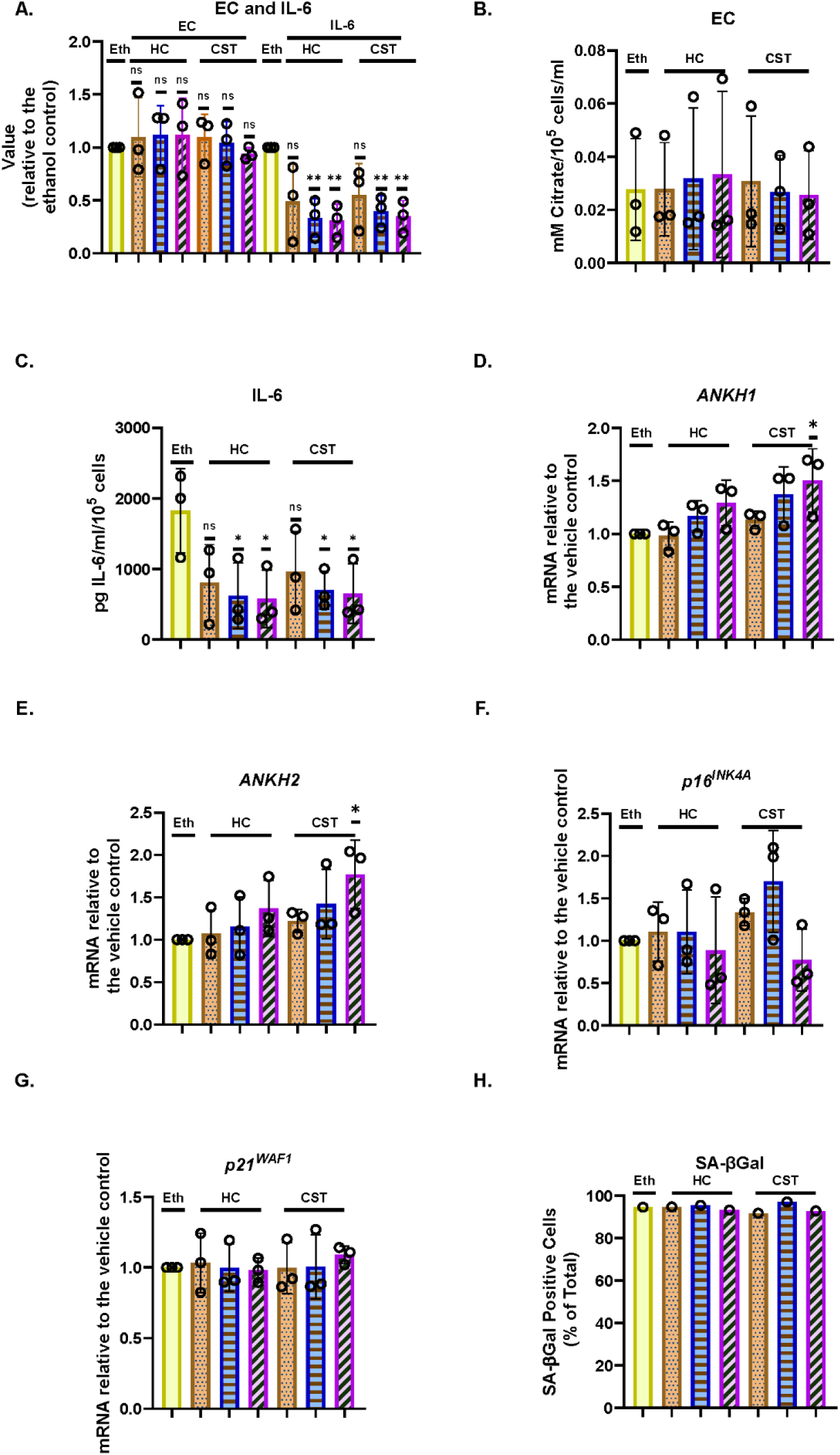
The effect of steroids on the EC, *ANKH* expression and secreted IL-6 in IrrDSBsen human fibroblasts. A. Shows the EC and IL-6 levels in the medium of IrrDSBsen BJ fibroblasts after treatment with the indicated doses of HC and CST. B. Shows the absolute EC levels in mM citrate per 10^5^ cells per mL from the experiment in A. C. Shows the absolute levels of IL-6 in pg/mL per 10^5^ cells from the experiment in A. D. The effect the indicated doses of HC and CST on the expression of *ANKH* mRNA using primer set 1. E. As in D. but the results of primer set 2 in the same experiments. F. As in D. but the expression of the senescence marker p16^INK4A^in the same experiments. G. As in D. but the expression of the senescence marker p21^WAF1^ in the same experiments. H. As in D. but the expression of senescence marker SA-βGal in the same experiments. Plain yellow bars DMSO (vehicle) control; brown stippled bars 1 µM; blue horizontal striped bars 3 µM; purple left-right diagonally striped bars 10 µm * = P< 0.05; ** = P < 0.01; *** = P < 0.001; * P > 0.05 < 0.1; ns = not significant. The results are averages +/- standard deviation. A. to G. N = 3. H. N=1. The experiments in A.- C. and D.- L. were performed at separate times but using identical protocols and reagents.

### interleukin-1 alpha inhibits the production of EC and *ANKH* expression in proliferating and IrrDSBsen human fibroblasts

Next, we tested the effect of the SASP regulator interleukin 1 alpha (IL-1α), which was reported to positively regulate IL-6 ^40^, on EC, secreted IL-6 and *ANKH* expression in BJ cells. Under the conditions described here, IL1α actually stimulated BJ proliferation in young cells (Figure 5A) in parallel with a strong induction of secreted IL-6 (Figure 5B), as previously reported ^40^, but reduced EC (Figure 5C) in parallel with *ANKH* expression (Figure 5D and 5E) whilst having a minimal effect on p16^INK4A^ (Figure 5F), p21^WAF^ (Figure 5G) and SA-βGal (Figure 5H). We also tested the effect of IL-1α on IrrDSBsen fibroblasts (Supplementary Figure 8). IL-1α strongly stimulated the secretion of IL-6, an effect that was antagonised by HC, whereas the effect on EC was the reverse. In this separate set of experiments, HC alone also inhibited IL-6 secretion (Supplementary Figure 8A) but had no effect on EC (Supplementary Figure 8B).

**Figure 5.**
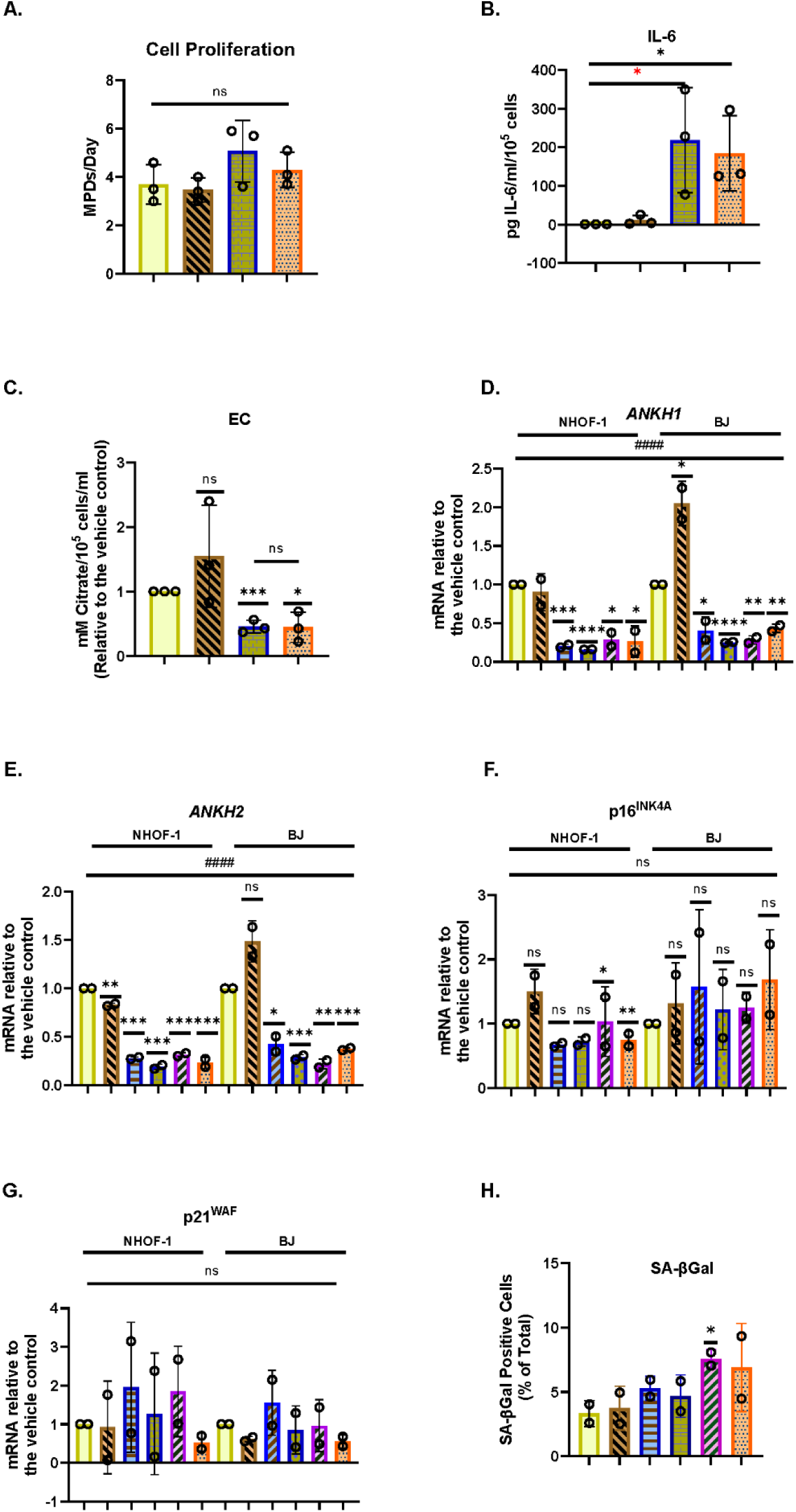
The effect of IL-1α on EC, *ANKH* expression and secreted IL-6 in proliferating human fibroblasts. The figure shows the effects of an 8-day treatment with the indicated doses of IL-1α and HC on proliferating NHOF-1 and BJ cells. A. Shows the rate of proliferation of BJ cells in MPDs per day. B. Shows the absolute levels of IL-6 in pg/mL per 10^5^ cells in BJ cells. C. Shows the absolute EC levels in mM citrate per 10^5^ cells per mL in BJ cells. D. The effect the indicated doses of IL-1α and 300 nM HC and on the expression of *ANKH* mRNA in BJ and NHOF-1 cells using primer set 1. E. As in C. but the results of primer set 2 in the same experiments. F. As in C. but the expression of the senescence marker p16^INK4A^in the same experiments. G. As in C. but the expression of the senescence marker p21^WAF1^ in the same experiments. H. The expression of senescence marker SA-βGal in the same BJ cell experiments. A.- C. Yellow bars control; brickwork dark blue bars 1ng/mL IL-1α; right-left brown bars 300nm HC; stippled orange bars = 1 ng/mL IL-1α and 300nm HC. D.-H. Plain yellow bars control; horizontal light blue bars 0.5 ng/mL IL-1α; brickwork blue bars 1ng/mL IL-1α; left-right diagonally striped purple bars 2.5 ng/mL IL-1α; right-left diagonally striped brown bars 300nm HC; stippled orange bars = 2.5 ng/mL IL-1α and 300nm HC * = P< 0.05; ** = P < 0.01; *** = P < 0.001; ns = not significant. The results are averages +/- standard deviation. A. and B. N = 3. C-G. N=2 for each cell line, (N =4 in total). All data was also analysed by ordinary two-way ANOVA (hatched lines). # = P< 0.05; ## = P < 0.01; ### = P < 0.001; #### = P < 0.0001; ns = not significant. The experiments in A.- C. and D.- H. were performed at different times but using identical cell culture protocols and reagents.

### The senescence inducer sodium butyrate induces IL-6 and proliferation arrest independently of EC

In addition, we tested another regulator of IL-6, sodium butyrate (NaB), which had been reported to induce secreted IL-6 independently of DNA double strand breaks, or senescence^43^. Again, the data showed that in BJ cells and two oral fibroblast lines (NHOF-2 and NHOF- 7) a four-day treatment of the cells with 1-4 mM NaB inhibited cell proliferation (Supplementary Figure 9A), caused cell flattening, increased cell volume and induced IL-6 secretion (Supplementary Figure 9B), without significantly affecting histone acetylation (H3K27ac) (Supplementary Figure 9C) and collagen (Supplementary Figure 9D). 53BP1 levels were slightly increased in all cell lines at but only significantly at 1mM NaB (Supplementary Figure 9E) and SA-βGal levels were increased in BJ cells (Supplementary Figure 9F). In contrast, p16^INK4A^ (Supplementary Figure 9G) or p21^WAF1^ (Supplementary Figure 9H) mRNA expression was not increased. In all cell lines, EC was modestly and insignificantly induced over this time frame, and not as high as in IrrDSBsen cells (Supplementary Figure 9I and Supplementary Figure 9J). In BJ cells, *ANKH* expression was, if anything, reduced (Supplementary Figure 9K; Supplementary Figure 9L).

These results confirm the association between EC and *ANKH* expression and the independent regulation of EC from IL-6.

### EC and hydrogen peroxide are negatively regulated by p53 independently of *ANKH* expression and are independent of the senescence effectors p16^INK4A^ and p21^WAF^^1^

To evaluate the effect of p53 on EC regulation we assessed a panel of human wild type, p53 null and p21^WAF1^ null embryonic lung fibroblasts (Loxo26WT, Loxo26 p53-/- and Loxo 26 p21^WAF1^ -/-). The results showed that whether growing, or following IrrDSBsen, Loxo26 p53-/- showed much higher levels of EC (Supplementary Figure 10A) and hydrogen peroxide (Supplementary Figure 10B) than their wild type and/ or p21^WAF1^ null counterparts. qPCR data confirmed the absence of p53 in the p53 -/- cells (Supplementary Figure 10C) and showed that the p21^WAF1^ null Loxo26 cells had silenced p16^INK4A^ in addition to p21^WAF^ during passage after receipt (Supplementary Figures 10D and 10E). This means that EC regulation was independent of the senescence effectors p21^WAF1^ and p16^INK4A^. However, EC was restrained by p53, analogous to its reported regulatory function on the SASP (Supplementary Figure 10A -see also reference ^37^)

The ability of p53 to restrain the accumulation of hydrogen peroxide (Supplementary Figure 10B) was consistent with its reported antioxidant function ^44^ but under the conditions reported here there was no effect of p21^WAF1^ and p16^INK4A^ on hydrogen peroxide levels. To evaluate whether the restraining effect of p53 on EC was mediated by *ANKH,* we assessed the cell line panel for *ANKH* expression but found no consistent upregulation of *ANKH* in p53-/- Loxo26 cells or p21^WAF1^/p16^INK4A^ -/- Loxo26 cells. (Supplementary Figure 10F and 10G). There was no dramatic effect of p53 deletion on SA-βGal levels (Supplementary Figure 10H) although p16^INK4A^ and p21^WAF1^ levels were significantly reduced (Supplementary Figures 10D and 10E). We also evaluated the effect of inducible *TP53* shRNAs on the expression of *ANKH* mRNA and EC in growing NHOF-1 fibroblasts. shRNA #2 did reduce both *TP53* mRNA and that of its downstream target p21^WAF^ relative to the NT control by 59% and 53% respectively following induction by DOX (Supplementary Figure 10I) but did not upregulate *ANKH* mRNA (Supplementary Figure 10I) or EC in these cells (Supplementary Figure 10J). *TP53* shRNA #2 significantly downregulated *ANKH* mRNA which may indicate a role for *TP53* in regulating *ANKH* independently of EC. Taken together the data suggests that *TP53* restrains EC independently of senescence or cell cycle inhibitors in certain fibroblast lines, analogous to its effects on the SASP, but may do so independently of *ANKH* expression.

### Sodium butyrate induces senescence and EC accumulation independently of *ANKH* expression

To evaluate the effect of senescence inducers other than those mediated by IrrDSBs or short telomeres, we assessed the ability of NaB to induce EC and senescence in the Loxo26 panel and p21^WAF1^/p16^INK4A^. We found that EC was elevated in the wild type cells following 3 weeks of 0.5 mM NaB treatment, but not in BJ cells (Supplementary Figure 11A). This NaB effect was dependent on p53 and p21^WAF1^/p16^INK4A^. We then evaluated whether the effect on EC was associated with *ANKH* expression. We treated the Loxo26 panel, NHOF-1, BJ and IMR90 cells for 17 days and observed the classical senescence effect of cell flattening, increased cell size, increased p16^INK4A^ (Supplementary Figure 11B) and p21^WAF1^ (Supplementary Figure 11C) expression and in BJ and Loxo26 cells, increased SA-βGal (Supplementary Figure 11D). However, despite inducing features of senescence NaB did not induce *ANKH* expression in BJ, NHOF-1 or Loxo26, and only did so in IMR90 at high doses (Supplementary Figures 11E and 11F). This indicates that the induction of EC by NaB-induced senescence is variable between different fibroblast lines but is not in any case associated with *ANKH* induction.

### Transforming growth factor β (TGFβ) induces *ANKH* expression prior to inducing senescence

TGFβ will induce reversible cell cycle arrest in many cell types ^45^. The TGFβ family has also been implicated in the transfer of the senescence phenotype between cells ^46^. To examine the effect of TGFβ on *ANKH* expression and senescence we treated the cells for 4 days with 1 −8 ng/mL of TGFβ1 in 4 mM HCl (Figure 6). The data showed that TGFβ1 significantly induced *ANKH* in NHOF-1 at low doses (Figure 6A and 6B) and showed a trend for induction in BJ cells (Figure 6A, 8B) in parallel with increased p16^INK4A^ (Figure 6C) and p21^WAF1^ (Figure 6D) expression. However, there was no evidence of senescence as assessed by SA-βGal (Figure 6E). These data indicate that TGFβ1 mediates *ANKH* expression in parallel with fibroblast activation prior to, or independently from senescence ^47^. Next, to test the role of the TGFβ family members in regulating *ANKH* and senescence markers in established IrrDSBsen fibroblasts, we treated three lines of IrrDSBsen fibroblasts with two TGFβ type I receptor kinase inhibitors that had been reported to suppress the SASP ^46^. Figures 6F and 6G show that although the effect was small, both inhibitors significantly suppressed *ANKH* expression by 15-30% in all three lines without reversing the levels of p16^INK4A^ (Figure 6H), p21^WAF^ (Figure 6I) and SA-βGal (Figure 6J), at least in the case of inhibitor I. However, there was a significant reduction in p21^WAF^ expression (Figure 6I) and a trend for SA-βGal reduction Figure 6J) in the case of inhibitor II. We conclude that the TGFβ family contributes to the regulation of *ANKH* in IrrDSBsen fibroblasts.

**Figure 6.**
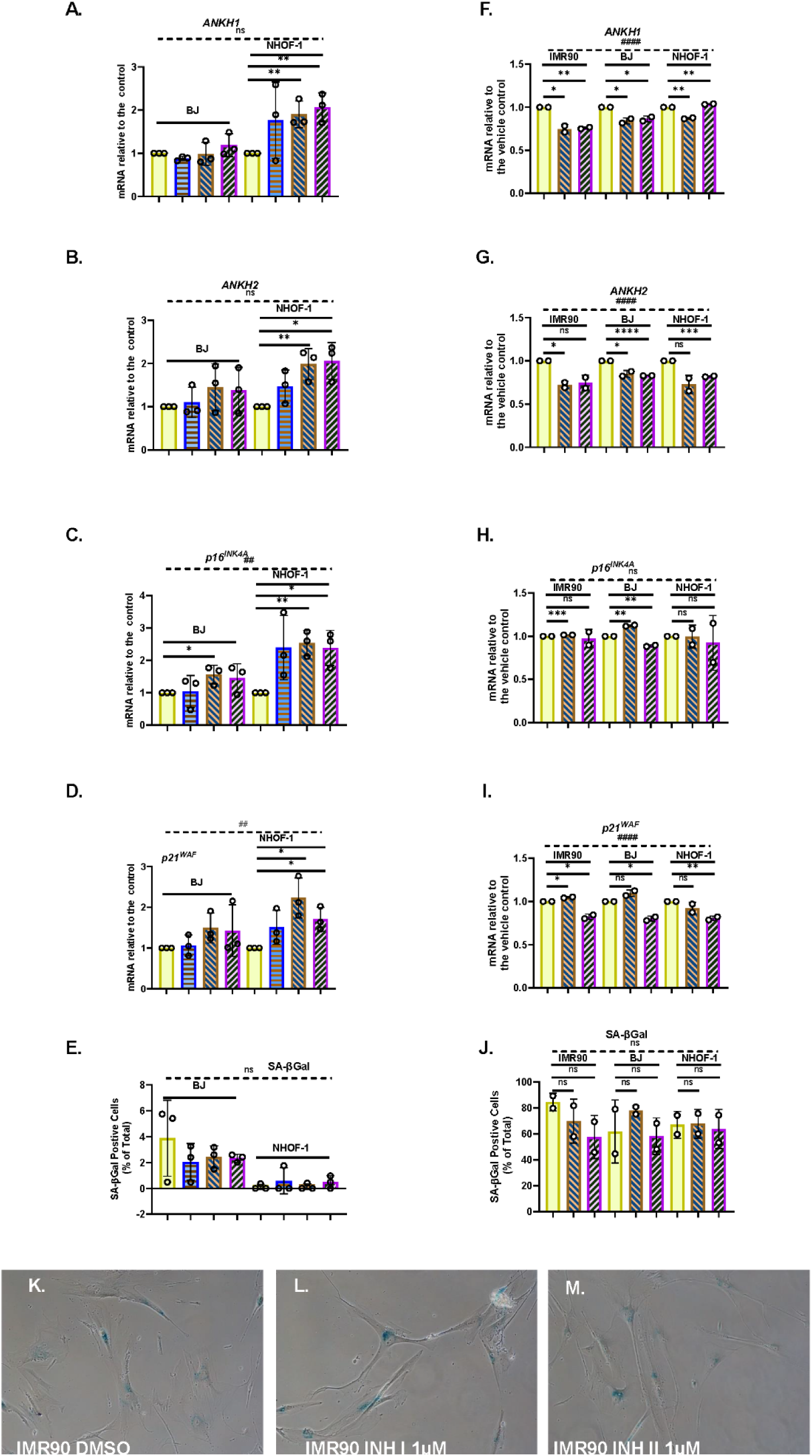
The effect of TGFβ on cellular senescence and the expression of *ANKH* in human fibroblasts. A.- E. The effect of TGFβ at doses of 1-10ng/mL for 4 days on NHOF-1 and BJ fibroblasts on the expression of ANKH and senescence markers in growing young fibroblasts. A. The expression of *ANKH* mRNA using primer set 1. B. As in A. but the results of primer set 2 in the same experiments. C. As in A. but the expression of the senescence marker p16^INK4A^in the same experiments. D. As in A. but the expression of the senescence marker p21^WAF1^ in the same experiments. E. As in A. but the expression of senescence marker SA-βGal in the same experiments. Plain yellow bars control; blue horizonal striped bars 1 ng/mL; brown right-left diagonally striped bars 3 ng/mL ; purple left-right diagonally striped bars 10ng/mL * = P< 0.05; ** = P < 0.01; *** = P < 0.001; ns = not significant. The results are averages +/- standard deviation. N = 3 for each line (N = 6 in total). F. - J. The effect of TGFβ type I receptor kinase inhibitors on the expression of ANKH and senescence markers in IrrDSBsen IMR90, BJ and NHOF-1 fibroblasts. F. The expression of *ANKH* mRNA using primer set 1. G. As in A. but the results of primer set 2 in the same experiments. H. As in A. but the expression of the senescence marker p16^INK4A^in the same experiments. I. As in A. but the expression of the senescence marker p21^WAF1^ in the same experiments. J. As in A. but the expression of senescence marker SA-βGal in the same experiments. K. - M. Representative images of the SA-βGal staining of IMR90 cells in J. Plain yellow bars vehicle control (0.1% DMSO); brown right-left diagonally striped bars TGFβ type I receptor kinase inhibitor I at 1µM; purple left-right diagonally striped bars TGFβ type I receptor kinase inhibitor II at 1µM. * = P< 0.05; ** = P < 0.01; *** = P < 0.001; ns = not significant. The results are averages +/- standard deviation. N = 2 for each cell line (N = 6 in total). All data was also analysed by ordinary two-way ANOVA (hatched lines). # = P< 0.05; ## = P < 0.01; ### = P < 0.001; #### = P < 0.0001; ns = not significant.

### *ANKH* and EC are not upregulated in IrrDSBsen and PEsen keratinocytes or during keratinocyte differentiation, indicating cell type specificity in citrate export

As senescent mammary epithelial cells were reported not to shift their energy metabolism towards glycolysis and EC, albeit under very different culture conditions to those we used previously ^21^, we tested both EC accumulation and *ANKH* expression following IrrDSBsen in both 3T3-supported cultures and serum-free commercial media. We also tested the effect of ROCK inhibitor Y27632 (ROCKi) in the 3T3 system which has been reported to induce telomerase ^48^, thus replicating more accurately conditions in the epidermal basal layer^49^. In all five epidermal keratinocyte lines IrrDSBsen resulted in a downregulation of EC relative to controls (Supplementary Figure 12A) and the inclusion of ROCKi in the two 3T3-supported cultures did not change the result (Supplementary Figure 12B). We also evaluated the effect of PEsen on *ANKH* mRNA expression in regular 0.09 mM calcium medium after 4 and 7 days after the induction of differentiation and stratification by 0.4 mM and 1.0 mM calcium chloride addition to serum-free keratinocyte medium. There was no consistent increase in *ANKH* (Supplementary Figure 12C and Supplementary Figure 12D), whilst p16^INK4A^ (Supplementary Figure 12E) and p21^WAF^ (Supplementary Figure 12F) mRNA increased as reported previously for p16^INK4A^ protein following PEsen ^50^ in all conditions after 4 days, but not after the induction of differentiation (see below). We next addressed the role of differentiation and stratification on EC accumulation and *ANKH* expression in young cells by manipulating the calcium chloride concentration as above and monitoring the extent of differentiation by involucrin (*IVL*) and keratin 10 (*K10*) expression. Supplementary Figure 13 shows that in all five keratinocyte lines there was no consistent increase in EC (Supplementary Figure 13 A, B) following the induction of differentiation. In four of the five lines, there was no increase in *ANKH* (Supplementary Figures 13C, D), p16^INK4A^ (Supplementary Figure 13E) or p21^WAF^ (Supplementary Figure 13F) following the induction of differentiation as assessed by IVL (Supplementary Figure 13G), or stratification by K10 (Supplementary Figure 13H). Taken together our data shows there is no evidence that PEsen, IrrDSBsen or terminal differentiation regulates either EC or *ANKH* expression in human epidermal keratinocytes.

### *ANKH* mRNA is regulated in other types of IrrDSBsen and PEsen cell types relevant to aging and age-related disease and in chronologically aged mouse tissue

Next, we tested the relevance of *ANKH* upregulation in senescent cell types that are relevant to age-related disease in mice such as astrocytes (cognitive decline ^51^), adipocytes (frailty ^52^ and type II diabetes ^53^) and myoblasts (frailty ^54^). We now show that *ANKH* was upregulated in adipocytes following confluence-induced adipocyte differentiation (Figure 7A, 7B) and senescence (as assessed by high p16^INK11A^ and p21^WAF1^; Figure 7C and 7D); but not in PEsen or IrrDSBsen pre-adipocytes (Supplementary Figure 14). We, and others, have shown that long-term quiescence induced by confluence can induce senescence ^55,56^.

**Figure 7.**
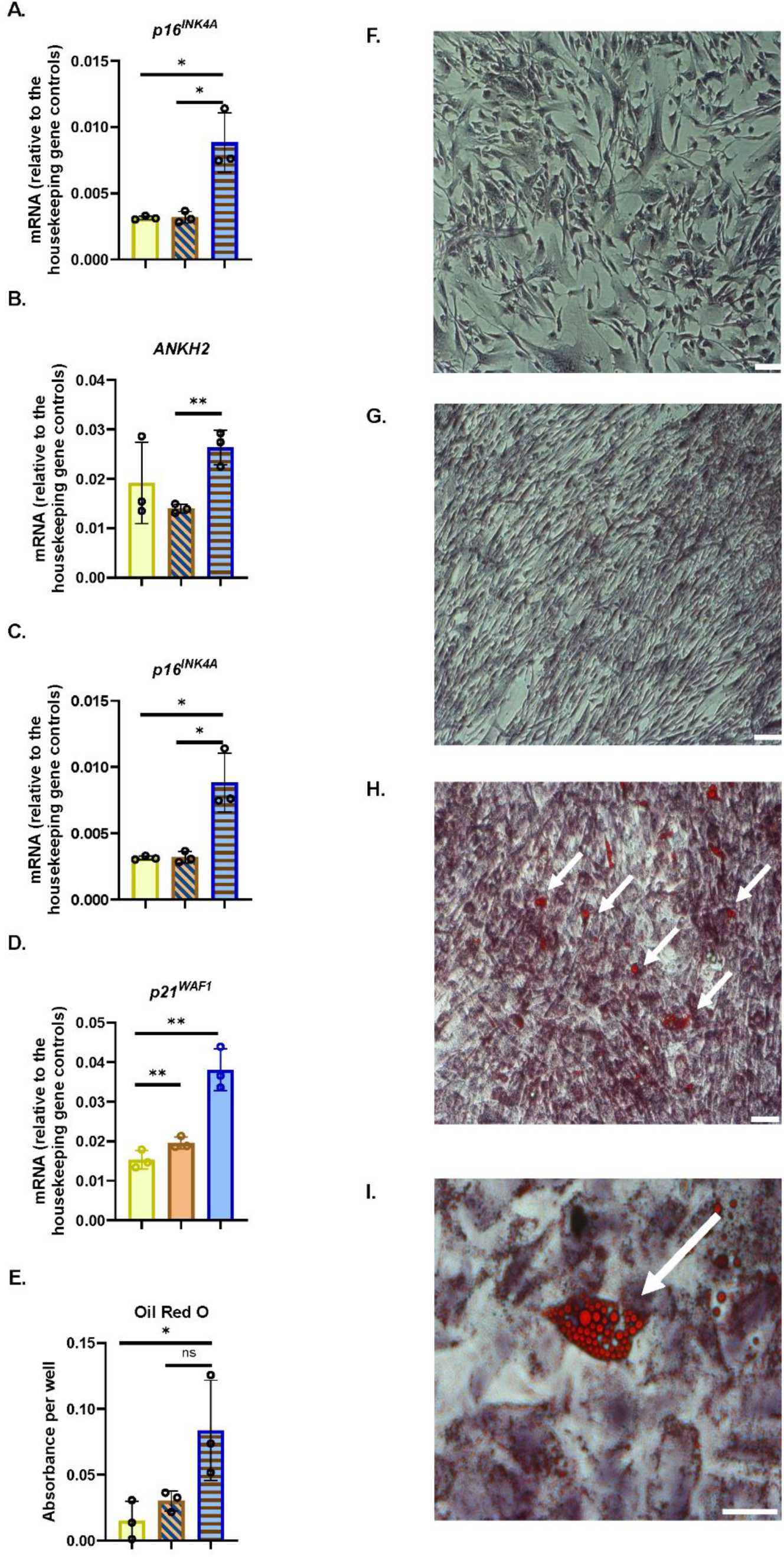
*ANKH* mRNA is upregulated in differentiated adipocytes following confluence-induced senescence. A. The expression of *ANKH* mRNA using primer set 1. B. The expression of *ANKH* mRNA using primer set 2. C. The expression of p16^INK4A^ mRNA. D. The expression of p21^WAF1^ mRNA. E. Oil Red O staining to indicate adipocyte differentiation. Plain yellow bars = young growing control; brown right-left diagonally striped bars = young confluent (4 days control); blue horizontal striped bars = confluent (15 days) senescent differentiated adipocytes. * = P< 0.05; ** = P < 0.01; *** = P < 0.001; * P > 0.05 <0.1; ns = not significant. The results are averages +/- standard deviation. N = 3. F. Young growing control. G. Young confluent control. H. Young confluent differentiated showing cells with Oil Red-positive lipid droplets (arrows). Bar = 100µM. I. A higher power image of H. Bar = 100µM.

Human myoblasts display markers of senescence and telomere-associated foci with increased chronological age ^54^. Human myoblasts cannot be immortalised by telomerase alone, but we were able to compare senescent and pre-senescent human myoblasts (HMC) transduced with the catalytic subunit of telomerase, *TERT*, (HMC *TERT* POOL or HMC *TERT* clone T2) and control (HMC) myoblasts. We were also able to compare these cells with a clone of *TERT*-transduced myoblasts that had spontaneously lost p16^INK4A^ (HMC *TERT* T15 ^57^). In senescent *TERT*-expressing cells, *ANKH* was upregulated following PEsen (Supplementary Figure 15A, B) in parallel with p16^INK4A^ ^(^Supplementary Figure 15C), p21^WAF^ (Supplementary Figure 15D) and SA-βGal (Supplementary Figure 15E). However, *ANKH* and all markers except p21^WAF^ were strongly downregulated following immortalisation in HMC *TERT* T15. These data also show that *ANKH* is associated with senescence in human myoblasts.

Importantly, human astrocytes express *ANKH* (Figure 8A, 8B, 8G, 8H and *ANKH* is over-expressed in parallel with SA-βGal, p16^INK4A^and p21^WAF1^ in both IrrDSBsen (Figure 8A-E), together with old and PEsen human astrocytes (Figure 8F-J, see also Supplementary Figure 16). Upon PEsen the astrocytes became much flatter, lost their bipolar appearance, and increased in size (Figure 8N) compared to young astrocytes (Figure 8L). Several SASP cytokines were upregulated in PEsen astrocytes along with Complement Factor 3, indicative of astrocyte activation (Supplementary Figure 16).

**Figure 8.**
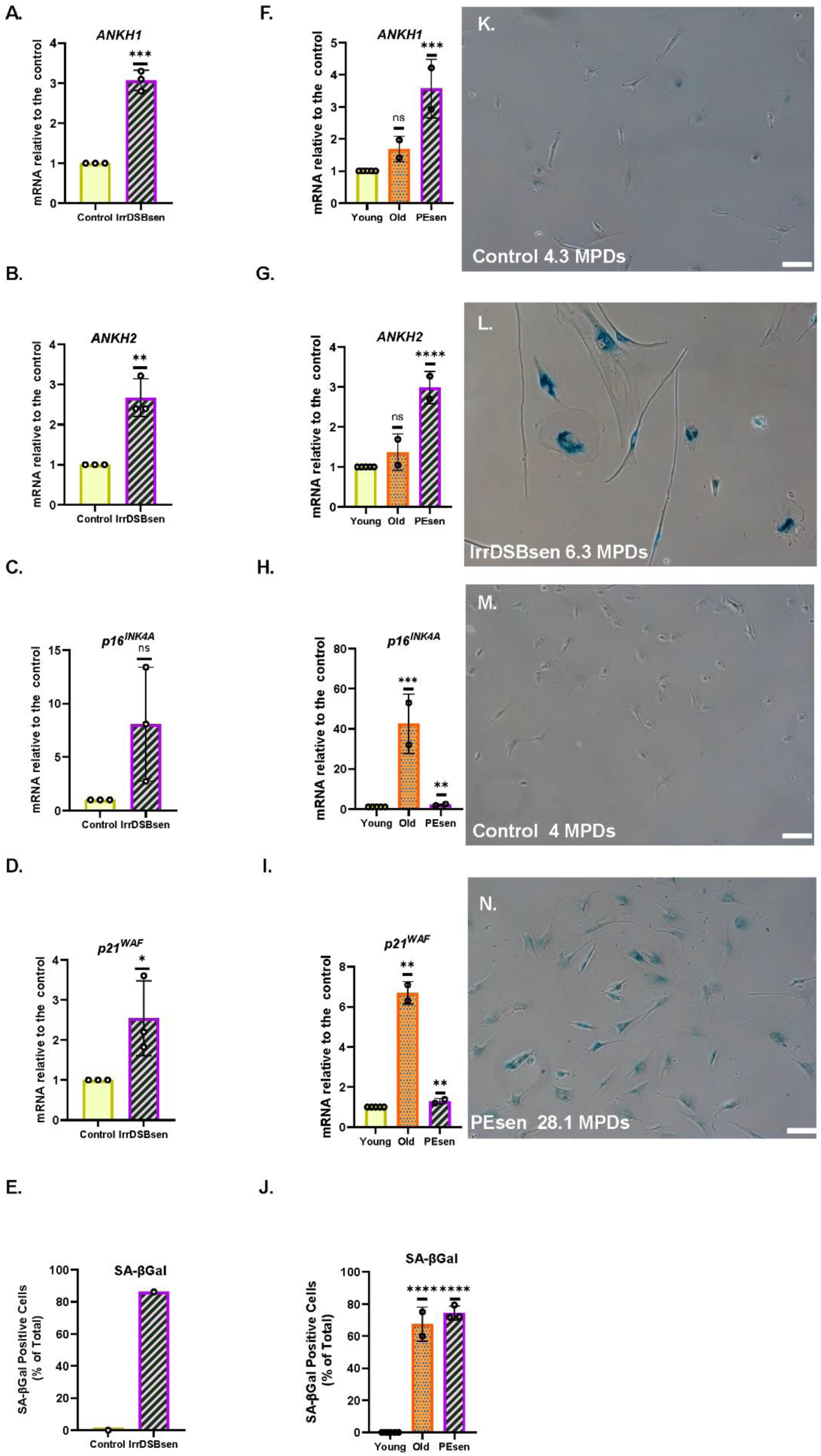
*ANKH* mRNA is upregulated in IrrDSBsen and PEsen astrocytes. A.- E. IrrDSBsen; F-J. PEsen. A. and F. The expression of *ANKH* mRNA using primer set 1. B. and G The expression of *ANKH* mRNA using primer set 2. C. and H. The expression of p16*^INK4A^* mRNA. D. and I. The expression of p21^WAF1^ mRNA. E. and J. SA-β Gal expression (%). A.-J. Plain yellow bars = Young growing control; purple left-right diagonally striped bars A.-E. = IrrDSBsen; F-J PEsen astrocytes (27-28.9 MPDs). F-J. orange stippled sbars = old (22-23 MPDs) astrocytes. * = P< 0.05; ** = P < 0.01; *** = P < 0.001; **** = P < 0.0001; * P > 0.05 < 0.1; ns = not significant. The results are averages +/- standard deviation. A- D N = 3; E is the result of a single experiment; F-J N = 2 (see also Supplementary Figure 16). K. and L. Representative images of SA-βGal staining in E; K. Young growing astrocytes (4.3 MPDs), L. IrrDSBsen astrocytes (6.3 MPDs). M. and N. Representative images of SA-βGal staining in J. M. Young growing astrocytes (4.0 MPDs), N. PEsen astrocytes (28.1 MPDs): Bar = 100µm.

### *Ank* is downregulated following mouse chronological ageing in parallel with senescence markers

To assess the role of *ANKH* (*Ank*) in mouse chronological ageing, we examined the levels of the *Ank* transcript in C57BL6 mouse tissues where body fluid levels of citrate have been reported to be elevated at advanced chronological age. Figures 9A and 9B show that *Ank* is highly expressed in brain tissue relative to liver or kidney. In all three types of tissue, there was a trend for decreased levels of *Ank* transcript with chronological age, which was significant in liver (Figure 9A) and the brain (Figure 9B). Older liver and kidney tissues do show increased *Ink4a* and *itgb3* transcript levels ^58^ confirming the increased presence of senescent cells ^58^. As regards the brain, immunohistochemistry revealed increased levels of cells in older mouse tissues expressing p53, p19^ARF^, p21^WAF1^ (Figure 9C), p16^INK4A^, and ɤH2AX/53BP1) double stained foci (Figure 9D). All aged brain cells (mainly microglia and neurons) and neurons alone, showed a trend for increased telomere-associated foci with age (Figure 9D). Older mice showed increased levels of the SASP cytokines IL-1β, and TNFα and reduced levels of IL-10 in brain tissue (Figure 9E) and increased IL-6, TNFα and CXCL1 in serum (Figure 9F). Images of the immunostaining in Figures 9C and 9D are shown in Figure 10. These results are consistent with the repression of EC and *ANKH* by IL-1α in human fibroblasts and the elevated levels of cytokines in the aged mouse serum and tissues ^58^, including brain (Figure 9E).

**Figure 9.**
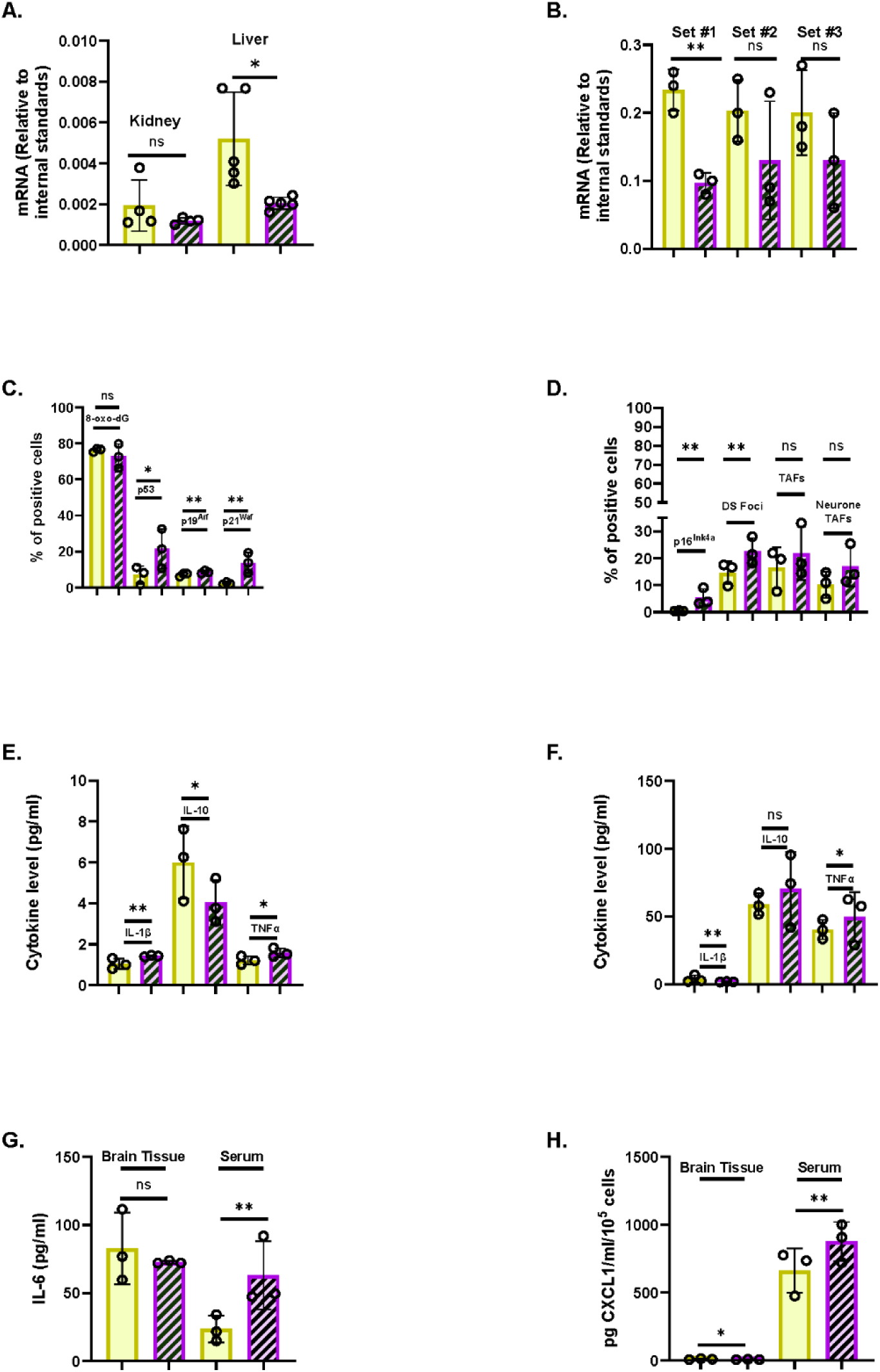
Aged mouse tissues show increased levels of senescence markers and inflammatory cytokines in parallel with reduced levels of *ANKH* mRNA. A. *ANKH* mRNA levels in young (4 month) and old (25 month) kidney (N = 4) and liver (N =5) tissue. B. *ANKH* mRNA levels in young (2.5 month) and old (17 month) brain tissue (N =3). C. Immunofluorescence staining of adult and aged C57BL/6 mice. (A) Percentage of cells expressing 8-oxo-dg, (B) Percentage of cells expressing p53, (C) Percentage of cells expressing p21^Waf1^, (D) Percentage of cells expressing p19^Arf^. D. Immunofluorescence staining of adult and aged C57BL/6 mice. (A) Percentage of cells expressing p16^Ink4a^, (B) Percentage of cells expressing γH2AX.53BP1 double foci (DF), (C) Percentage of cells expressing telomere-associated foci (TAF), (D) Percentage of neurons expressing TAF. E. Different cytokine levels (pg/mL) in brain samples. Interleukin-1 beta (IL-1β), Interleukin-10 (IL-10), Tumour necrosis factor alpha (TNF-α). F. Different cytokine levels (pg/mL) in serum samples. (Interleukin-1 beta (IL-1β), Interleukin-10 (IL-10), (C) (E) Tumour necrosis factor alpha (TNF-α). G. IL-6 levels (pg/mL) in brain and serum samples. H. CXCL1 levels (pg/mL) in brain and serum samples. A. and B. Results analysed by the Student’s two-tailed unpaired test. * = P< 0.05; ** = P < 0.01; *** = P < 0.001; ns = not significant. The results are averages +/- standard deviation. C.-H. Results represent the median, minimum and maximum. Data were analysed using the Mann-Whitney U test; *p<0.05. 3 animals for Adult C57BL/6, and 3 animals for Aged C57BL/6. A. Plain yellow bars = young (4-month-old mice); purple left-right diagonally striped bars = old (25-month-old mice). B. – H. Plain yellow bars = young (2.5-month-old mice); purple left-right diagonally striped bars = old (17-month-old mice).

**Figure 10.**
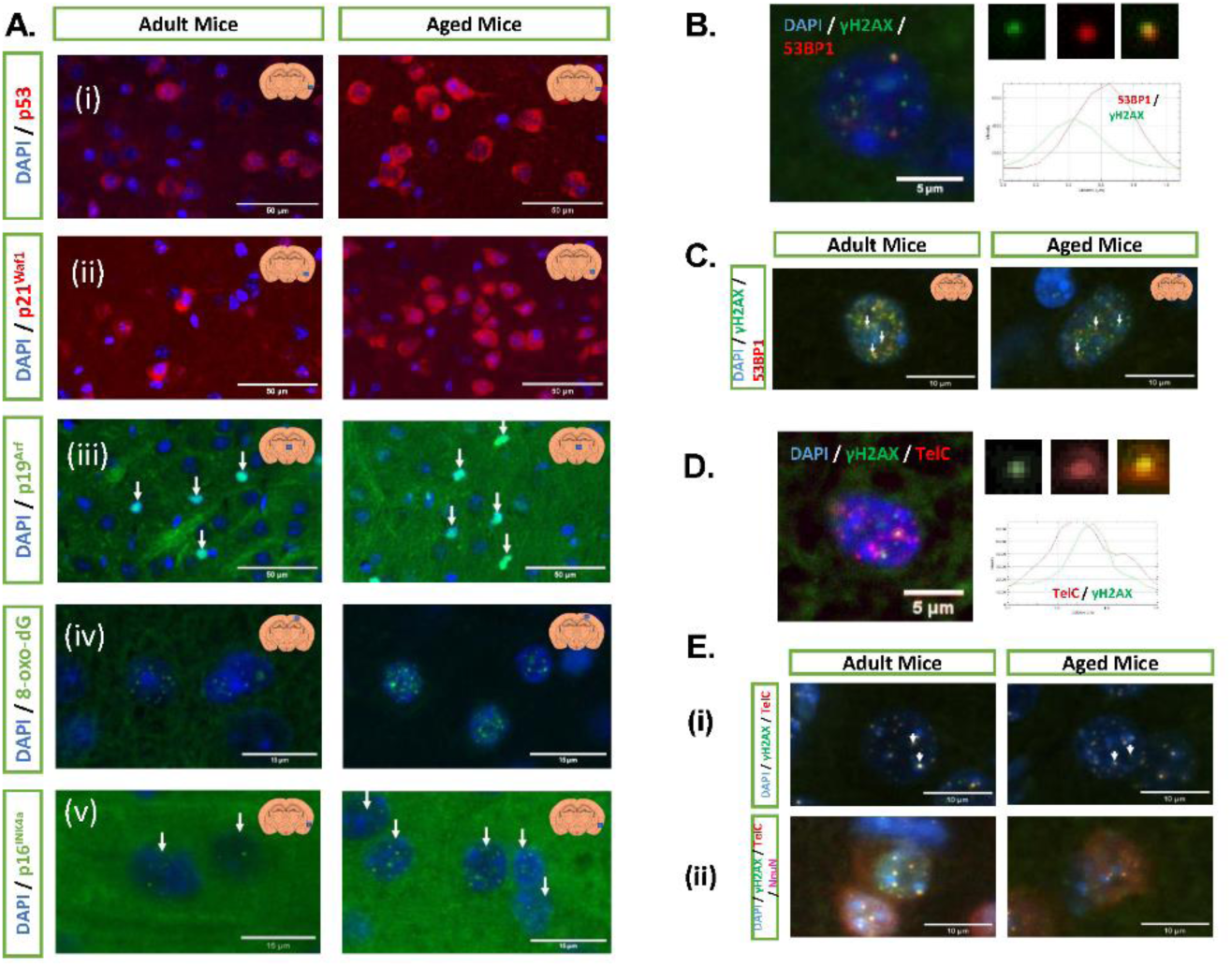
Immunofluorescence staining of multiple cellular senescence markers. A. Representative images of (i) p53 signal in the cerebral cortex, (ii) p21^Waf1^ signal in the cerebral cortex, (iii) p19^Arf^ signal in the thalamus, (iv) 8-oxo-dG signal in the cerebral cortex, (v) p16^INK4a^ signal in the cerebral cortex. All images were taken at 20X magnification. Bar = 50µm. B. Example of γH2AX and 53BP1 signals in a nucleus, the arrow demonstrates a DF, and the histogram displays their overlap. C. Representative images of both groups. D. Examples of TAF, γH2AX and TelC signals in a nucleus, the arrow demonstrates a DF, and the histogram displays their overlap. E. Representative images of TAF in a (i) nucleus, (ii) neuron. Images from B. to E. were taken at 60X magnification. Bar = as indicated in the image.

## Discussion

There is now considerable evidence that energy ^59^ and citrate uptake ^8–12,60^ are important for both ageing and age-related disease. For example, we have previously reported that EC is regulated by the canonical function of telomerase *in vitro* and in a human disease showing telomere attrition, Dyskeratosis Congenita (DC), *in vivo* ^19^. This supports the hypothesis that senescent cell derived EC may contribute to ageing and age-related disease, but the mechanisms underlying EC export were still unclear.

We show here that EC and its plasma membrane transporter *ANKH,* but no other plasma membrane transporters are associated with fibroblast senescence and are independent of cell cycle arrest and cell size. However, the mitochondrial form of SLC25A1/mCiC, is upregulated in some senescent cells, consistent with increased flux of citrate from the mitochondria to the cytoplasm. We have established that *ANKH* was required for EC accumulation in human fibroblasts using shRNA- mediated knock down by three commercially validated inducible constructs. Furthermore, *ANKH* expression tracked with EC across a wide range of in vitro experiments. This new data together with previous work ^3^ supports the hypothesis that *ANKH* is at least partially responsible for the observed EC accumulation in senescent human fibroblasts.

However, the accumulation of EC in senescent cells and the increased expression of *ANKH* was cell type specific as neither occurred following keratinocyte or pre-adipocyte PEsen or IrrDSBsen, or indeed following keratinocyte differentiation and stratification. Nevertheless, increased expression of *ANKH* mRNA was observed in senescent astrocytes, differentiated adipocytes and myoblasts as well as fibroblasts, and the increase in plasma EC in DC where the telomeres of leukocytes and fibroblasts are short ^19^ implies that increased *ANKH* and EC are common to many cell types involved in ageing.

We previously established that the canonical function of telomerase downregulated EC in parallel with senescence ^19^ and we have now established that *ANKH* is similarly regulated. As the kinetics of EC accumulation closely parallel that of the expression of IL-6 and other SASP proteins, we investigated the pathways reported to regulate IL-6 to test whether the same pathways also regulate EC. p38MAPK and its downstream kinase, MK2/3, were required for EC and *ANKH* transcript accumulation in established senescent cells, independently of cell size and cell cycle arrest under the culture conditions we used, although the effect was slightly less marked than that on IL-6. As p38MAPK is activated in numerous types of senescent cells, including astrocytes ^61^, these results suggested EC may also accumulate in these senescent cell types. TGF-β did induce *ANKH* expression in some fibroblast types in parallel with senescence markers, though the magnitude of the effect varied. However, as the TGF-β family was previously reported to spread the senescence phenotype in a paracrine fashion ^46^, we tested whether pharmacologically inhibiting the TGF-β type 1 receptor kinase could downregulate *ANKH* in established IrrDSBsen fibroblasts. We found that this was the case, although the effect was small and not associated with the reversal of senescence markers.

Taken together our data suggests that EC accumulation via ANKH is limited to PEsen and IrrDSBsen and perhaps by TGF-β in certain cell types, although the last of these requires further investigation.

p53 which is known to restrain IL-6 in senescent pathways ^35^ also appeared to restrain EC, but this varied with cell type, was independent of senescence and only partially associated with *ANKH* expression.

In addition, the histone deacetylase inhibitor NaB induced a large increase in cell size and an increase in IL-6 as reported by others ^43^, but did not induce significant levels of EC within 4 days. Furthermore, although longer-term NaB treatments induced senescence and in some instances EC accumulation, this was independent of *ANKH* induction, and so *ANKH* does not mediate all forms of EC accumulation following senescence.

Of further significance, we found that whilst the established orchestrator of the senescence-associated inflammasome, IL-1α ^46^, induced high levels of IL-6 as reported ^40,46^ and its steroid inhibitors restrained it ^40^, these manipulations had no effect on EC or *ANKH* in established senescent and proliferating cells. This last observation suggests that EC accumulation and *ANKH* are regulated independently from IL-6 and are not merely a consequence of the inflammasome or inflammation. IL-1α increased proliferation and reduced both EC and *ANKH* expression in the absence of any marked effect on senescence markers, whilst at the same time increasing IL-6 secretion. These data further underline the differential regulation of EC and IL-6 secretion despite their similar kinetics and common regulation by the p38MAPK, MK2/3 and TGF-β pathways.

Astrocytes are the source of citrate supply in the nervous system, although no candidate plasma membrane transporter has previously been identified ^62^. Our data showing that *ANKH* is expressed by astrocytes is significant as it suggests that *ANKH* is a candidate for a citrate exporter in astrocytes. Interestingly, we showed that *ANKH* is over-expressed in senescent human astrocytes *in vitro*. Moreover, deletion of senescent astrocytes and microglia ameliorate tau-dependent pathology ^63^ and senescent oligodendrocyte precursors amyloid beta pathology ^64^ in mouse models of AD.

There was no evidence for an increase in *Ank* transcript in three mouse tissues with chronological age, despite reported evidence for EC accumulation in several mouse biofluids with age. This data might suggest that *Ank* upregulation does not mediate EC accumulation in mice ^6,7^ but is consistent with *ANKH* and EC downregulation by IL-1α in human fibroblasts and with the upregulation of mouse SASP cytokines in the aged mouse brain we examined, as in both situations inflammatory cytokines such as IL-6 are elevated. Furthermore, single cell RNA sequencing data of the aged mouse brain (Broad Institute Atlas of the Aging Mouse Brain ^65^) showed considerable heterogeneity of *Ank* expression in different cell types. High levels of expression were seen in astrocyte and oligodendrocyte clusters, low levels in microglia, endothelial and vascular smooth muscle clusters and variable expression in mature neurone clusters. This data is consistent with our new data obtained from aged microglial and neurone tissue.

The role of *ANKH* in age-related disease in humans has yet to be determined, but genome-wide analysis has associated the *ANKH* gene with a risk of developing AD ^27,28^, other forms of dementia ^29^ and type II diabetes ^30^. Furthermore, there is recent evidence linking *ANKH* downregulation in human vascular smooth muscle cells with aortic aneurism ^13^. The association of *ANKH* with AD particularly interesting given that astrocytes are thought to be the major source of citrate supply to neurones in the brain ^62^. Moreover, we have shown that *ANKH* is upregulated in PEsen and IrrDSBsen astrocytes, which in turn are instrumental for AD development in mouse models ^63^. However, we observed a downregulation of *Ank* in aged mouse brain tissue, mainly in neurons and microglial cells. Therefore, it is also possible that the link of *ANKH* to AD in humans ^27^ is also linked to its downregulation, as the associated single nucleotide polymorphism (rs112403360) is a non-coding intronic variant, so it is currently impossible to deduce how it pre-disposes to AD and the cell types in which it operates. However, cognitively healthy centenarians are enriched for the protective allele rs112403360. The protective role of the rs112403360 allele this has been suggested to be linked to its known role in bone and vascular health ^28^ and there is evidence to support this ^13,25,29,30^. However, the protective allele rs112403360 increased *ANKH* function in astrocytes this could also benefit citrate supply to the brain. Significantly, loss of function mutations in the citrate plasma membrane importer, *SLC13A5*, in humans result in neonatal epilepsy and injection of citrate into the brain can induce seizures (reviewed in ^2^), so citrate levels need to be tightly regulated, and too little citrate or too much uptake or production may be pathogenic.

ANKH itself may not be a suitable drug target due to the reported deleterious effects of its dysfunction in both mice and humans ^3^. However, the upstream negative regulators of EC and *ANKH* identified in this report such as telomerase ^66^ and MK2/3 ^67^ are established drug targets that ameliorate senescence and age-related pathologies in mice. Furthermore, the recent identification of ATP citrate lyase as a target for excessive intracellular citrate accumulation following *Ank* loss of function ^13^ presents another pharmacological avenue to exploit citrate transport and *ANKH*/*Ank* depletion.

## Conclusions and future directions

In summary, our new data identifying that the novel citrate plasma membrane exporter *ANKH* is upregulated in a variety of senescent cell types relevant to ageing and its regulation by established drug targets, informs on how countering the deleterious effects of senescent cells and telomere attrition might be addressed in the future. However, as *ANKH* and citrate may have opposing effects in humans and mice as well as in different ageing tissues, depending on dietary factors ^8–12^, further research will be necessary to exploit these observations, especially in human tissues and organoids.

### Limitations of the study

We have demonstrated a link between *ANKH* and EC upregulation in senescent human cells *in vitro* and offered a potential explanation for the downregulation of *Ank* in aged mouse brain. However, although there is extensive evidence that *ANKH*/*Ank* regulate citrate export in both human cells and mouse tissues, the knockdown achieved by shRNA in human senescent cells was incomplete and so it remains to be established whether *ANKH* is the sole transporter mediating citrate export in human senescent cells. It also remains to be demonstrated that astrocyte Ank mediates the supply of EC to neurones and that its reduction mediates neurodegenerative pathologies such as AD. This could be addressed by targeted knockout of *Ank* in astrocytes in future studies.

## Materials and Methods

### Cell culture

#### Cells

BJ cells at 16 mean population doublings (MPDs) were a generous gift from Professor Woodring Wright of Southwestern University, Houston, Texas, USA and used at between 22 and 25 MPDs. Later passage BJ cells were obtained from the American Type Culture collection and passaged until senescent (PEsen). Normal human oral fibroblast lines NHOF- 1, NHOF-2 and NHOF-7 cells were derived from explant cultures of normal oral mucosa and used at early passage between 18 and 22 MPDs and NHOF-1 was serially passaged until PEsen at between 65 and 70 MPDs ^18^. IMR90 cells at early passage were used between 1.9 and 15 MPDs after receipt. Late passage IMR90 cells were obtained from the American Type Culture collection at 28 MPDs and passaged until PEsen at between 58 and 60 MPDs.

Cells were cultured in DMEM supplemented with penicillin and streptomycin antibiotics to a concentration of 50 U/mL and L-glutamine to a concentration of 2 mM, containing 10% vol/vol fetal bovine serum at 37°C in an atmosphere of 10% CO_2_/90% air. Flasks were kept at roughly 80% confluence, and medium was replenished every 3-4 days. Once cells became more than 80% confluent, or were needed for an experiment, they were washed once with warm (37°C) PBS containing 0.02% weight/vol EDTA before being incubated for 5 minutes with PBS containing 0.1% weight/vol trypsin and 0.01% weight/vol EDTA (1mL/10cm^2^ dish). Following cell detachment, the trypsin was neutralized by the addition of serum containing media (3 mL media for every 1 mL trypsin solution), and cells were counted manually using a haemocytometer to enable calculation of the cumulative MPDs (MPDs). MPDs were used throughout the study as a measure of chronological age and were calculated using the formula: MPDs = 3.32((log10cell number yield) -(log10cell number input) ^56^.

### Mycoplasma testing

NHOF-1 was originally evaluated for mycoplasma using MycoFluor mycoplasma detection kit and found to be negative. Conditioned media both BJ and NHOF-1 cell line panels were evaluated for mycoplasma using a Lonza MycoAlert^TM^ Mycoplasma Detection Kit and found to be negative.

### Astrocyte Culture

Human astrocytes were obtained from ScienCell Research Laboratories and cultured in astrocyte growth medium (ScienCell Research Laboratories, AM, Cat. #1801) supplemented with 2% vol/vol foetal bovine serum (Cat. #0010), 1% vol/vol Astrocyte Growth Supplement (AGS, Cat. #1852) and 1% vol/vol of penicillin/streptomycin solution (Cat. #0503).

Upon thawing, the ampoules from liquid nitrogen or subculturing the astrocytes were plated on poly-L-lysine-coated culture vessels. Vessels were pre-treated by adding 2 μg/cm^2^, (10 mL of sterile water to a T-75 flask plus 15 μL of poly-L-lysine stock solution (10 mg/mL, Cat. #0413) or the equivalent thereof) to the appropriate culture vessel. The vessels were incubated at 37°C overnight and washed twice with sterile double distilled water before use. The cells were disaggregated as above when around 50% confluent and replated at around 1 x10^5^ cells per T25 flask before use in experiments. The cells were maintained in an atmosphere of 5% CO_2_/95% air.

The cmpds and the state of senescence PEsen and IrrDSBsen was analysed as indicated above.

### Adipocyte Culture

Human sub-cutaneous adipocytes were obtained from Cell Applications Inc. and cultured in pre-adipocyte growth medium (Cell Applications Inc. Cat# K811-500) supplemented with 5% vol/vol foetal bovine serum, 0.4% vol/vol endothelial cell growth supplement, 10 ng/mL recombinant epidermal growth factor, 1μg/mL hydrocortisone and 1% vol/vol and penicillin and streptomycin). The cells were disaggregated as above when around 50% confluent and replated at around 1 x10^5^ cells per T25 flask before use in experiments. The cells were maintained in an atmosphere of 5% CO2/95% air.

The cmpds and the state of senescence PEsen and IrrDSBsen was analysed as indicated above.

### Induction of adipocyte Differentiation

To induce differentiation the cells were grown plated at 4.4 x 10^5^ per cm to reach confluence and maintained in Cell Applications adipocyte differentiation medium (Sigma Aldrich Cat# 811D-250) for 15 days but were not starved as this might have affected cellular senescence and citrate metabolism.

### Oil Red staining

Adipocytes and pre-adipocytes were stained with Oil Red O using a commercial kit from Biovision (Cat# K580-42) and the intensity of staining semi-quantified by measuring the absorbance. After staining with hematoxylin and washing with distilled water, the cells were washed an additional three times with 60% isopropanol (5 min each time with gentle rocking). The Oil Red O stain was then extracted with 100% isopropanol for 5 min with gentle rocking (250 μL per well) 80% of extraction volume was used to measure. 100% isopropanol was used as a background control and subtracted from the background reading. Absorbance was read at 492 nm using a CLARIOstar plate reader (BMG LABTECH).

### The induction of senescence by ionizing radiation (IrrDSBsen)

Fibroblasts, keratinocytes astrocytes and adipocytes were irradiated in suspension with 10Gy of γrays from a Cs source at a dose rate of 1.4 Gy/min ^18^ or in later experiments with 10 Gy of X rays at a dose rate of of 3.6 Gy/min to induce irreparable DNA double strand breaks. The cells were left between 0 and 20 days in culture before analysis. Where indicated the keratinocytes were irradiated with γ rays in situ following removal of the 3T3 feeder cells with EDTA and vigorous pipetting as described^68^. 3T3 feeder cells were irradiated with 60 Gy of either γ- or X rays as indicated above ^68^.

### Experimental design and drug treatment

Cells were seeded in such a way as to ensure that the density of both senescent and growing controls were similar although to induce short-term confluence higher numbers of cells were seeded to ensure short-term reversible growth arrest, and the numbers of senescent cells seeded to match the level of confluence where indicated.

Cells were seeded for 3 to 5 days prior to medium changing for 16 h before lysis or collection of conditioned media. Except for the long-term NaB experiments all drugs, growth factors, cytokines and inhibitors were added for the indicated time and during the final 16 h prior to analysis. Controls were either culture medium with no additives or vehicle controls (ethanol or DMSO at 0.1% vol/vol)

For shRNA induction, cells were cultured in the indicated level of DOX for one week prior to setting up the experiment and for the entire experiment. Non-induced cells and DOX-induced cells expressing the non-targeting vectors served as controls.

### Characterisation of senescent cells

The cellular senescence status of the cells was confirmed through senescence-associated beta galactosidase (SA-βGal) activity using commercial kits (Biovision K-320-250 and later Generon senescence activity assay kit Cat# KA6035), which was measured again at the time at which the experiments were conducted. Briefly, the kits were brought to room temperature for 20 minutes. Following this, the cells were washed twice with calcium and magnesium-free phosphate buffered saline (PBS) fixed for 10 min and washed twice more before adding the SA-βGal reagent. The plates were then sealed with parafilm to prevent dehydration, covered in foil to protect from light and incubated at 37°C in a hot room or hotbox in an atmosphere of air (0.03% CO_2_. Late passage IMR90 cells were used as a positive control.

### Western blotting

The following antibodies were used: pmCiC and mCiC (SLC25A1) rabbit polyclonal antibodies ^69^ were custom-made by GenScript Inc., ANKH (Rabbit VivaSystems, cat#OAAB06341), (MCT1 (rabbit Santa Cruz cat#sc-365501), MCT4 (goat Santa Cruz cat# sc-14930), Actin C-4 (mouse Santa Cruz cat# sc-47778), ASCT2 (rabbit Cell Signaling cat# 5345).

Briefly, cell extraction was performed to isolate proteins from confluent cultures. Cells were washed with PBS and extracted with 2× sodium dodecyl sulphate (SDS) Laemmli sample buffer (0.5 M Tris-Cl pH 6.8, 4% SDS, 20% glycerol; 10% (v/v) 2-mercaptoethanol was added after protein assay) immediately. DC Protein Assay (Bio-Rad, Hercules, CA, USA) was used to determine the protein concentration for each sample. use approx. 25 µg (10-30 µg) of protein was prepared with 5µL LaemmLi Buffer, centrifuged at 15,000 rpm (20,000 g), boiled at 95°C for 5 min, spin down, centrifuged again and stored on ice.

The proteins were separated on 10% SDS polyacrylamide gels at 100V for 90 to 150 min, transferred to nitrocellulose filters (Whatman) in SDS running buffer and methanol at 100 V for 50-60 min. After blotting, the membrane was placed in 5% w/v-milk powder in Tris-buffered saline containing 10% w/v SDS, glycine, 0.1% vol/vol Tween 20 (TBS-T) on shaker for 1 h. Following blocking, the primary antibodies were diluted 1:1000 in TBS-T and 5% w/v/ milk powder and incubated at 4°C overnight. The following day the membranes were washed twice for 15 min each in TBS-T with agitation. The membranes were then incubated with the appropriate secondary antibody at a dilution of 1:2500 in TBS-T and 5% w/v/ milk powder for 1 h at room temperature. The membranes were washed twice for 15 min each in TBS-T with agitation before incubating with freshly prepared ECL for 2 min and developing with X-ray film developer in the dark and marking the molecular weight markers on the film. The membranes were washed with TBS-T as above before repeating with the loading control.

### Collection of conditioned medium

Conditioned medium was collected after 24 h from the cells, centrifuged at 800 x g for 2 min, the supernatant removed and centrifuged again at 15,000 rpm for 2 min and the final supernatant snap frozen on an ethanol-dry ice bath for 15 min before storage at −80°C. Unconditioned medium was also prepared identically.

### Targeted measurement of extracellular citrate by gas chromatography/mass spectroscopy (GC-MS)

Deuterated citrate (Citrate-d_4_) was added to each sample to a final concentration of 0.1 mM as an internal standard. Metabolites were then extracted using cold methanol before being dried under vacuum desiccation. The samples were re-suspended in anhydrous pyridine containing the derivatisation agents methoxyamine hydrochloride followed by N-Methyl-N-trimethylsilyltrifluoroacetamide with 1% 2,2,2-Trifluoro-N-methyl-N-(trimethylsilyl)-acetamide and Chlorotrimethylsilane (MSTFA+1%TMCS). GCMS was performed in pulsed split less mode on a Hewlett Packard HP6890 series GC system with Agilent 6890 series injector and a 30 m x 250 µm capillary column (Agilent, model number 19091s-433HP5MS) using a flow rate of 1 mL/minute, and a Hewlett Packard 5973 mass selective detector. The acquisition was conducted in selective ion monitoring mode, with the m/z values 273, 347, 375 and 465 for citrate, and 276, 350, 378 and 469 for citrate-d_4_. The dwell time for all these ions was 50ms.

### Targeted measurement of extracellular citrate by liquid chromatography/mass spectroscopy (LC-MS)

We measured citrate using a targeted LC-MS-MS method with a mixed-mode UPLC chromatographic separation step for retention and discrimination of organic acids, including excellent discrimination of citrate from isocitrate. The media samples were diluted 10fold with cold methanol and then diluted again 5fold with cold methanol containing an internal standard (1 µg / mL L-Phenyl-d_5_-alanine (Sigma-Aldrich, 615870)). The mixture was vortexed at 6 °C for 20 minutes, and then centrifuged (17,000 g, 10 minutes, 4 °C). 490 µL aliquots of the supernatant were dried under reduced pressure and stored at −80 °C until analysis.

We then resuspended the dried samples in 50 µL of water containing the stable-isotope-labelled (SIL) standard ^13^C_3_-citrate (Cambridge Isotope Laboratories, CLM-9876), and measured citrate essentially as described by Smith et al ^70^ with modification of the LC gradient. Specifically, the initial binary elution gradient was 0% B; 0.1 min: 0% B; 8.1 min: 25% B; 11.1 min: 95% B; 12.1 min: 95% B; 13 min: 0% B and 15 min: 0% B. The separation was performed using an Acquity Premier CSH Phenyl-Hexyl column (2.1 × 100 mm, 1.7 μm). The analysis was performed on a XEVO TQ-S (Waters UK) using negative ionization mode, with settings of capillary voltage, 1.5 kV; source offset, 50 V; desolvation temperature 500°C; source temperature, 150°C; desolvation gas flow, 1000 L/h; cone gas flow, 150 L/h; nebuliser gas 7.0 bar; collision gas, 0.15 mL/min. The transitions used in MS were detailed as follow: Citrate Q1 191.0197 *m/z* → Q3 111.1000 *m/z*, collision energy 12 V; Q1 191.0197 *m/z* → Q3 87.1000 m/z, collision energy 18 V and ^13^C_3_ citrate Q1 194.0197 *m/z* → Q3 113.1000 *m/z*, collision energy 12 V; Q1 191.0197 m/z → Q3 87.1000 m/z, collision energy 18 V. All the transitions set a dwell time of 0.008 second and a cone voltage of 45 V.

We quantitated citrate against an authentic standard (Supelco) using a 6-point calibration curve (from 0.025 – 1 mg/l, run immediately before and after the samples), together with the SIL ratio. The results were checked against both water and method blanks.

### Reverse transcription and quantitative PCR (RT-qPCR)

Cultured cells were lysed in lysis buffer and mRNA extraction was performed as per manufacturer’s instruction using Dynabeads mRNA Direct kit #61012 (ThermoFisher, UK). Purified mRNA sample was then combined with qPCRBIO SyGreen 1-Step Go (PCR Biosystems, PB25.31- 12) master mix and qPCR primers for one-step relative quantification of target genes and 2 reference genes using a 384-well format LightCycler 480 qPCR system (Roche) based on our previously published protocols ^71–74^ which are MIQE compliant ^75^. Briefly, thermocycling begins with 45°C for 10 min for reverse transcription followed by 95°C for 30 s prior to 45 cycles of amplification at 95°C for 5 s, 60°C for 5 s, 72°C for 5 s, 78°C for 1 s (data acquisition). A ‘touch-down’ annealing temperature intervention (66°C starting temperature with a stepwise reduction of 0.6°C/cycle; 8 cycles) was introduced prior to the amplification step to maximise primer specificity. Melting analysis (95°C for 30 s, 75°C for 30 s, 75-99°C at a ramp rate of 0.57°C/s) was performed at the end of qPCR amplification to validate single product amplification in each well. Relative quantification of mRNA transcripts was calculated based on an objective method using the second derivative maximum algorithm ^76^ (Roche). All target genes were normalised to two reference genes (HPRT1 and B2M) validated previously ^72^ to be amongst the most stable reference genes across a wide variety of primary human epithelial cells, dysplastic and squamous carcinoma cell lines, using the GeNorm algorithm ^77^. Relative expression data were then exported into Microsoft Excel for statistical data analysis. No template controls (NTC) were prepared by omitting tissue sample during RNA purification and eluates were used as NTCs for each qPCR run.

### Human primer sequences

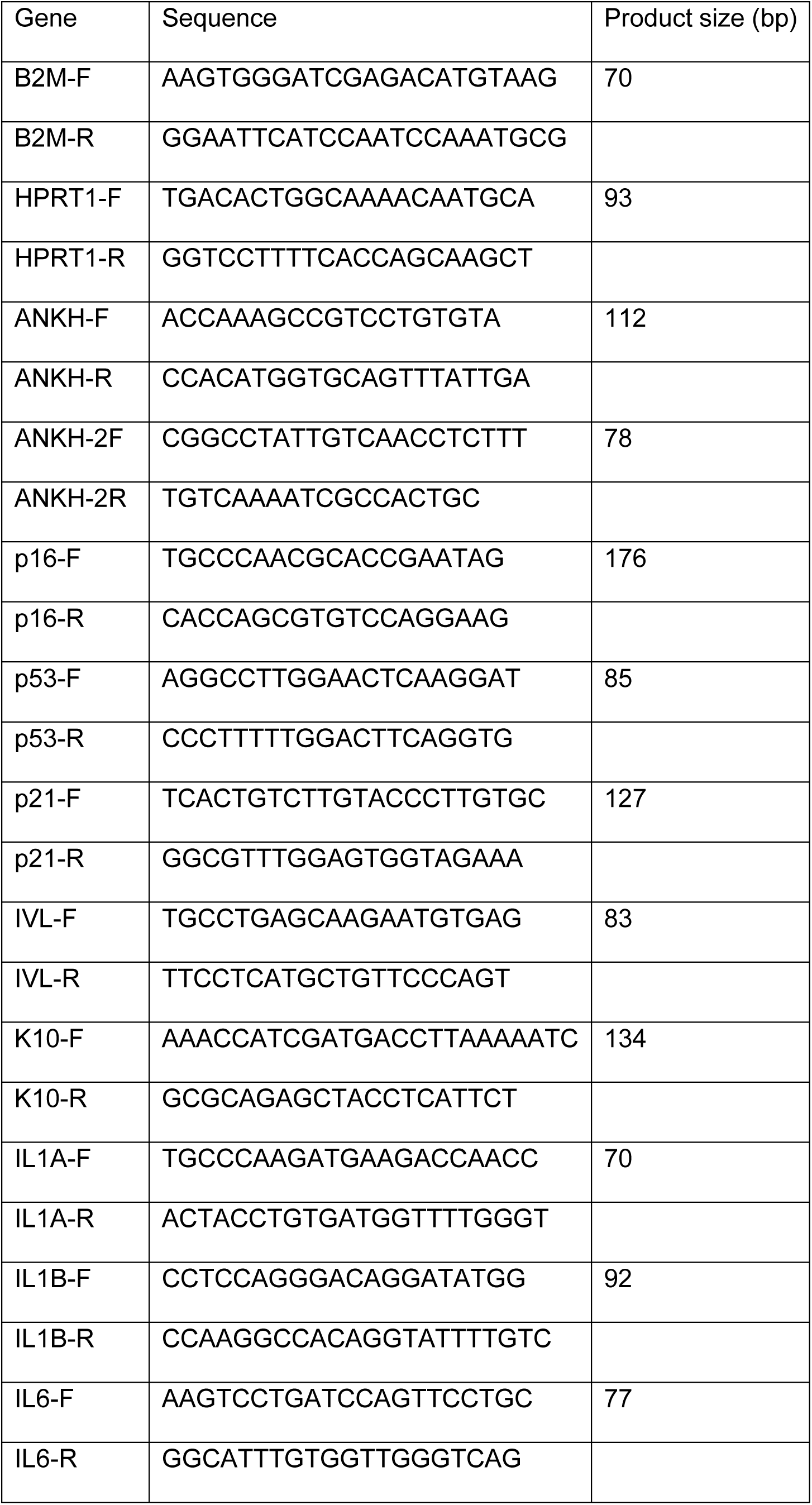
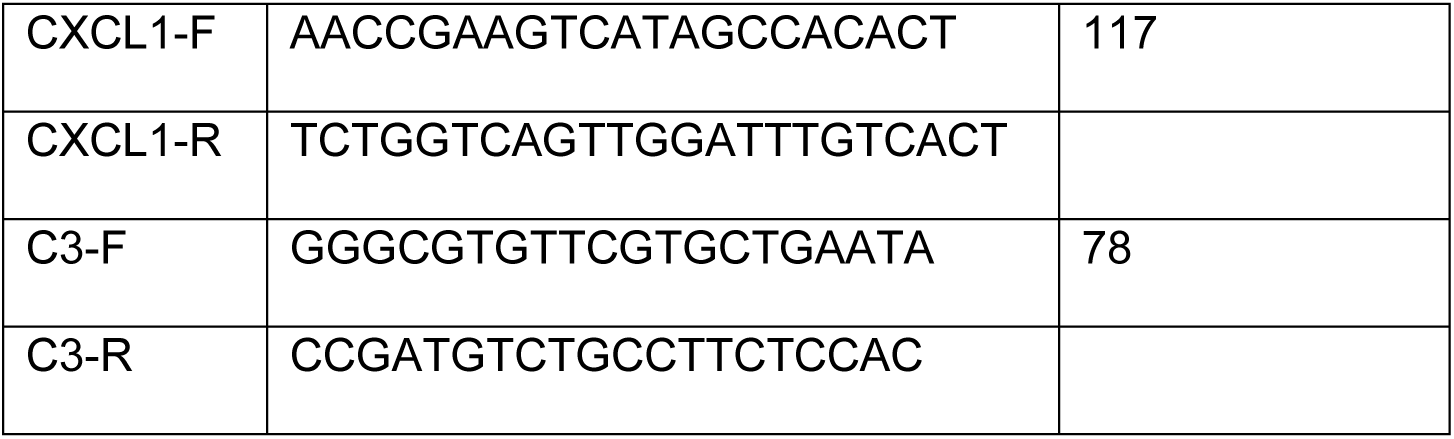

### The knock down of target genes by conditionally expressed short hairpin RNA (shRNA) constructs

We used pre-prepared lentiviral particles to infect the indicted human cells from Horizon Discovery (SMARTvector Inducible Lentiviral shRNAs).

First, we identified the promoter, which gave the best induction of shRNA based on either Green Fluorescent Protein (GFP) or Red Fluorescent Protein (RFP). We did this by utilizing the SMARTchoice inducible non-targeting control 4-Pack (Cat #VSC6847) and following the manufacturer’s instructions. Based on these pilot experiments we identified the mouse cytomegalovirus promoter as giving the best induction and so proceeded to work with constructs on this backbone.

The indicated cells were infected with lentiviral particles in transduction medium (serum-free Dulbecco’s Modified Eagle Medium (DMEM) High Glucose without L-Glut or Sodium Pyruvate at a multiplicity of infection of 0.3 virus particles per cell. The following day the cells were changed into regular culture medium and on day 3 trypsinised and plated at 1 x10^5^ cells per T75 flask. After a further incubation time of 4 days 1 µg/mL puromycin was added until all the mock-infected cells were dead and then the puromycin was removed prior to expansion of the cell populations and cryopreservation. We used this protocol because culturing normal cells in selecting agents can reduce replicative lifespan.

The dose of doxycyclin hyclate (DOX; Thermo Scientific, Cat #ICN19895510) was optimised by measuring GFP or RFP fluorescence following 4 days of induction. The toxicity of DOX was estimated in parallel by the addition of MTT for 1 hour in serum-free DMEM followed by dissolving the cells in Dimethyl Sulphoxide (DMSO) and measurement of the absorbance at a wavelength of 570nm. The level of knockdown by each shRNA was monitored by qPCR as indicated below. To achieve maximum levels of knock down the induced cells were purged of low expressing cells by flow sorting for cells highly positive for GFP or RFP after induction with DOX using non-induced cells as background. We chose 3 shRNAs validated by Horizon Discovery for both *ANKH* and *TP53*.

The sequences of each shRNA targeting sequence are given below.

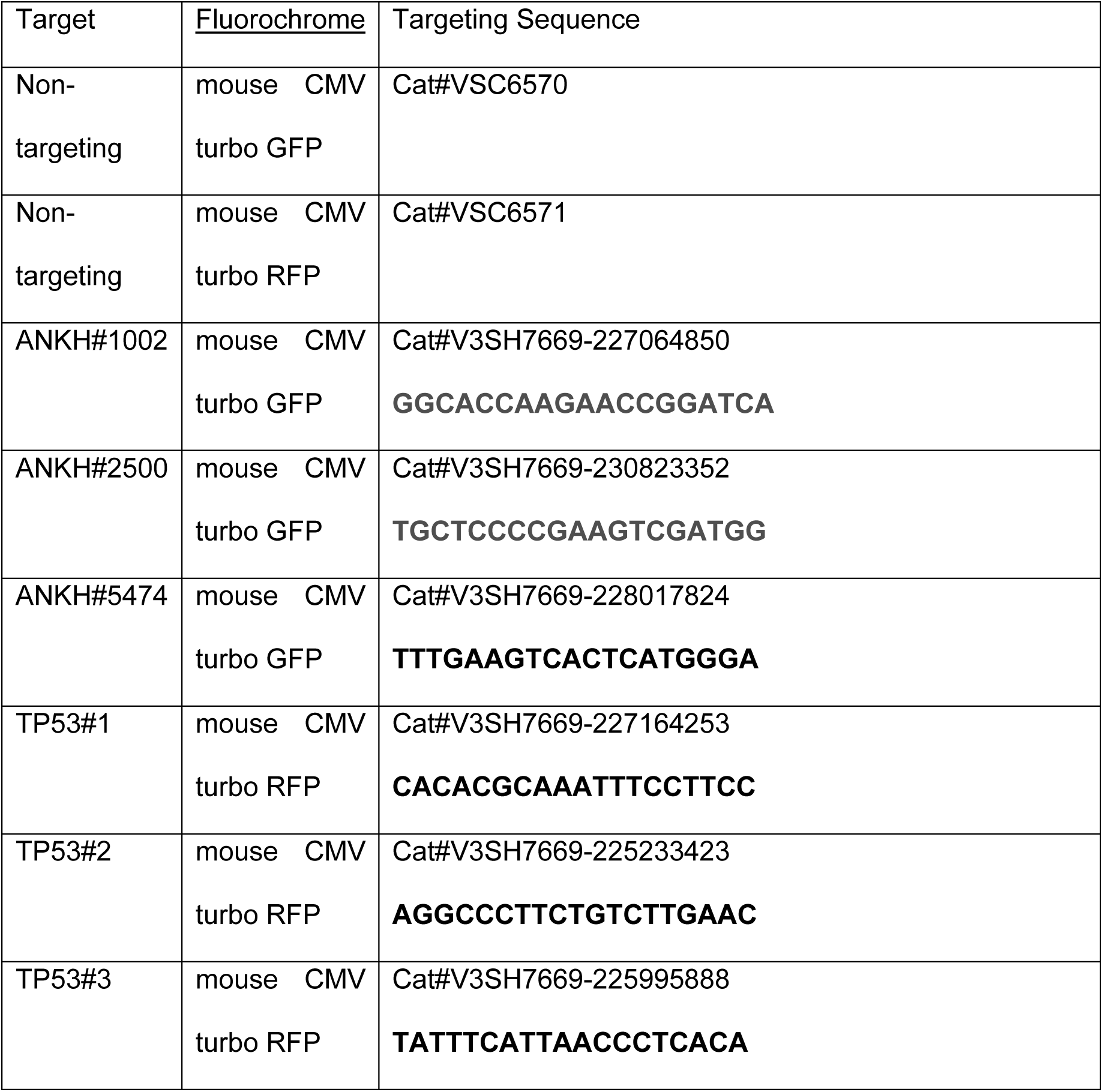

### Enzyme-Linked Immunosorbent Assay (ELISA)

To measure IL-6 in conditioned media a sandwich ELISA method (Quantikine® ELISA Immunoassay, R&D Systems, Abingdon, UK) was employed following the manufacturer’s protocol. The detection limit for IL-6 was 3.13-300 pg/mL.

### Mouse experiments

The details and characterisation of the mouse liver and kidney tissues have been published previously ^58^.

#### Animals for aged brain analysis

The available strain for studying the role of ageing was C57BL/6, adult (8-10 weeks old) and aged (17 months old) male C57BL/6 mice, weighing 20-24 g at the start of the study (Janvier Laboratory, France). The Animal Welfare and Ethical Review Body, at Queen Mary University of London and the United Kingdom Home Office, following the EU Directive 2010/63/EU, approved all animal procedures.

#### Animal culling and sample collection

In order to collect tissue, animals were deeply anaesthetised using sodium pentobarbital (50 mg/kg) administered intraperitoneally (i.p.) in a volume of 0.5 mL/kg. Brain tissue and blood samples were collected from all animals. Animals were either perfused with phosphate-buffered saline (PBS) followed by 4% paraformaldehyde (PFA) for the analysis of the tissue by immunostaining, or with PBS only for the cytokine analysis that required fresh frozen tissue. Blood samples were collected in lithium heparin tubes and then centrifuged at 10,000 g for plasma separation. Plasma samples were stored at −80°C for future analysis.

#### Immunostaining

All tissue staining was performed on 8 μm coronal brain sections cut between bregma −1.9 mm and −2.1 mm. Sections were deparaffinised in two washes xylene (10 min each) and then hydrated in a series of ethanol (EtOH) baths (100% EtOH, 100% EtOH, 90% EtOH and 70% EtOH, 5 min each) and two washes of PBS (5 min each). Then sections were treated with a heated antigen retrieval solution containing 10 mM citric buffer (pH 6.0) for 10 min in a microwave. After that, sections were blocked with 8% bovine serum albumin (BSA), 0.5% Tween-20 and 0.1% Triton X-100 in PBS overnight in a humidified chamber at 4°C. On the next day, sections were washed twice in 0.5% Tween-20 and 0.1% Triton X-100 in PBS (PBS-TT) and then incubated with primary antibodies (Table 2.2) overnight in a humidified chamber at 4°C. Markers were visualised using secondary goat-raised antibodies, which were labelled with either Alexa 488 or Alexa 555 (Invitrogen, United States; 1:250), and nuclei were visualised using Hoechst 33342 stain (Tocris Bioscience, United Kingdom; 1:500). Slides were mounted and cover-slipped using Vectashield fluorescent mounting medium (Vector Laboratories, United States).

#### Antibodies used

| Marker | Antibody full name | Species | Company | Product ID | Dilution |
| --- | --- | --- | --- | --- | --- |
| Anti-p53 | Anti-tumour protein P53 | Rabbit | Insight Biotechnology, United States | GTX102965 | 1:150 |
| Anti-p21<sup>Waf</sup> | Anti-cyclin-dependent kinase inhibitor 1 | Rabbit | Invitrogen, United States | 14-6715-81 | 1:150 |
| Anti-p16<sup>Ink4A</sup> | Anti-cyclin-dependent kinase inhibitor 2A | Rat | Abcam, United Kingdom | ab241543 | 1:150 |
| Anti-p19<sup>Arf</sup> | Anti-ARF tumour suppressor | Rat | Novus Biological, United States | NB200-169 | 1:150 |
| Anti-γH2AX | Anti-phosphorylated H2A histone family member X | Mouse | Merck KGaA, Germany | 05-636 | 1:500 |
| Anti-53BP1 | Anti-tumour suppressor p53-binding protein 1 | Rabbit | Cambridge Bioscience, United Kingdom | A300-273A | 1:1600 |

#### ImmunoFISH telomere associated foci (TAF) staining

ImmunoFISH staining was used to detect senescent cells. First, sections were deparaffinised in two washes of xylene (10 min each), washed in a gradient of ethanol (100% EtOH, 100% EtOH, 90% EtOH and 70% EtOH, 5 min each) and then in two changes of PBS (5 min each). After that, sections were treated with ice-cold (−20°C) 70% ethanol for 20 min then washed three times in PBS for 15 min. Next, sections were blocked with 8% BSA (Sigma-Aldrich, Saint Louis, United States), 0.5% Tween-20 (Sigma-Aldrich, Poole Dorset, United Kingdom) and 0.1% Triton X-100 (Sigma-Aldrich, Poole Dorset, United Kingdom) in PBS overnight at 4°C in a humidified chamber. On the next day, sections were washed with PBS- TT for 5 min, then one or two primary antibodies were added and then left overnight at 4°C in a humidified chamber. One of them was an anti-phospho-histone γH2A.X antibody (Cell Signalling Technology, Danvers, United State; 1:250) and the second was anti-NeuN (Merck Millipore, Burlington, United States; 1:250). On day 3, sections were washed in PBS-TT for 5 min twice then incubated with the secondary antibodies: biotinylated anti-rabbit IgG (Vector Laboratories, Burlingame, United States; 1:250) and Alexa Fluor 647 (Far red; Invitrogen, Carlsbad, United States; 1:250) for 1 h in a humidified chamber. After that, sections were washed in PBS-TT for 5 min three times, then incubated with DSC-Fluorescein (Vector Laboratories, Peterborough, United Kingdom, 1:500 in PBS) for 20 min. Later, sections were washed twice with PBS-TT, twice with PBS for 5 min, then treated with 4% PFA in PBS for 20 min. Next, sections were washed 3 times with PBS for 5 min followed by a dehydrating step using an ice-cold gradient of ethanol (70% EtOH, 90% EtOH, and 100% EtOH, 3 min each). Sections were air-dried and then treated with a hybridisation mix which contained 0.5 μg/mL (C3TA2)3-Cy3-labeled peptide nucleic acid (PNA) telomeric probe, 70% formamide, 12 mM Tris-HCl (pH 7.4), 5 mM KCl, 1 mM MgCl_2_, 0.001% Triton X-100, and 2.5 mg/mL acetylated BSA. Sections were coverslipped and then incubated in the oven (82°C) for 10 min. The hybridisation process was continued in a humidified chamber for two hours. Finally, sections were washed in 70% formamide/2xSSC for 10 min, twice in 2xSSC for 10 min, twice in PBS for 10 min then another change of PBS for 30 min. Slides were mounted using Vectashield fluorescent mounting medium with 4′,6-diamidino-2-phenylindole (DAPI) (Vector Laboratories, Burlingame, United States).

#### Quantification of IF and TAF signals

Four coronal sections were used per animal for each antibody staining; the sections were situated along the rostral-caudal axis, with a spacing of 0.1 mm between sections. Images were captured at x40 magnification using the In Cell Analyser 2200 (INCA2200) System (Cytiva, Marlborough, United States) for the markers: 8-oxo-dG, p53, p19^ARF^, p21^WAF^, and p16^INK4A^. For the γH2AX.53BP1 signal overlap and TAF staining, fields used for the quantification were taken at x60 magnification using the In Cell Analyser 6000 (INCA6000) Confocal System (Cytiva, Marlborough, United States). A minimum number of 200 fields of view (FoV) were captured per animal covering all 4 sections, with a total of nuclei counts of at least 3 x 104 nuclei.

All quantitative analyses were conducted using the In Cell Developer Toolbox v1.9.2 (Cytiva, Marlborough, United States) and dedicated custom-made protocols were written for each marker evaluated, using the build-in parameters selection tools. A cell was considered positive for a marker when the fluorescent signal co-localised with a nucleus by at least a 95% overlap. Results were expressed as a percentage of the total nuclei detected.

### Cytokine analysis

#### Brain samples

Snap-frozen brain tissue was used for cytokine analysis. 25 mg tissue was used from each sample, and the tissue was cut into small pieces on ice. The tissue fragments were suspended in lysate buffer (RIPA Buffer containing protease/phosphatase inhibitors) and then crushed with Pellet Pestle® (Sigma-Aldrich, United Kingdom). After that, samples were placed on rotating wheels and incubated for 20 min at 4°C. Next, samples were sonicated 3 times, 15 to 20 sec each, followed by centrifugation for 20 min at 10,000 g (4°C). Then, the supernatant was collected from each sample and used for the bicinchoninic acid (BCA) assay to measure protein concentration and subsequently cytokine concentrations.

For the BCA assay, the Pierce BCA protein assay kit (Invitrogen, Life Technologies Ltd, United Kingdom) was used to measure the concentration of protein extracted from brain tissue and protocol followed manufacturer instruction. Samples were diluted with the supplied MSD solution, Diluent 41, to reach a concentration of 2.5 mg/µL which was the recommended protein concentration.

#### Plasma samples

Thawed plasma samples were diluted with the MSD supplied solution, Diluent 41, with a ratio of 1:3 before being used in the assay.

##### Running the experiment and data processing

Cytokine levels were measured in brain tissue and plasma samples using the V-PLEX proinflammatory panel 1 mouse kit, catalogue No. K15048D-1 (Meso Scale Diagnostics (MSD), United States), protocol followed manufacturer instructions and prepared plates were analyzed using MESO QuickPlex SQ 120MM (MSD, United States).

#### Mouse qPCR and primer sequences

The qPCR was carried out for the mouse Ank gene in the same way as for the human ANKH gene. The mouse primers are shown below.

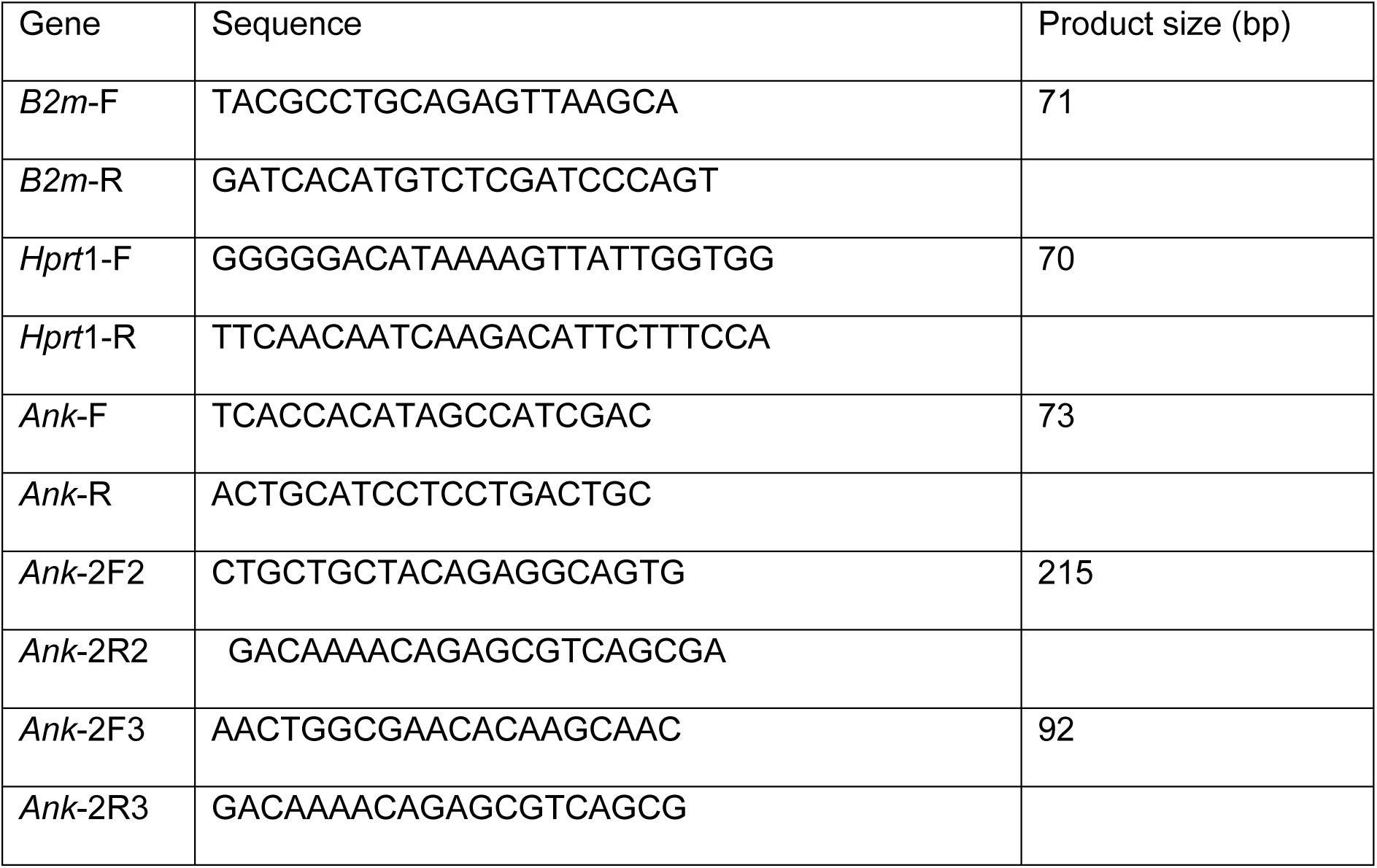

### Statistical methods

Human and mouse data where the numbers of repeats were the same were analysed by the Student’s unpaired t-test. Data sets containing more than one variable were also analysed by one-way or ordinary two-way ANOVA to confirm the results of the t-test. Where the repeat numbers were uneven human data were analysed by the Welch’s T test and the mouse data was analysed using the Mann-Whitney Test, the significance level was set at p < 0.05.

## Author Contributions

Emma Naomi James and Mark Bennett performed the gas chromatography/mass spectroscopy analysis; Yufeng Li and Jacob Bundy performed the liquid chromatography/ mass spectroscopy analysis; Muy Teck The performed the multiplex quantitative PCR analysis; Zahra Falah Hassan Al-Khateeb and Adina Michael-Titus provided the mouse brain RNA and senescence data; Ana O’Loghlen provided the previously characterised mouse liver and kidney RNA samples; Christine Wagner performed the western blots for metabolite transporters in human cell extracts^1^, Lee Peng Karen-Ng and Linnea Synchyshyn performed the cytokine analysis; Terry Roberts performed the TERT qPCR and telomerase activity assays, Amy Lewis and Andrew Silver performed the In Cell analysis of the sodium butyrate experiments; Eric Kenneth Parkinson performed al the human in vitro work and wrote the first draft of the manuscript; Maria Mycielska and Eric Kenneth Parkinson conceived the study and wrote the first draft of the manuscript.

## Acknowledgements

The authors wish to thank Cleo Bishop and Emily Anne O’Sullivan for advice regarding knockdown using Horizon Discovery shRNAs, Gary Warnes for performing the cell sorting and Simon McArthur for advice on markers of brain ageing. We also wish to thank John Sedivy of Brown University, Rhode Island for the gift of the Loxo26 cell lines.

## Funding

The study was funded in part by the Dunhill Medical Trust (grant number R452/1115) and Barts and the London Charity (grant number MRD&U0004). Karen-Ng Lee Peng received a Ph.D. scholarship (Hadiah Latihan Persekutuan) from the Malaysian Ministry of Education. Ana O’Loghlen received funding from Barts Charity Grants (G-002158) and Project I+D+i PID2021-125656OB-I00, financed by MCIN/ AEI /10.13039/501100011033/FEDER, UE and Project SenesceX-CM P2022/BMD-7393.

## Conflict of interest statement

Maria Mycielska is a co-inventor on the Patent Application no. EP15767532.3 and US2020/408741 (status patent pending) and US2017/0241981 (patent issued) “The plasma membrane citrate transporter for use in the diagnosis and treatment of cancer” owned by the University Hospital Regensburg. The other authors declare no conflict of interest.

## Data availability statement

All data sets used in this study will be provided on request by the authors.

## Notes

### Summary of Updates

The introduction has been reduced in size and both the introduction and discussion have now been focused more on the ANKH/Ank transporter specifically. The number of repeats have now been made much clearer to avoid confusion about n = 2 for example by explaining this better in the figure legends and referring to the supplementary information. We have additionally analysed some datasets with multiple variables with two-way ANOVA. A mouse database has been accessed and discussed in relation to our own findings. The validation of the shRNAs has now been explained. Our conclusions have not changed.

