## Supplementary Information for "Membrane transporter Progressive Ankylosis Protein Homolog (*ANKH*/*Ank*) partially mediates senescence-derived extracellular citrate and is regulated by DNA damage, inflammation and ageing"

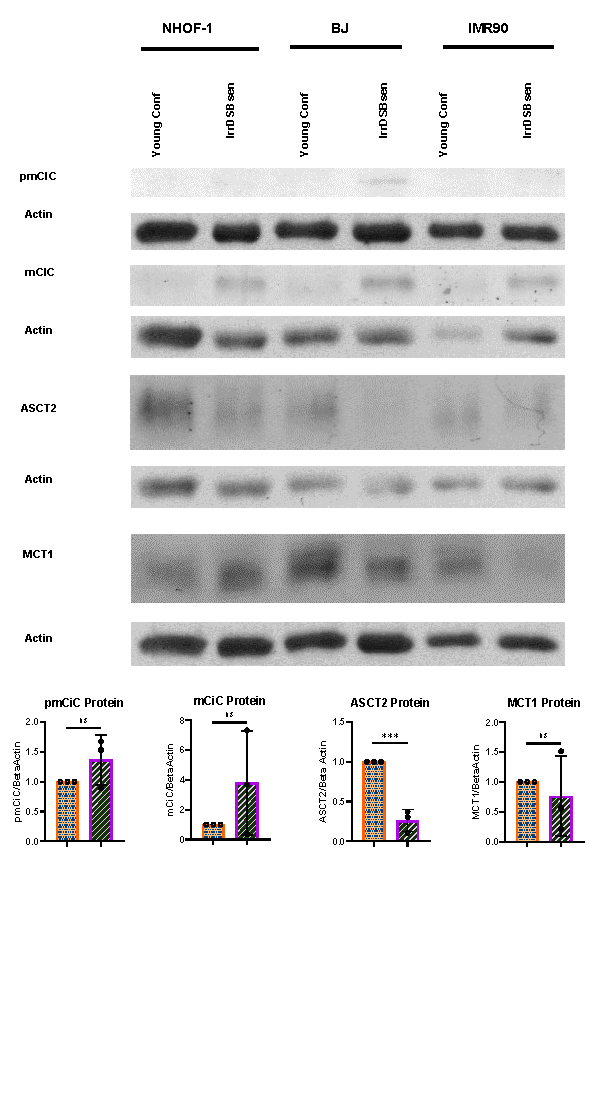
**Supplementary Figure 1 The expression of other metabolite transporters in confluent and IrrDSBsen human fibroblasts.**

A. Shows western blots of mCiC, pmCiC, MCT1 and ASCT2 proteins in 3 lines of quiescent confluent (Conf) young human fibroblasts and the same lines induced to senesce by ionising radiation (IrrDSBsen). Beta actin was used as a loading control.

B. Mean Image J quantitation of the blots in A. showing the levels of the transporter signals versus beta actin in all conditions after background subtraction.

Orange stippled bars = confluent cells; purple left-right diagonally striped bars = IrrDSBsen cells. *** = P < 0.001; ns = not significant. The results are means +/- standard deviation. N = 3.

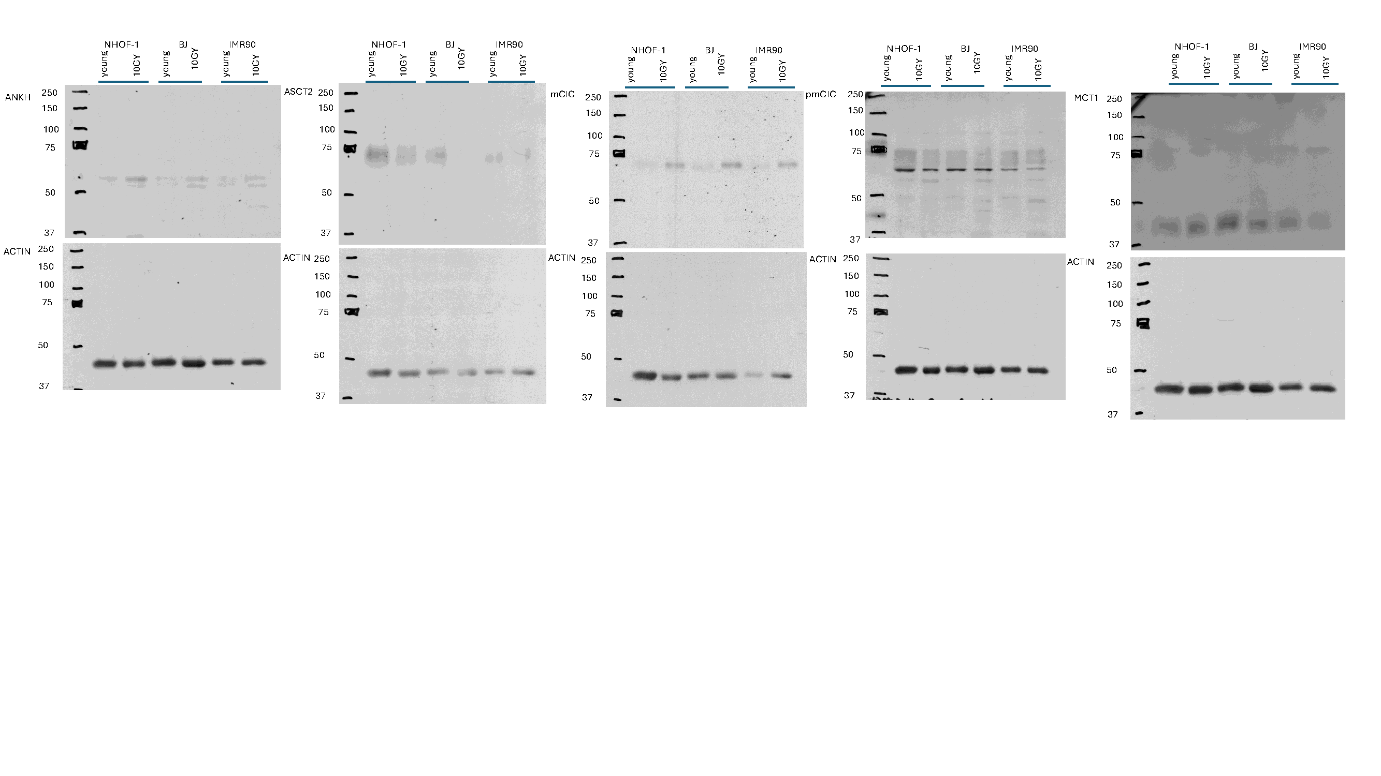

**Supplementary Figure 2 The expression of other metabolite transporters in confluent and IrrDSBsen human fibroblasts showing the full-length western blots of the cropped blots shown in Figure 1 and Supplementary Figure 1.**

The figure shows the full-length western blots of ANKH, mCiC, pmCiC, MCT1 and ASCT2 proteins in 3 lines of quiescent confluent (Conf) young human fibroblasts and the same lines induced to senesce by ionising radiation (IrrDSBsen). Beta actin was used as a loading control.

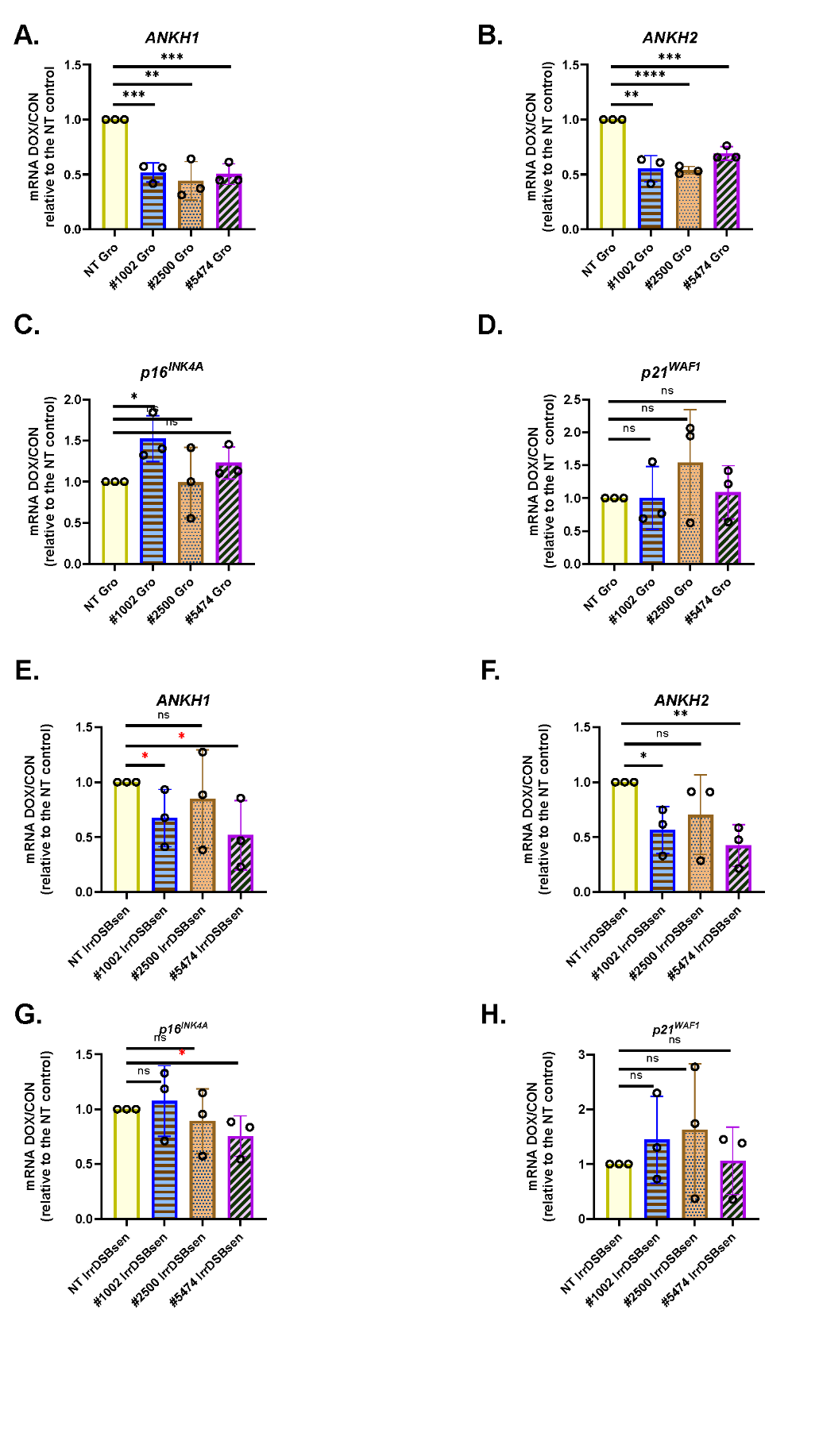

**Supplementary Figure 3 The effect of the different *ANKH* shRNAs on the expression of *ANKH*, p16^INK4A^ and p21^WAF^ in growing (A.-D.) and IrrDSBsen NHOF-1 fibroblasts (E.-I.).**

A. and **E.** *ANKH1*
B. and **F.** *ANKH2*
C. and **G.** *p16^INK4A^*
D. and **H.** *p21^WAF^*

Plain yellow bars = NT control; blue horizontal striped bars = shRNA #1002; orange stippled bars = shRNA #2500; purple left-right diagonally striped bars = shRNA #5474.

* = P< 0.05; ** = P < 0.01; *** = P < 0.001; * P > 0.05; < 0.1; ns = not significant.

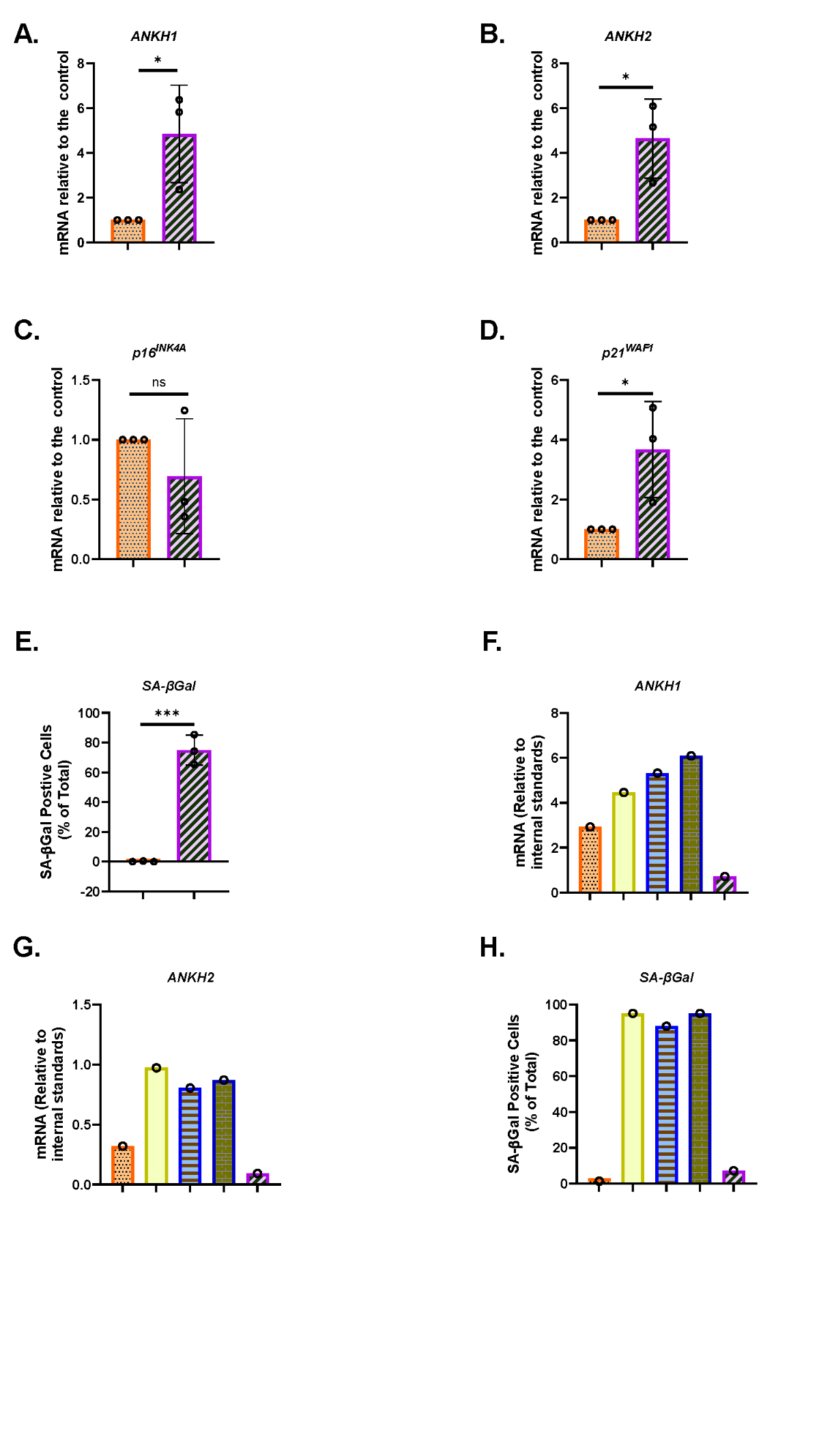
**Supplementary Figure 4 *ANKH* transcript is upregulated in PEsen human fibroblasts and downregulated by telomerase.**

A. The expression of *ANKH* mRNA in growing and PEsen BJ cells using primer set 1.

B. As in A. but the results of primer set 2 in the same experiments

C. As in A. but the expression of the senescence marker p16^INK4A^in the same experiments.

D. As in A. but the expression of the senescence marker p21^WAF1^ in the same experiments.

E. As in A. but showing SA-β Gal expression.

A-E. Orange stippled bars = growing cells; purple left-right diagonally striped bars = PEsen cells. * = P< 0.05; ** = P < 0.01; *** = P < 0.001; * trend for significance; ns = not significant. The results are means +/- standard deviation. N = 3.

F. The expression of *ANKH* mRNA in BJ PURO early passage, PURO and TERT-HA, DN TERT and TERT all at late passage using primer set 1.

G. As in F. but the results of primer set 2 in the same experiments

H. As in F. but showing SA-β Gal expression.

F-H Orange stippled bars = PURO early; plain yellow bars PURO late; blue horizontal striped bars TERT-HA, dark blue brick pattern bars DN TERT and purple left-right diagonally striped bars TERT.

The data in F-H is from a single experiment.

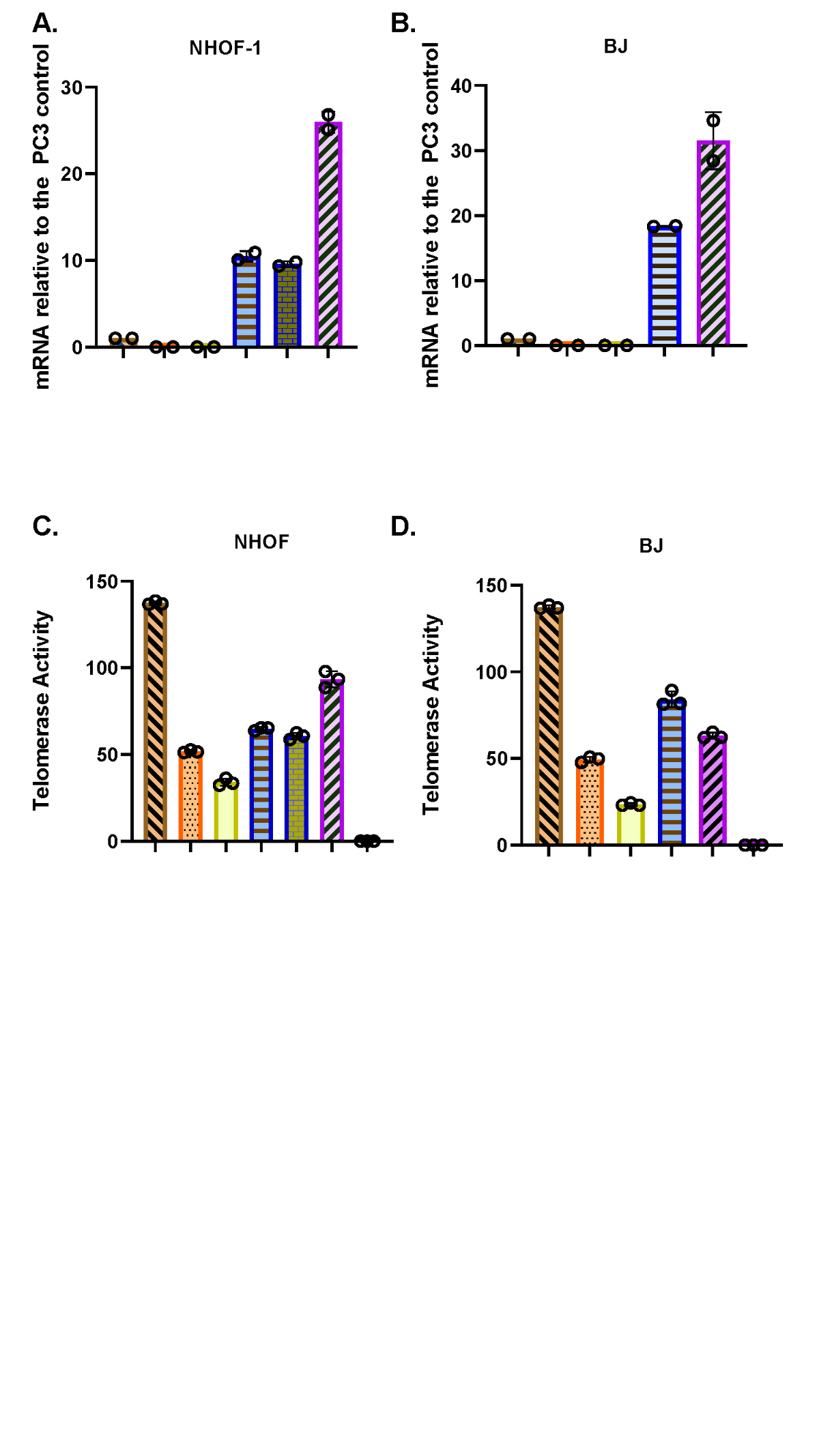
**Supplementary Figure 5 *TERT* mRNA and telomerase activity are expressed in late passage NHOF-1 and BJ fibroblasts.**

A. *TERT* mRNA expression in NHOF-1 fibroblasts after more than 50 MPDs.

B. BJ mRNA expression in NHOF-1 fibroblasts after more than 50 MPDs.

C. *TERT* telomerase activity in NHOF-1 fibroblasts after more than 50 MPDs.

D. BJ telomerase activity in NHOF-1 fibroblasts after more than 50 MPDs.

A-D. Brown right-left diagonally striped bars = PC3 positive control; Orange stippled bars = NHOF-1 and BJ young empty vector (PURO) controls; Plain yellow bars = NHOF-1 and BJ old empty vector (PURO) controls; blue horizonal striped bars = NHOF-1 and BJ TERT-HA; dark blue brick pattern bars = NHOF-1 DNTERT; purple left-right diagonally striped bars = NHOF-1 and BJ TERT; C. and D. black chequered bars = PC3 Heat-treated CHAPs extract.

No activity was detected in any heat-treated CHAPS extract.

A. and B. The means of two independent determinations run in triplicate +/- standard deviation. C. and D. The means of one determination run in triplicate +/- standard deviation.

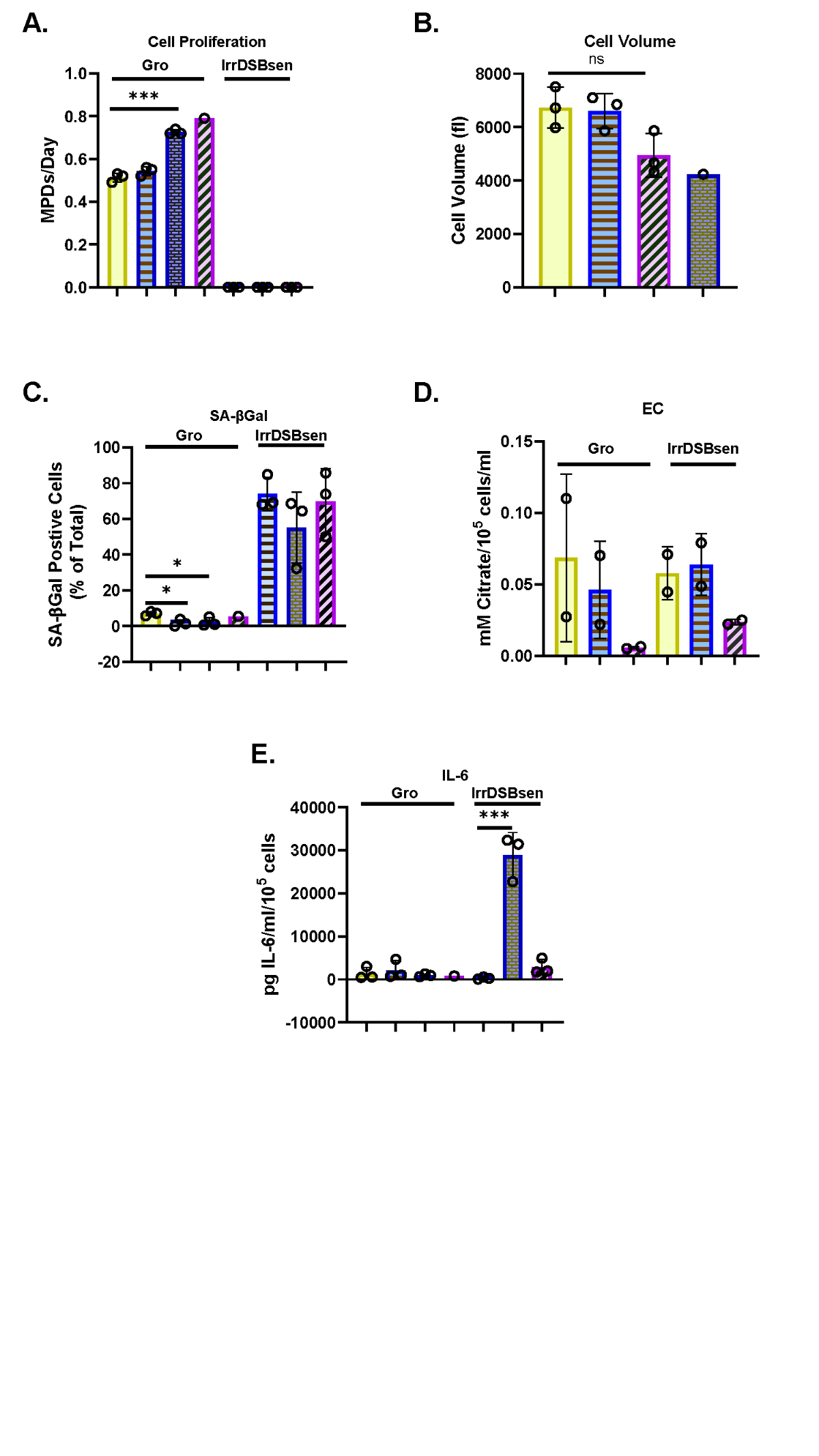

**Supplementary Figure 6** **The effect of the p38 mitogen-activated kinase and the Ataxia Telangiectasia Mutated (*ATM*) kinase on EC and secreted IL-6 in human fibroblasts.**

A. The effect of KU55933 and SB203580 individually and in concert on growing and IrrDSBsen BJ cell proliferation (N =3) except SB203580 alone (N =1).

B. The effect of KU55933 and SB203580 individually and in concert on growing and IrrDSBsen BJ cell volume (N =3) except SB203580 alone (N =1).

C. The effect of KU55933 and SB203580 individually and in concert on growing BJ SA-β Gal expression.

D. The effect of KU55933 and SB203580 individually and in concert on growing and IrrDSBsen BJ EC. (N =2).

E. The effect of KU55933 and SB203580 individually and in concert on growing and IrrDSBsen BJ IL-6. (N =3) except SB203580 alone (N =1).

Plain yellow bars = DMSO (vehicle) control; blue horizonal striped bars = 10 µM KU55933; dark blue brick pattern bars 10µM KU55933 and 10 µM SB203580; purple left-right diagonally striped bars = 10 µM SB203580.

* = P< 0.05; ** = P < 0.01; *** = P < 0.001; ns = not significant. The results are means +/- standard deviation. A. to G. N = 3. H. N=1.

* = P< 0.05; ** = P < 0.01; *** = P < 0.001; * P > 0.05 < 0.1; ns = not significant.

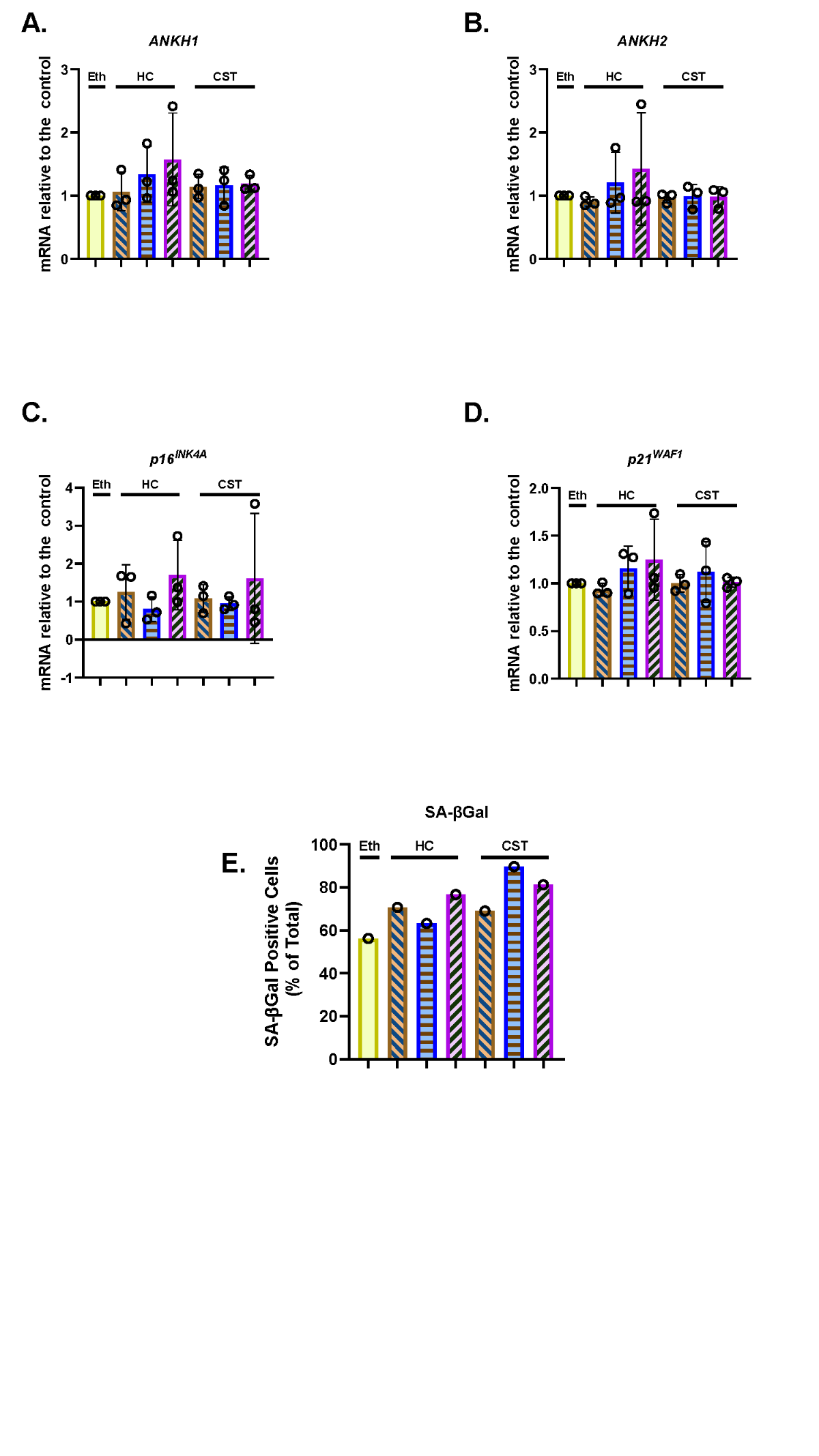
**Supplementary Figure 7 Steroids do not downregulate *ANKH* mRNA or senescence markers in IrrDSBsen NHOF-1 fibroblasts.**

A. The effect the indicated doses of HC and CST on the expression of *ANKH* mRNA using primer set 1.

B. As in A. but the results of primer set 2 in the same experiments

C. As in A. but the expression of the senescence marker p16^INK4A^ in the same experiments.

D. As in A. but the expression of the senescence marker p21^WAF1^ in the same experiments.

E. As in A. but the expression of senescence marker SA-βGal in the same experiments.

Plain yellow bars = control; brown right-left diagonally striped bars = 30 nM; blue horizontal striped bars = 100 nM; purple left-right diagonally striped bars = 300 nM.

* = P< 0.05; ** = P < 0.01; *** = P < 0.001; ns = not significant. The results are means +/- standard deviation. A. to G. N = 3. H. N=1.

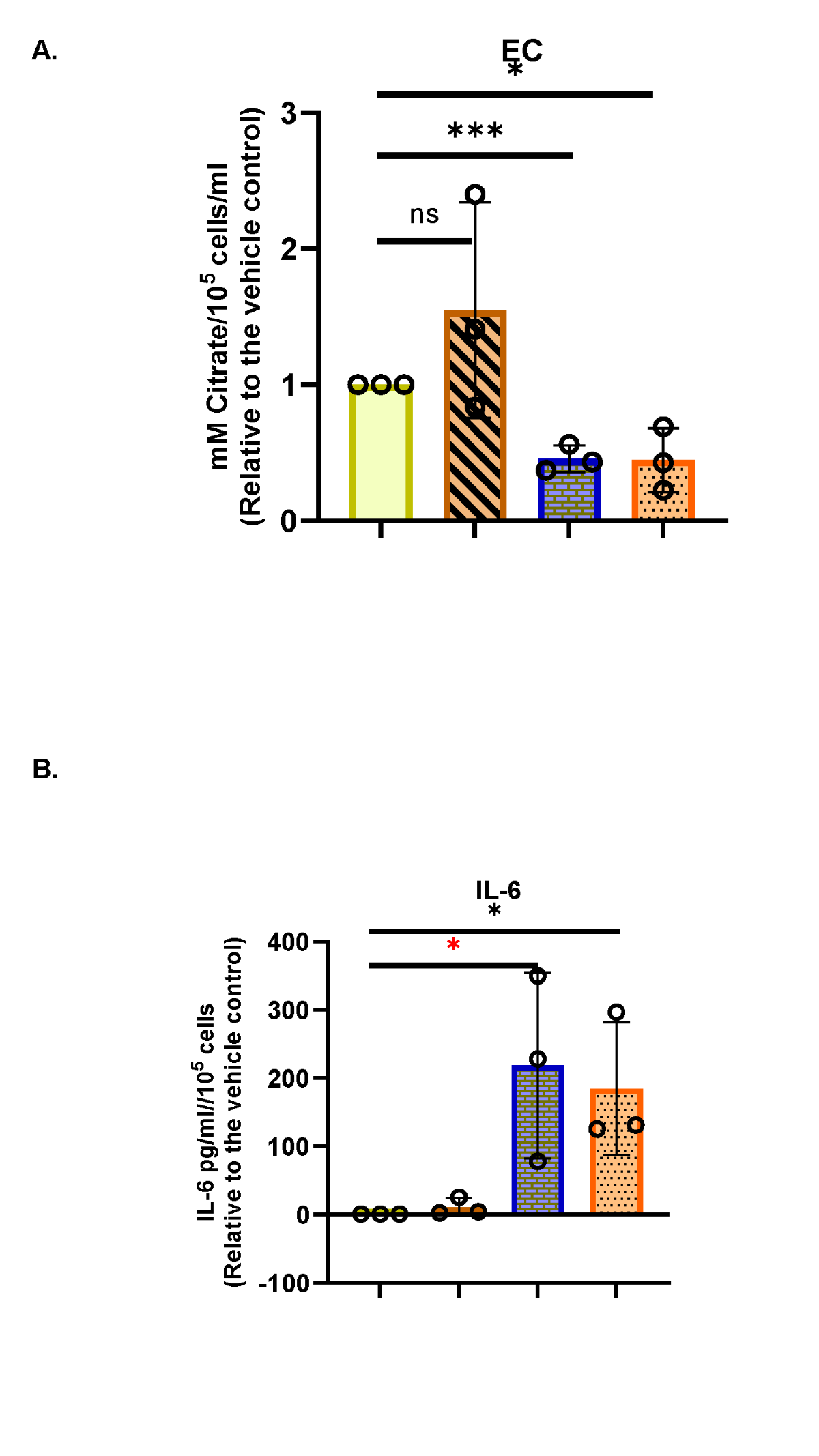
**Supplementary Figure 8 The effect of IL-1α and HC on human fibroblast levels of EC and IL-6.**

A. The effect of IL-1α and HC on EC.

B. The effect of IL-1α and HC on secreted IL-6.

Plain yellow bars = 0.1% vol/vol Ethanol (vehicle) control; brown right-left diagonally striped bars = 300 nM HC; dark blue brick pattern bars = 1 ng/mL IL-1α 1; orange stippled bars = IL-1α and 300 nM HC.

* = P< 0.05; ** = P < 0.01; *** = P < 0.001; ns = not significant. The results are means +/- standard deviation. N = 3.

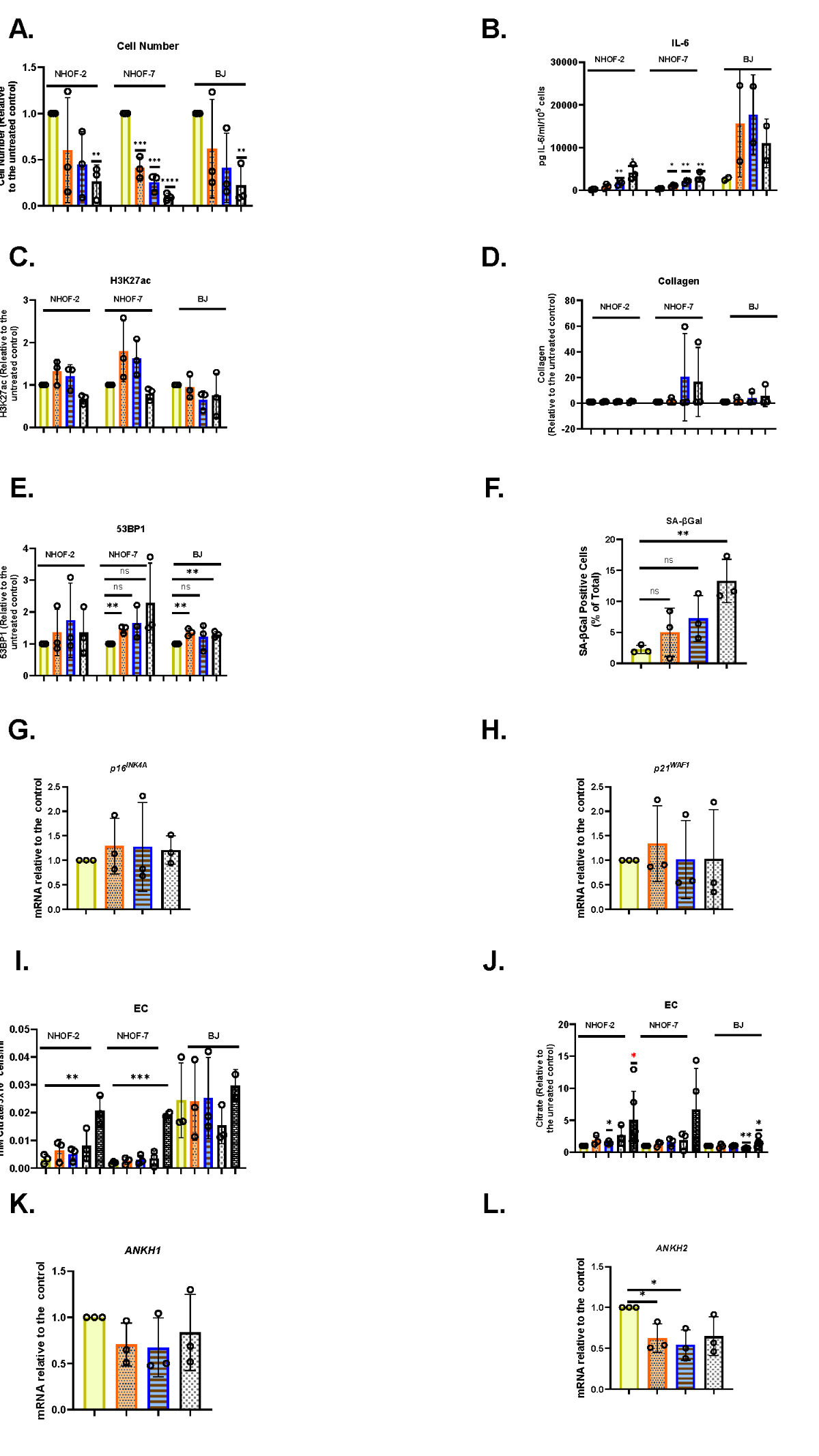
**Supplementary Figure 9 NaB induces proliferation arrest and IL-6 secretion but does not induce EC.**

A. The effect of 1-4 mM NaB on cell number in NHOF-2, NHOF-7 and BJ cells after treatment with 1-4 mM NaB for 4 days as assessed by Dapi fluorescence levels relative to the untreated controls.

B. As in A. but shows the level of IL-6 secretion in pg/mL.

C. As in A. but showing the levels of histone acetylation (H3K27ac) antibody fluorescence levels.

D. As in A. but showing collagen antibody fluorescence levels.

E. As in A. but showing 53BP1 antibody fluorescence levels.

F. Shows the % cells positive for SA-βGal in BJ cells following a four-day treatment with NaB 1-4 mM.

G. As in F. but showing the level of p16^INK4A^ transcript.

H. As in F. but showing the level of p21^WAF1^ transcript.

I. As in A. but showing the level of EC in mM/mL/10^5^ cells

J. As in A. but showing the level of EC relative to the untreated controls.

K. As in F. but shows the level ANKH transcript in BJ using primer set 1.

L. As in F. but showing the level *ANKH* mRNA expression using primer set 2.

Plain yellow bars = control; orange stippled bars = 1mM NaB; 2 mM NaB; blue horizonal striped bars = 2 mM NaB; white chequered bars = 4 mM NaB. F and G. Black chequered bars = IrrDSBsen controls.

* = P< 0.05; ** = P < 0.01; *** = P < 0.001; ns = not significant. The results are means +/- standard deviation. N = 3.

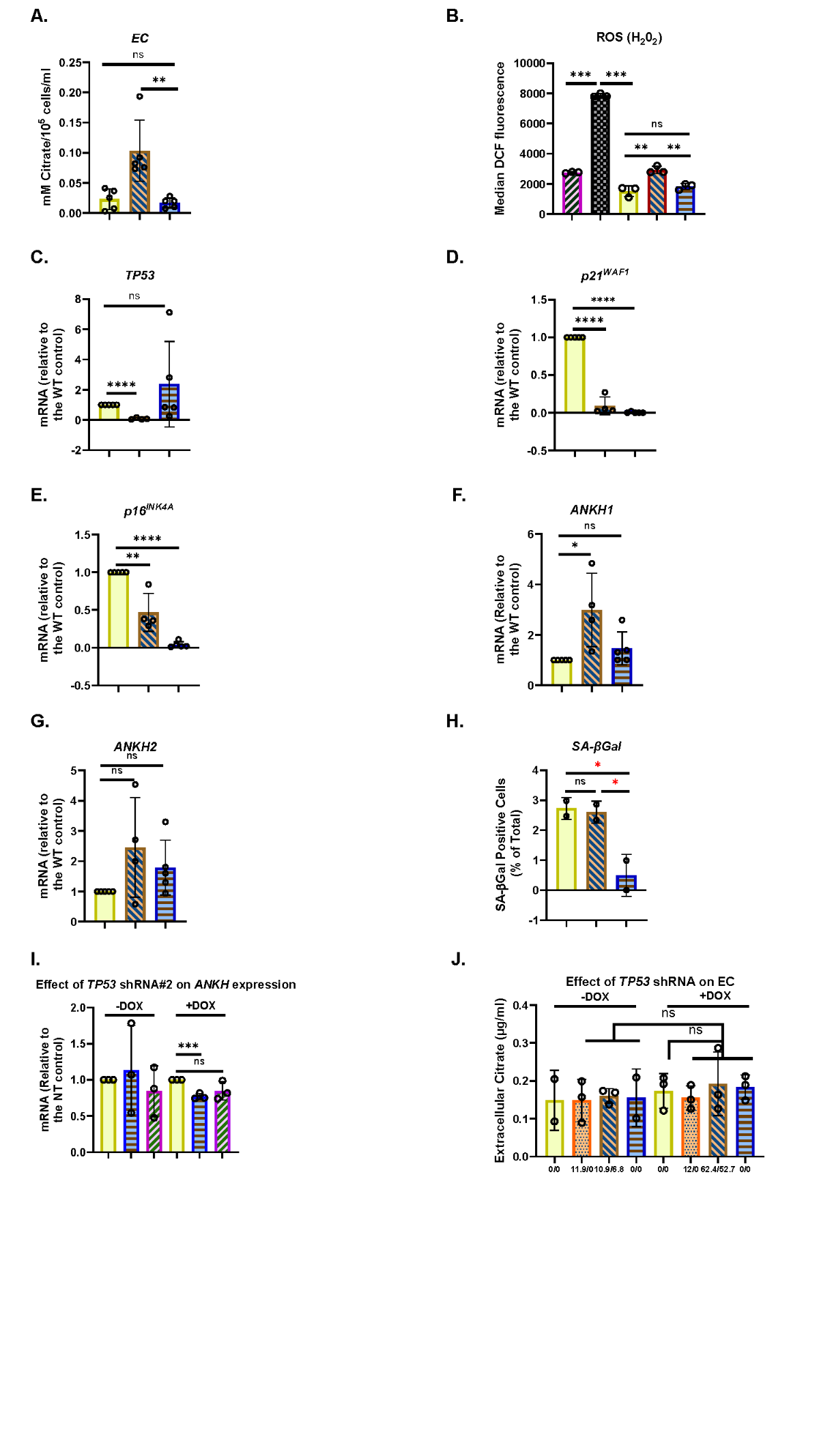

**Supplementary Figure 10 The effect of the senescence effectors on EC and *ANKH* expression in human fibroblasts.**

A. EC levels in wild type Loxo26 fibroblasts and their corresponding p53-/- and p21^WAF^-/- null lines. N = 5.

B. ROS (mainly H_2_O_2_) levels as assessed by DCF fluorescence and measured on the FACS in the same cell lines as in A. (N = 3).

C. TP53 transcript abundance of the cells in A. (N = 4).

D. p21^WAF1^ transcript abundance of the cells in A. N =4).

E. p16^INK4A^ transcript abundance of the cells in A. N =4).

F. *ANKH1* abundance of the cells in A as assessed by the primer set 1. N =4).

G. *ANKH1* abundance of the cells in A as assessed by the primer set 2. N =4).

H. The expression of senescence marker SA-βGal in the same experiments as in A. (N =2).

A.-H. Plain yellow wild type Loxo26; orange stippled bars p53-/- Loxo26; blue horizontal striped bars p21-/- Loxo26.

B. Grey brick bars = BJ cells; grey bars = Leiden p16^INK4A-^/- cells / p14^ARF^+/- cells.

I. The level of *ANKH* mRNA knockdown achieved by the *TP53* shRNAs relative to the Non-targeting (NT) control following doxycyclin (DOX) induction (*ANKH1*, blue horizontal striped bars), (*ANKH2*, Purple diagonally striped bars) or control (plain yellow bars) in growing NHOF-1 cells. N =3.

J. The EC level in growing NHOF-1 cells relative to the non-targeting (NT) control (plain yellow bars); *TP53* shRNA #1 (orange stippled bars); shRNA #2 (brown right-left diagonally striped bars); shRNA #3 (blue horizontal striped bars), following doxycyclin (DOX) induction (+DOX) or control (-DOX). N =3. Numbers under the bars indicate the % of *TP53*/p21^WAF^ knockdown, respectively.

* = P< 0.05; ** = P < 0.01; *** = P < 0.001; * P > 0.05 < 0.1; ns = not significant. The results are averages +/- standard deviation.

The experiments in A., B, C.- H and I.-J. were performed at separate times but using identical cell culture protocols and reagents.

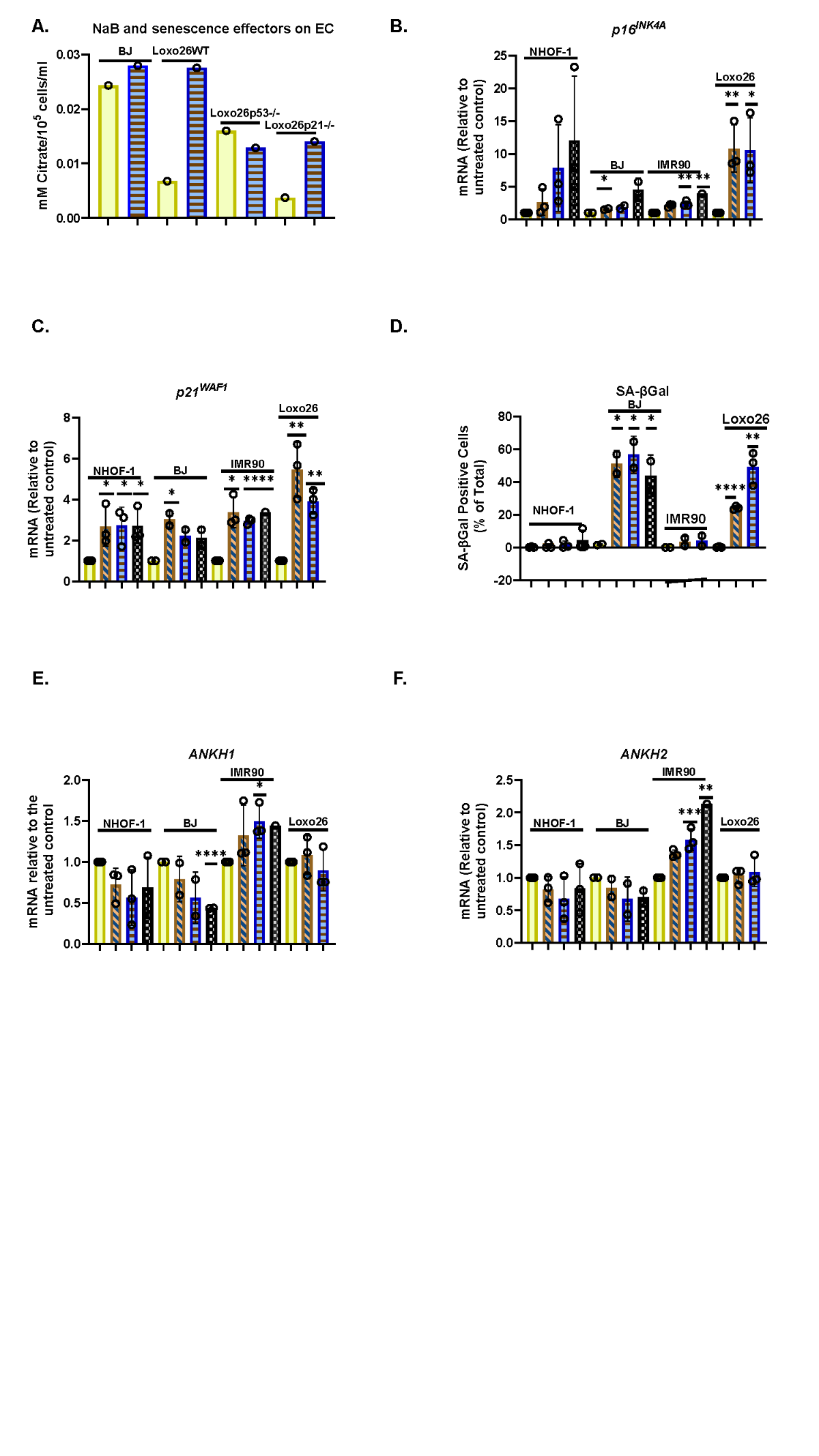

**Supplementary Figure 11 The effect of sodium butyrate (NaB) on cellular senescence, EC and the expression of *ANKH* in human fibroblasts**

A. A single experiment showing the effect of 0.5mM NaB on EC in BJ and Loxo26 human fibroblasts and the induction of EC only in Loxo26.

B. The effect of NaB at doses of 1-4mM for 17 days on NHOF-1, BJ, IMR90 and Loxo26 on the senescence marker p16^INK4A^in the same experiments.

C. As in B. but the expression of the senescence marker p21^WAF1^ in the same experiments.

D. As in B. but the expression of senescence marker SA-βGal in the same experiments

E. As in B. but the expression of *ANKH* mRNA using primer set 1.

F. As in B. but the results of primer set 2 in the same experiments.

A. Plain yellow bars = control; blue horizontal striped bars = 0.5mM NaB

B. – F. Plain yellow bars = control; brown right-left diagonally striped bars = 1 mM; blue horizontal striped bars = 2 mM; grey hatched bars = 4 mM NaB.

A. Plain yellow bars = control light blue horizontal striped bars = 0.5mM NaB * = P< 0.05; ** = P < 0.01; *** = P < 0.001; ns = not significant. The results are averages +/- standard deviation. N = 3. The experiments in A. and B-F. were performed at separate times but using identical protocols and reagents.

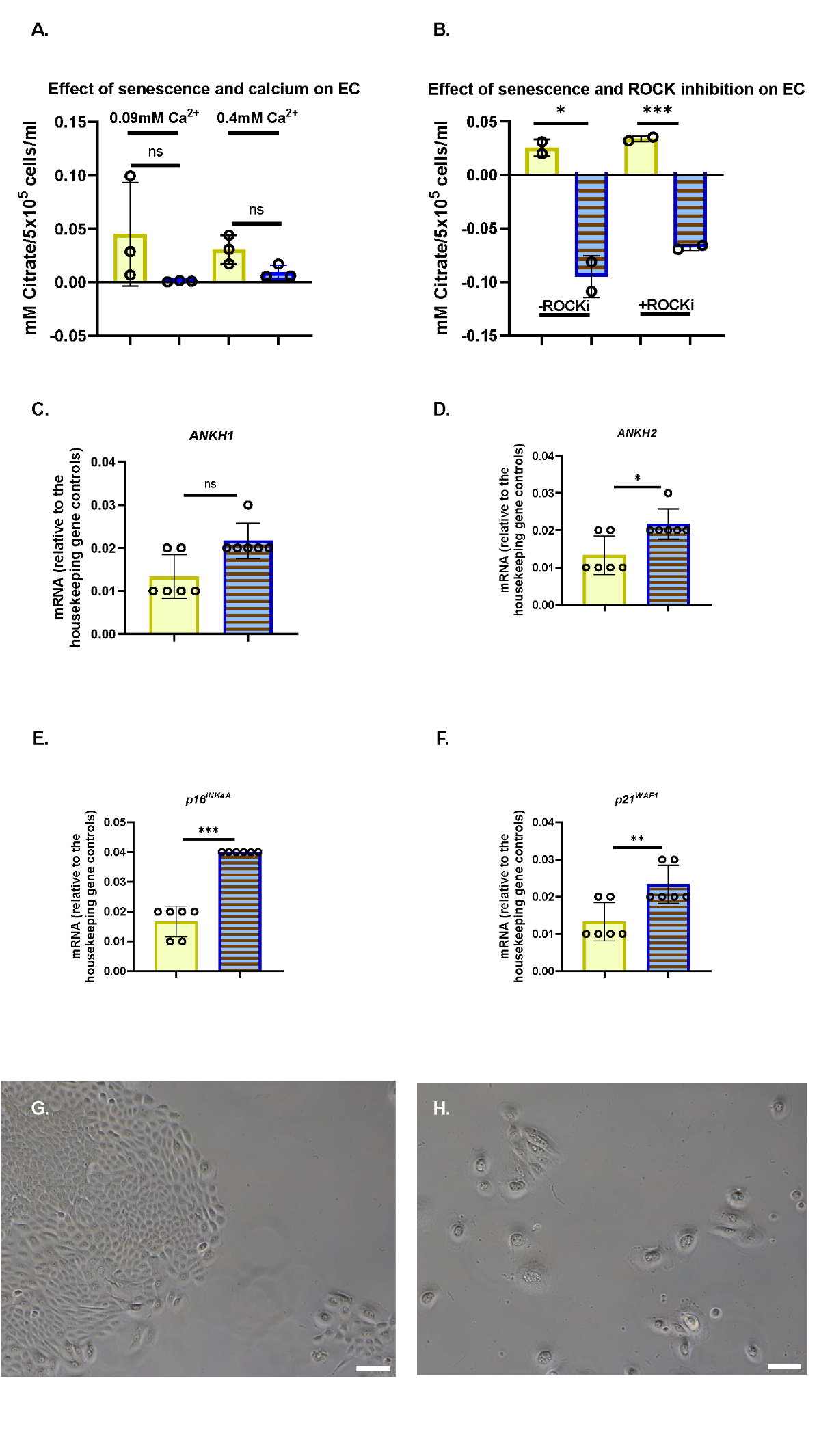
**Supplementary Figure 12 EC and *ANKH* are not upregulated following IrrDSBsen or PEsen in human epidermal keratinocytes**

A. EC in control (plain yellow bars) and IrrDSBsen keratinocytes (blue horizontal striped bars) cultured in serum-free medium (N=3)

B. The same lines as in A. cultured with irradiated 3T3 feeder cells with and without the ROCK inhibitor y27632. The 3T3 feeders were removed prior to the collection of the conditioned medium.

C. The expression of *ANKH* mRNA in PEsen HEK127 cells in 3 different Ca+ concentrations at 11.2 and 21.5 MPDs using primer set 1 (N = 6).

D. As in C. but the results of primer set 2 in the same experiments.

E. As in C. but the expression of the senescence marker p16^INK4A^in the same experiments.

F. As in C. but the expression of the senescence marker p21^WAF1^ in the same experiments.

A. Plain yellow bars = Control cells; blue horizontal striped bars = IrrDSBsen cells. 0.4mM Ca+ data stippled bars. B. As in A. but ROCKi treated cells stippled. C-F. Plain yellow bars = 11.2 MPDs; blue horizontal striped bars = 21.5 MPDs; * = P< 0.05; ** = P < 0.01; *** = P < 0.001; * P > 0.05 < 0.1; ns = not significant. The results are averages +/- standard deviation.

G. Image of young HEK127 (11.2 MPDs) in 0.4mM Ca+. H. Image of old HEK127 (21.5 MPDs) in 0.4mM Ca+. Bar = 100µM.

The experiments in A. and C.- G. were performed at separate times but using identical protocols and reagents. The data from C.-G. were derived from the same experiments as Supplementary Figure 13 C.-F.

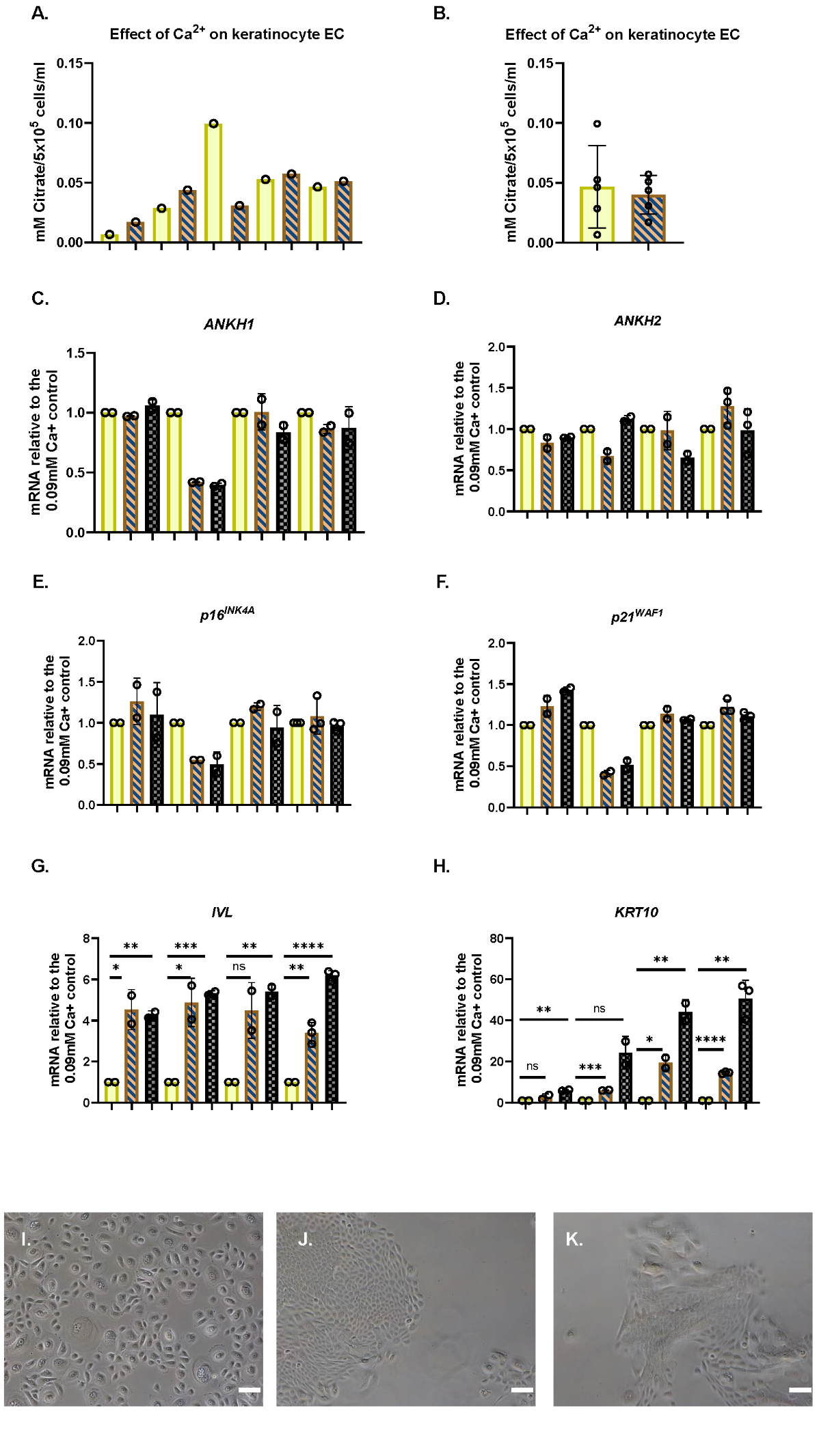

**Supplementary Figure 13** **EC and *ANKH* are not upregulated during keratinocyte differentiation.**

A. The effect of 0.4mM Ca+ on EC in five different epidermal keratinocyte lines in serum-free medium.

B. The average of the results in A. +/- standard deviation (n =5); ns = not significant.

C. The expression of *ANKH* mRNA in four epidermal lines in 3 different Ca+ concentrations

D. As in C. but the results of primer set 2 in the same experiments.

E. As in C. but the expression of the senescence marker p16^INK4A^in the same experiments.

F. As in C. but the expression of the senescence marker p21^WAF1^ in the same experiments.

G. As in C. but the expression of the differentiation marker INV in the same experiments.

H. As in C. but the expression of the terminal differentiation marker K10 in the same experiments.

Plain yellow bars = 0.09mM Ca+ cells; brown right-left diagonally striped bars = 0.4mM Ca+; dark grey hatched bars = 1.0mM Ca+. * = P< 0.05; ** = P < 0.01; *** = P < 0.001; * P > 0.05 < 0.1; ns = not significant. The results are averages +/- standard deviation (N =3).

I. Image of young HEK127 (11.2 MPDs) in 0.09mM Ca+ J. Image of the same cells as in I. in 0.4mM Ca+ (the same image as Supplementary Figure 12G). K. Image of the same cells as in I. in 1.0mM Ca+.

The experiments in A.-B. and C.- H. were performed at separate times but using identical protocols and reagents. The data from C.-H. were derived from the same experiments as Supplementary Figure 12 C.-G.

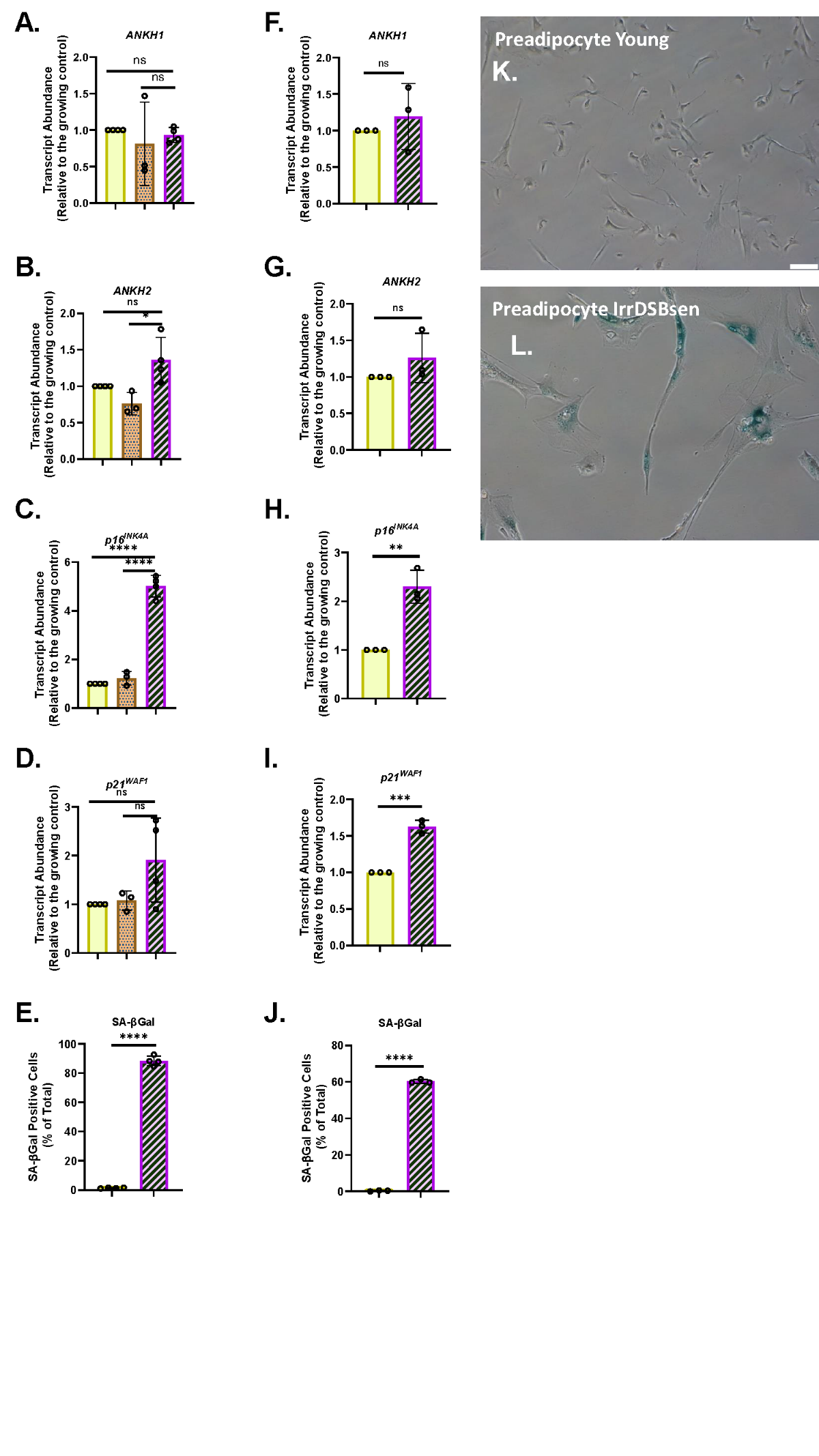
**Supplementary Figure 14 *ANKH* mRNA is not upregulated in IrrDSBsen and PEsen pre-adipocytes**

A.- E. IrrDSBsen; F-J. PEsen.

A. and F. The expression of *ANKH* mRNA using primer set 1.

B. and G The expression of *ANKH* mRNA using primer set 2.

C. and H. The expression of p16*^INK4A^* mRNA.

D. and I. The expression of p21^WAF1^ mRNA.

E. and J. SA-βGal expression.

Plain yellow bars A.-J. Young growing control. A-D brown stippled bars young confluent control; purple left-right diagonally striped bars A.-E. = IrrDSBsen ; F-J PEsen. * = P< 0.05; ** = P < 0.01; *** = P < 0.001; * trend for significance; ns = not significant. The results are means +/- standard deviation. N = 3.

K. Image of SA-βGal expression in the growing control from E. L. Image of SA-β Gal expression in the IrrDSBsen cells in E. Bar = 100 µm.

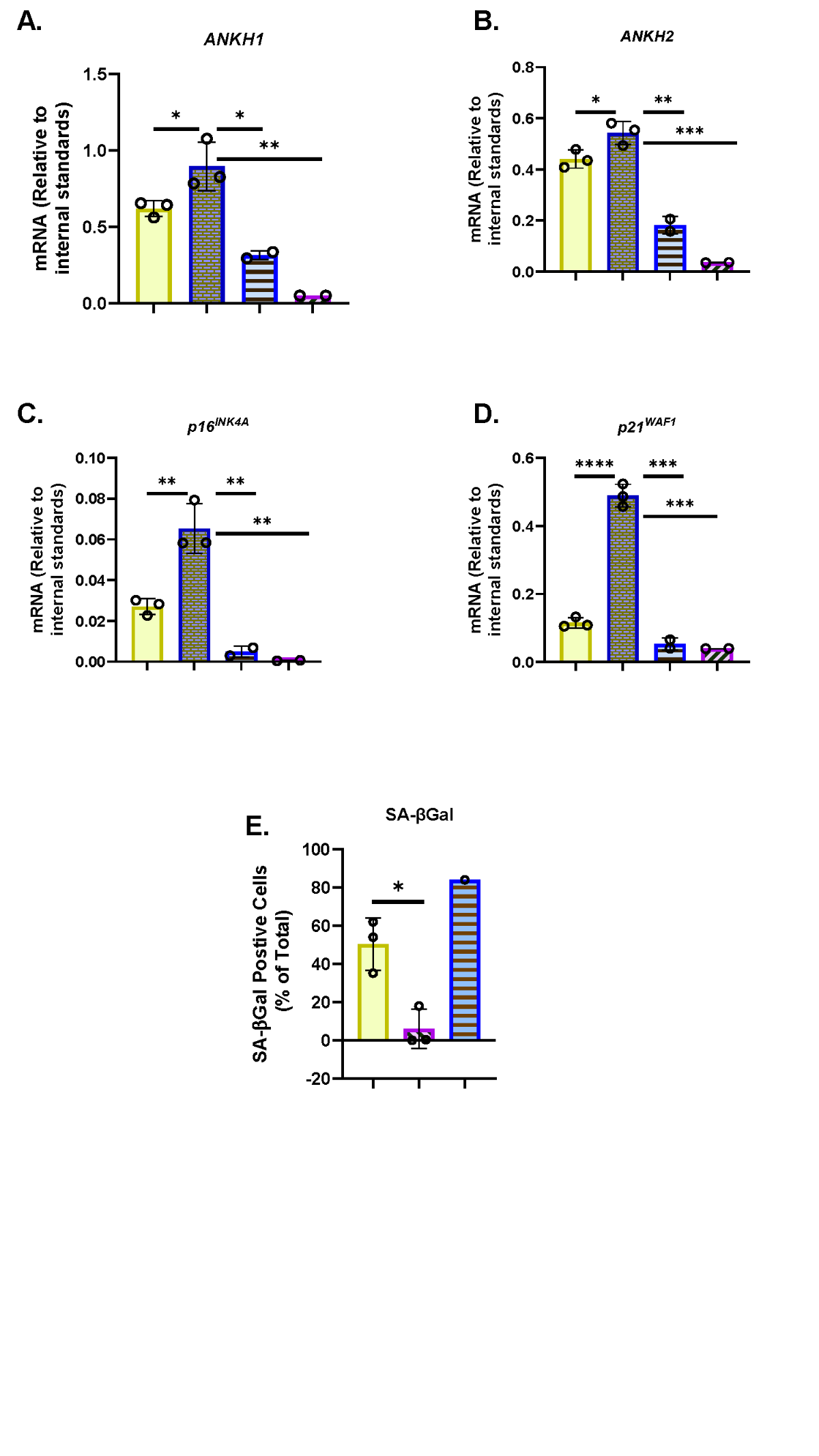
**Supplementary Figure 15 *ANKH* mRNA is upregulated in PEsen myoblasts and downregulated upon immortalisation.**

A. The expression of *ANKH* mRNA using primer set 1 in HMC TERT Clone T2 Young (Plain yellow bars) and old (dark blue brick pattern bars). (N = 3) and HMC TERT Pool Old (light blue horizontal striped bars) and HMC T15 (purple left-right diagonally striped bars). (N =2).

B. The expression of *ANKH* mRNA using primer set 2 in the same cells as A.

C. The expression of p16*^INK4A^* mRNA in the same cells as A.

D. The expression of p21^WAF1^ mRNA in the same cells as A.

E. SA-βGal expression in HMC TERT Pool (plain yellow bars) and HMC T15 (purple left-right diagonally striped bars) (N = 3) and senescent normal HMC cells (light blue horizonal bars (N =1).

* = P< 0.05; ** = P < 0.01; *** = P < 0.001; * P > 0.05 < 0.1); ns = not significant.

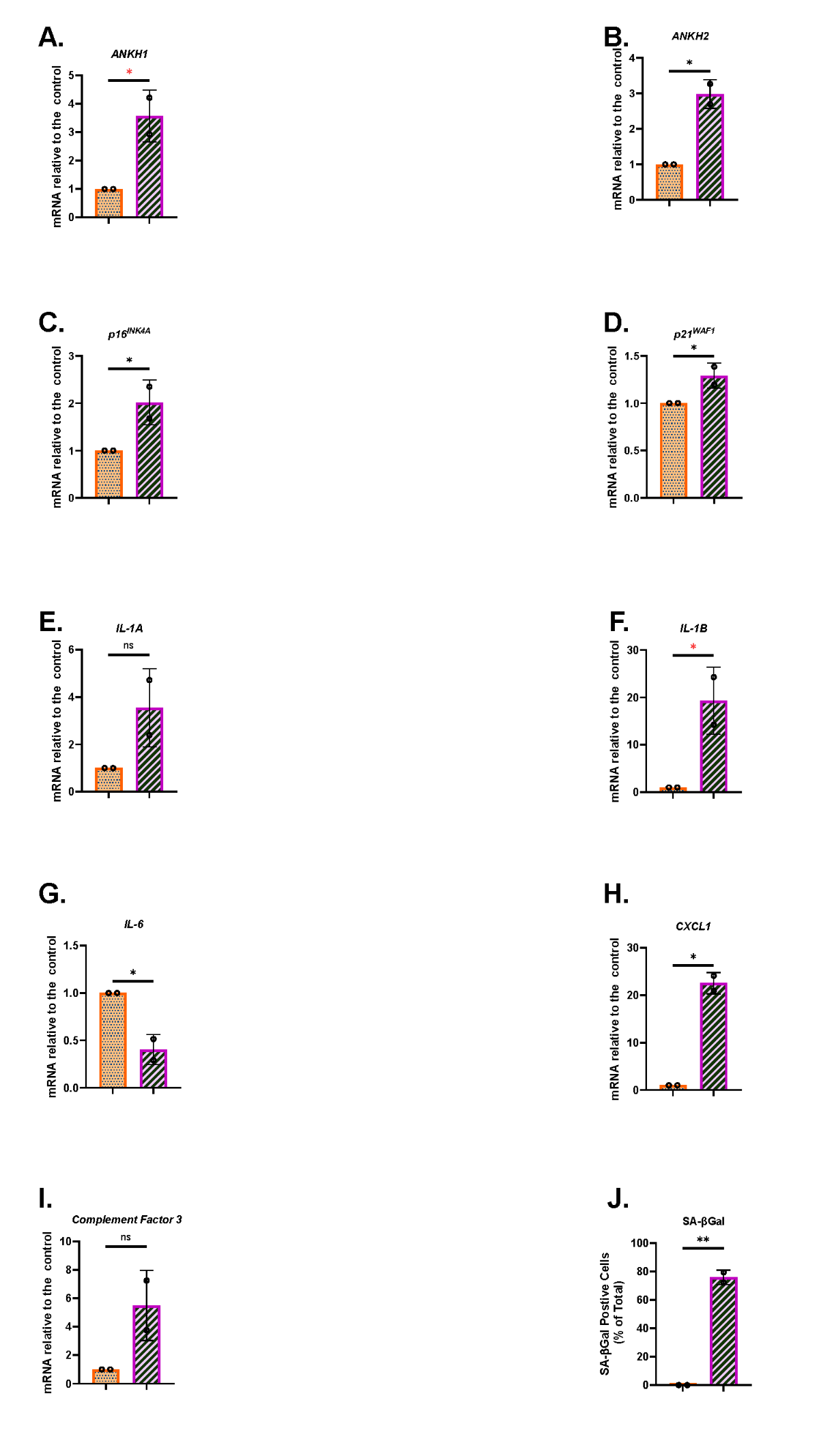
**Supplementary Figure 16 *ANKH* mRNA is upregulated in PEsen human astrocytes in parallel with SASP cytokines and markers of astrocyte activation.**

The figure represents a reanalysis of the experiments in Figure 12 (A-D. and J.) but this time including SASP cytokines and Complement Factor 3 (*C3* a marker of astrocyte activation), G.-I.

A. *ANKH* mRNA using primer set 1.

B. and G *ANKH* mRNA using primer set 2.

C. p16*^INK4A^* mRNA.

D. p21^WAF1^ mRNA.

E. *IL1A* mRNA.

F. *IL1B* mRNA.

G. *IL6* mRNA.

H. *CXCL1* mRNA.

I. *C3* mRNA.

J. SA-βGal expression.

Orange stippled bars = young growing astrocytes; purple left-right diagonally striped bars = PEsen astrocytes (28 MPDs). N =2.

* = P< 0.05; ** = P < 0.01; *** = P < 0.001; * P > 0.05 < 0.1; ns = not significant.
